# Re-education of myeloid immune cells to reduce regulatory T cell expansion and impede breast cancer progression

**DOI:** 10.1101/2023.08.14.553229

**Authors:** Hashni Epa Vidana Gamage, Sayyed Hamed Shahoei, Samuel T. Albright, Yu Wang, Amanda J. Smith, Rachel Farmer, Emma C. Fink, Elise Jacquin, Erin Weisser, Rafael O. Bautista, Madeline A. Henn, Claire P. Schane, Adam T. Nelczyk, Liqian Ma, Anasuya Das Gupta, Shruti V. Bendre, Tiffany Nguyen, Srishti Tiwari, Natalia Krawczynska, Sisi He, Evelyn Tjoanda, Hong Chen, Maria Sverdlov, Peter H. Gann, Romain Boidot, Frederique Vegran, Sean W. Fanning, Lionel Apetoh, Paul J. Hergenrother, Erik R. Nelson

## Abstract

Immune checkpoint blockade (ICB) has revolutionized cancer therapy but has had limited utility in several solid tumors such as breast cancer, a major cause of cancer-related mortality in women. Therefore, there is considerable interest in alternate strategies to promote an anti-cancer immune response. We demonstrate that NR0B2, a protein involved in cholesterol homeostasis, functions within myeloid immune cells to modulate the NLRP3 inflammasome and reduce the expansion of immune-suppressive regulatory T cells (T_reg_). Loss of NR0B2 increased mammary tumor growth and metastasis. Small molecule agonists, including one developed here, reduced T_reg_ expansion, reduced metastatic growth and improved the efficacy of ICB. This work identifies NR0B2 as a target to re-educate myeloid immune cells providing proof-of-principle that this cholesterol-homeostasis axis may have utility in enhancing ICB.

**Brief Summary:** Immune therapy has been disappointing for breast cancer. NR0B2 within myeloid immune cells reduces the expansion of T_regs_, a highly immune suppressive subtype historically challenging to target. NR0B2 within myeloid immune cells represses the inflammasome, leading to reduced T_reg_ expansion and subsequent tumor growth/metastasis. Activation of NR0B2 with small molecule agonists, including one developed herein, attenuates tumor growth and metastasis in murine models of mammary cancer.

## Introduction

In the last decade, immune therapy has revolutionized cancer treatment. Chimeric antigen receptor (CAR) T cells are remarkably effective, but are limited to tumors with known and unique antigens, or at least antigens on normal cells that are dispensable for life (ie: CD19 on B cells). In theory, immune checkpoint blockade (ICB) promised to apply to all solid tumors. However, in practice, it is only effective in certain tumor types or subtypes. Breast cancer is an example of this, where ICB has proven to be largely unsuccessful, and is currently only approved for those diagnosed with triple negative breast cancer (TNBC) whose tumors also stain positive for Programmed death-ligand 1 (PD-L1). Even amongst these patients receiving anti PD-1 (αPD1) in combination with nab-paclitaxel, only 20-30% show response with the majority being refractory^1,2^. Genentech recently announced that it will voluntarily withdraw atezolizumab (αPD-L1) in combination with nab-paclitaxel for consideration of accelerated approval for metastatic PD-L1-positive TNBC (August, 2021)^3^. Unfortunately, since TNBC lacks endocrine targets such as the estrogen receptor (ERα) or HER2, no targeted therapies exist other than for BRCA1/2 mutant tumors, resulting in a high recurrence and mortality rate. Therefore, there is urgent need to better understand alternate immune-suppressive networks in solid tumors such as TNBC, and develop therapies that combine ICB with pro-immune drugs that work in a checkpoint-independent manner.

Although undoubtedly multifactorial, one major obstacle to ICB is the highly immune-suppressive microenvironment of breast tumors – a phenomenon that is strongly maintained by myeloid immune cells, including macrophages, and regulatory T cells (T_regs_)^4,5^. Importantly however, myeloid cells are also critical for antigen presentation and a robust anti-tumor response. Therefore, it is now appreciated that rather than just eliminating or inhibiting myeloid cells, strategies are required to ‘re-educate’ myeloid cells away from being pro-tumorigenic and towards being anti-tumorigenic^6–8^.

Clinical observations have identified an association between hypercholesterolemia and breast cancer recurrence ^9^. On the other hand, large retrospective studies have now shown that patients taking cholesterol lowering medication (inhibitors of 2-hydroxy-3-methylglutaryl coenzyme A reductase; HMGCR; statins) demonstrate a significantly increased time to breast cancer recurrence reviewed by^10^. Recent studies found that statins were associated with improved breast cancer specific and overall survival in TNBC patients^11,12^. Collectively, these retrospective studies clearly highlight the direct clinical correlations between cholesterol and recurrence with metastatic disease. Our preclinical work has demonstrated that mice on a high cholesterol diet have increased tumor growth and metastatic burden^13,14^. Interestingly, it was found that a metabolite of cholesterol, 27-hydroxycholsterol (27HC), worked through myeloid cells to dramatically impair T cell expansion and function, and release extracellular vesicles that promote mammary cancer tumor growth and metastasis^14,15^. The effects of 27HC have been attributed to its ability to selectively modulate both the estrogen receptors (ERs) and liver x receptors (LXRs), major regulators of cholesterol homeostasis^13,15,16^.

Since myeloid cells appeared particularly susceptible to perturbations in cholesterol homeostasis, we explored whether other factors in this regulatory cascade might provide useful therapeutic targets, with a focus on nuclear receptors given that they are amenable to small molecule targeting^17^. Here, we identify a regulatory nuclear receptor, NR0B2 (small heterodimer partner) whose tumoral expression were associated with improved survival. NR0B2 within the liver is known to inhibit LXRs, but it also has unique attributes in myeloid cells resulting in T cell expansion away from T_regs_^18^. We show these myeloid-modulatory effects can be leveraged to promote a pro-immune and anti-cancer response.

## Results

### NR0B2 is expressed within breast tumors and is associated with increased recurrence free survival

When screening for proteins involved in cholesterol homeostasis that were also (1) potentially druggable and (2) implicated in breast cancer survival, we identified NR0B2. We initially surveyed mRNA expression of NR0B2 in different subtypes of breast cancer tumors, and found that it was expressed consistently lower in the Basal and Normal breast cancer subtypes compared to others (**SFig. 1A**). NR0B2 had slightly higher expression in ERα+ tumors compared to ERα-ones (**SFig. 1B**). mRNA expression of NR0B2 across different breast cancer cell lines was relatively low compared to a liver cancer line (**SFig. 1C**). Likewise, NR0B2 in murine mammary cancer lines was low compared to liver tissue, with myeloid cell lines having slightly higher expression (**SFig1C**).

**Figure 1:**
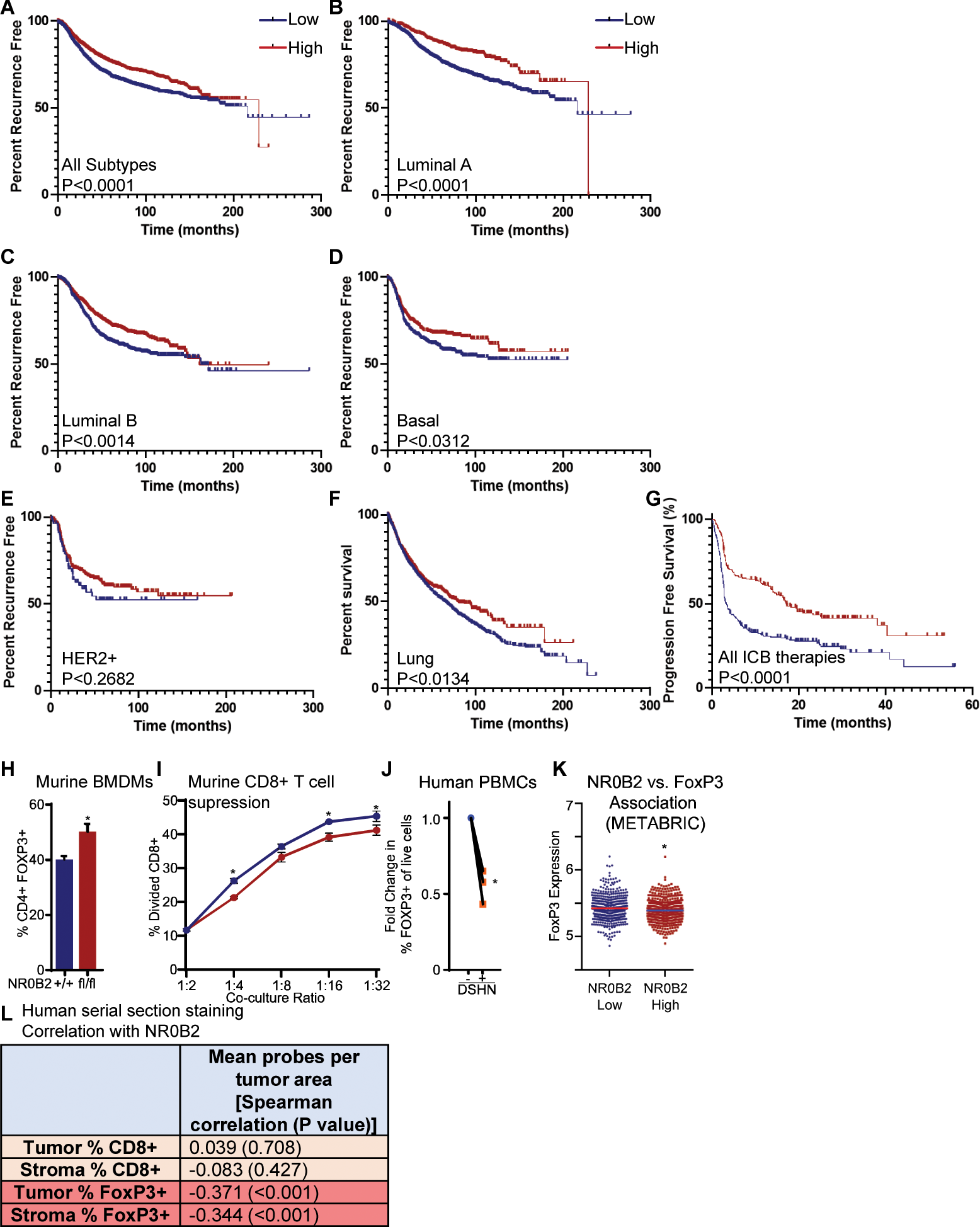
NR0B2 mRNA expression within breast and lung tumors is associated with an increased recurrence free survival time. **(A)** All breast cancer subtypes are considered. **(B)** Luminal A only. **(C)** Luminal B only. **(D)** Basal only. **(E)** HER2+ only. **(F)** Lung cancer Kaplan Meier plotter, autocutoff. **(G)** NR0B2 mRNA expression within tumors from patients treated with immune checkpoint blockers (ICB) is associated with an increased progression free survival time. The Kaplan-Meier plotter was used to probe associations between NR0B2 expression in both genders and all cancers within the database. Complementary data in **SFig. 1. (H)** Loss of NR0B2 in murine bone marrow derived macrophages (BMDMs) results in decreased T_reg_ expansion in T cells activated with PMA and ionomycin (N=3/group, t test). **(I)** CD4+ T cells expanded in co-culture with BMDMs lacking NR0B2 (NR0B2^fl/fl^;LysMcre) suppressed CD8+ T cell expansion compared to those co-cultured with control (NR0B2^+/+^;LysMcre) macrophages. Expanded CD4+ T cells were cocultured at different ratios with splenic CD8+ T cells that had been activated with CD3/28 Dynabeads and subsequent proliferation of CD8+ cells was assessed. **(J)** DCs derived from PBMCs of healthy human volunteers resulted in decreased T_reg_ expansion in T cells when treated with DSHN, a NR0B2 agonist (N=3/group). **(K)** FoxP3 expression is lower in human tumors with high NR0B2 expression (upper quartile compared to lower quartile depicted here, correlation regression analysis in **SFig. 5**, N=477-478/group, t test). Data obtained from METABRIC. **(L)** There is an inverse correlation between NR0B2 and FoxP3 in a French cohort of breast cancer samples. Breast tumors were serially sectioned and stained with NR0B2 (*in situ* hybridization), FoxP3 (IHC) or CD8 (IHC). Sections were counterstained with cytokeratin to differentiate between tumoral and stromal regions. Representative counterstained section above table indicating Spearman correlation and P value (N=93). * indicates P<0.05 by indicated statistical test. Extended analysis in **SFig. 6**.

Despite its lower expression in breast cancer cells compared to liver, elevated NR0B2 expression within human breast tumors was associated with increased recurrence free survival (**Fig. 1A-E**). This was apparent when all breast cancer subtypes were considered, or when Luminal A, B, or Basal were considered independently. The HER2 subtype was underpowered to draw firm conclusions. We also found that lung cancer tumors with increased NR0B2 were associated with improved survival (**Fig. 1F**), an important finding given the prevalence of lung cancer, and that the lung is a clinically important metastatic site for breast cancer. Several other cancers that were assessed also showed an association between elevated NR0B2 expression and overall survival, providing a clue that commonalities within the host microenvironment may be responsible for any potential effects of NR0B2 (significant associations found in bladder cancer, renal clear cell carcinoma and lung adenocarcinoma; **SFig. 2A**). Even more strikingly, there is a strong association between NR0B2 expression and both increased overall and progression free survival in patients treated with ICB, a trend that holds when only those patients treated with either αPD-1 or αPD-L1 are considered, with a non-significant effect for αCTLA-4 (P=0.17 for overall survival) (pan-cancer analysis, **Fig. 1G** and **SFig. 2B-C**).

**Figure 2:**
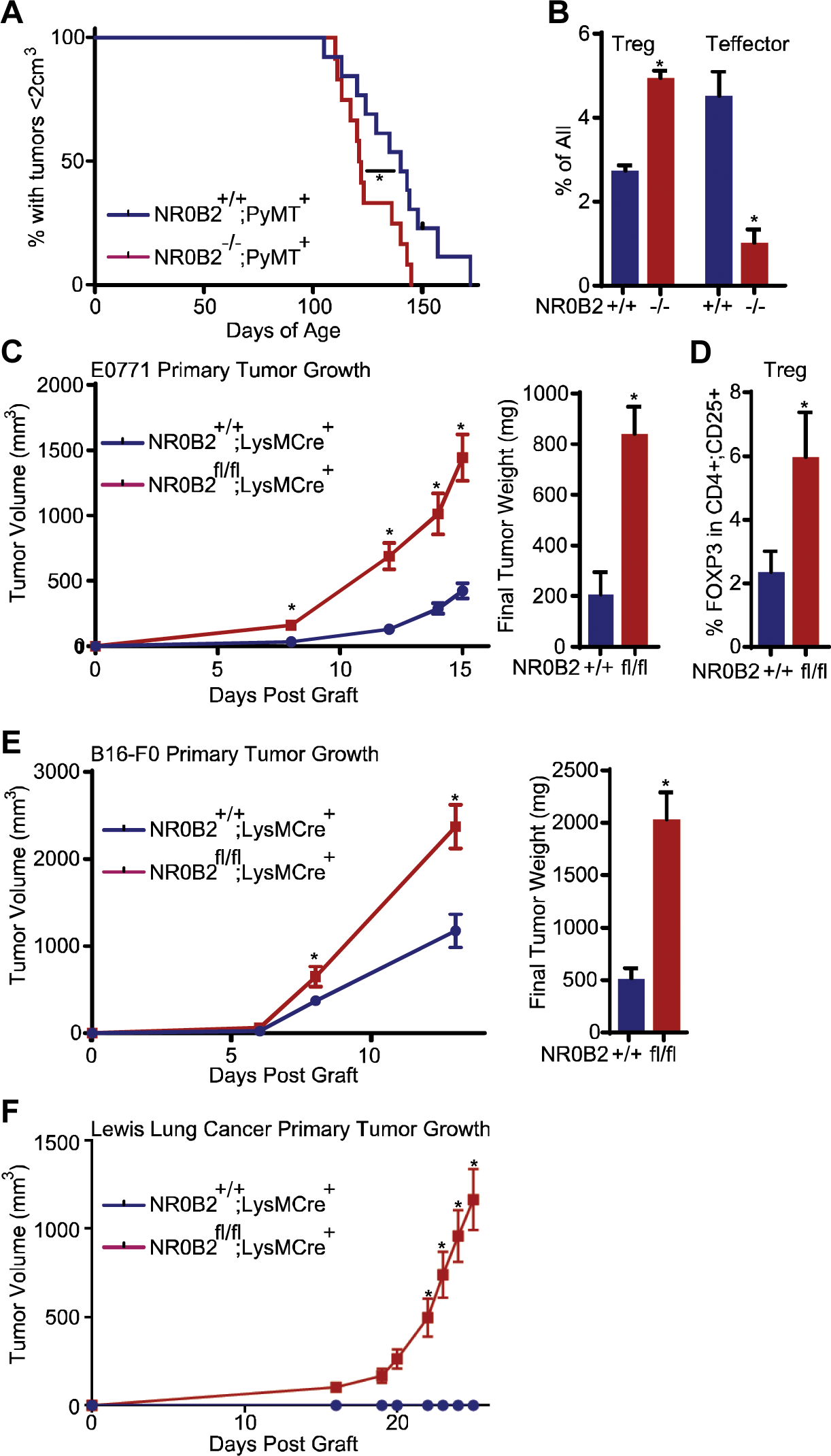
Loss of NR0B2 results in increased Tregs and tumor growth in murine models. **(A)** Germline deletion of NR0B2 promotes growth of spontaneous mammary tumors in MMTV-PyMT mice. Time from birth to developing a tumor burden of >2000mm^3^ is depicted. NR0B2^-/-^;PyMT^+^ and NR0B2^+/+^;PyMT^+^ littermates were monitored frequently after birth for development of mammary masses and subsequent tumor growth was assessed by caliper measurements (N=12 per group, Log-rank test). **(B)** MMTV-PyMT tumors from NR0B2^-/-^ mice had increased T_regs_ (CD4^+^;CD25^+^;FoxP3^+^) and decreased T_eff_ (CD4^+^;IFN-γ^+^) cells compared to wildtype controls (N=3-6/group, t test). **(C)** Orthotopic, syngeneic E0771 mammary tumors grew at a faster rate in mice with NR0B2 selectively knocked out in myeloid cells compared to control mice (NR0B2^fl/fl^;LysMCre^+^ vs. NR0B2^+/+^;LysMCre^+^). Tumor growth curve to the left of the final tumor weight at necropsy (N=10-15/group, 2-way ANOVA followed by Šidák’s multiple comparison, or t test). **(D)** In a separate experiment, tumors isolated from NR0B2^fl/fl^;LysMCre^+^ mice had significantly more T_reg_ cells compared to NR0B2^+/+^;LysMCre^+^ control mice (N=10-11/group, t test). **(E)** Murine melanoma B16-F0 tumors grew at a faster rate in NR0B2^fl/fl^;LysMCre^+^ compared to NR0B2^+/+^;LysMCre^+^ control mice. Tumor growth curve to the left of the final tumor weight at necropsy (N=12-16/group, 2-way ANOVA followed by Šidák’s multiple comparison, or t test). **(F)** Murine Lewis Lung tumors grafted at suboptimal cell numbers grew at a faster rate NR0B2^fl/fl^;LysMCre^+^ compared to NR0B2^+/+^;LysMCre^+^ control mice (N=11-14/group, 2-way ANOVA followed by Šidák’s multiple comparison). * indicates P<0.05 using indicated statistical test.

### Protective attributes of NR0B2 are likely due to its expression in myeloid cells

Neither overexpression nor siRNA mediated knockdown of NR0B2 had a significant effect on proliferation on various breast cancer cell lines, despite eliciting expected changes in target gene expression (**SFig. 3A-B**). Likewise, treatment with the NR0B2 agonist DSHN^19^ had no effect on proliferation (**SFig. 3C**). These approaches to modulate NR0B2 activity also failed to impact cellular migration (**SFig. 3D**). This would suggest that the protective effects of NR0B2 observed in **Fig. 1** were mediated by cells within the microenvironment, not the cancer cells themselves. scRNA-seq analysis of normal breast tissue indicates that the only cell types that express NR0B2 to appreciable levels are breast glandular cells and macrophages, although fibroblasts and breast myoepithelial cells also had some expression (data from The Human Protein Atlas, **SFig. 4A**). Among human PBMCs, various subtypes of dendritic cells (DCs) and monocyte populations had elevated expression of NR0B2 (**SFig. 4B**). Our findings that NR0B2 did not impact breast cancer cell proliferation, coupled with previous data indicating that bone marrow and myeloid cells have high expression of NR0B2 compared to other tissues^18^, suggest that the favorable prognostic association with NR0B2 in breast tumors (**Fig. 1A-G**) was potentially mediated through myeloid immune cells.

**Figure 3:**
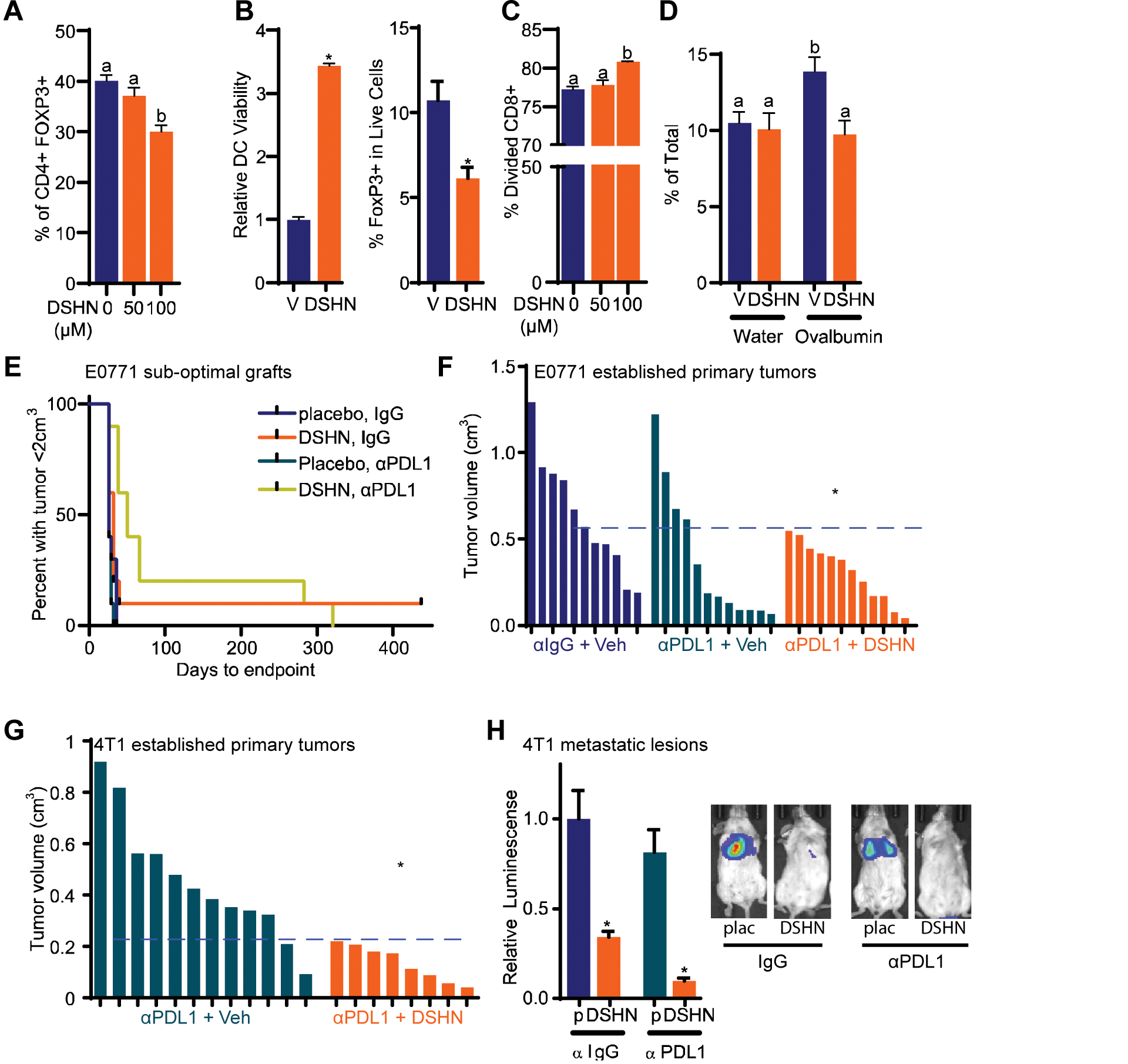
DSHN, a tool small molecule agonist of NR0B2 reduces macrophage/dendritic cell induced T_reg_ expansion and tumor growth in murine models. **(A)** Co-culture of DSHN pre-treated BMDMs results in decreased T_reg_ expansion in a dose dependent manner. **(B)** Dendritic cells (left - CD11B^+^;CD11C^+^) and CD4^+^ T cells (right) were isolated from MMTV-PyMT tumors by FACS, and treated with vehicle or DSHN for 3d. Viability for extracted DCs was determined (left graph), as well as the resulting frequency of T_reg_ (CD4^+^;CD25^+^;FoxP3^+^) was quantified by flow cytometry (right graph), (N=3-5/group, t test). **(C)** T cells expanded in the presence of DSHN treated BMDMs were less suppressive than those expanded in the presence of vehicle treated BMDMs, in terms of a subsequent co-culture of naïve T cells. The percentage of divided CD8+ T cells in the second round was quantified. **(D)** Long term treatment with DSHN reduced T_regs_ (CD4+,FoxP3+) in the spleen of mice provided a chronic exposure to an immunogenic protein (ovalbumin) in their water (N=5/group, 1-way ANOVA followed by Šidák test). **(E)** The effects of DSHN on subsequent tumor outgrowth of a sub-optimal E0771 orthotopic graft. C57BL/6 Mice were orthotopically grafted with a low number of E0771-luc mammary cancer cells. After allowing one day for tumor establishment, daily treatment with placebo or DSHN commenced. Immune checkpoint inhibitor treatment (αPDL1) was initiated 3 days post-graft, and continued every 2 days for six total treatments. 18d postgraft, all treatments were stopped and mice were monitored for subsequent tumor out-growth. Time from graft to developing a tumor burden of >2000mm^3^ is depicted. Black dash represents censored mouse that had no palpable tumor 437d post graft (N=10/group, Log-rank test). **(F)** DSHN increases efficacy of αPDL1 therapy on orthotopic E0771 tumors. An optimal number of E0771 cells were grafted into FoxP3-EGFP mice and allowed to grow until tumors reached 100mm^3^. Mice were euthanized on day 19 (waterfall plot, Fisher’s exact test). **(G)** DSHN increases efficacy of αPDL1 therapy on orthotopic 4T1 tumors. 4T1 cells were orthotopically grafted into mice and allowed to establish for 5d prior to treatment start. Mice were euthanized on day 16 (waterfall plot, Fisher’s exact test). **(H)** DSHN reduces metastatic outgrowth of 4T1 tumors and enhances efficacy of αPDL1 compared to placebo (p) (2-way ANOVA followed by Šidák’s multiple comparison test for day 16). 4T1 cells were grafted intravenously, and lung metastatic lesions allowed to establish for 3d before treatment commenced. At the end of the study, lungs were removed and imaged *ex vivo*, quantified data indicated (N=10, 1-way ANOVA followed by Newman-Keuls multiple comparison test). Representative luciferase images shown to the right. * or different letters denote P<0.05 using indicated statistical test. C57BL/6 Mice were orthotopically grafted with a low number of E0771-luc mammary cancer cells. After allowing one day for tumor establishment, daily treatment with placebo or DSHN commenced. Immune checkpoint inhibitor treatment (αPDL1) was initiated 3 days post-graft, and continued every 2 days for six total treatments. 18d postgraft, all treatments were stopped and mice were monitored for subsequent tumor out-growth. Time from graft to developing a tumor burden of >2000mm^3^ is depicted. Black dash represents censored mouse that had no palpable tumor 437d post graft (N=10/group, Log-rank test). **(F)** DSHN increases efficacy of αPDL1 therapy on orthotopic E0771 tumors. An optimal number of E0771 cells were grafted into FoxP3-EGFP mice and allowed to grow until tumors reached 100mm^3^. Mice were euthanized on day 19 (waterfall plot, Fisher’s exact test). **(G)** DSHN increases efficacy of αPDL1 therapy on orthotopic 4T1 tumors. 4T1 cells were orthotopically grafted into mice and allowed to establish for 5d prior to treatment start. Mice were euthanized on day 16 (waterfall plot, Fisher’s exact test). **(H)** DSHN reduces metastatic outgrowth of 4T1 tumors and enhances efficacy of αPDL1 compared to placebo (p) (2-way ANOVA followed by Šidák’s multiple comparison test for day 16). 4T1 cells were grafted intravenously, and lung metastatic lesions allowed to establish for 3d before treatment commenced. At the end of the study, lungs were removed and imaged *ex vivo*, quantified data indicated (N=10, 1-way ANOVA followed by Newman-Keuls multiple comparison test). Representative luciferase images shown to the right. * or different letters denote P<0.05 using indicated statistical test.

**Figure 4:**
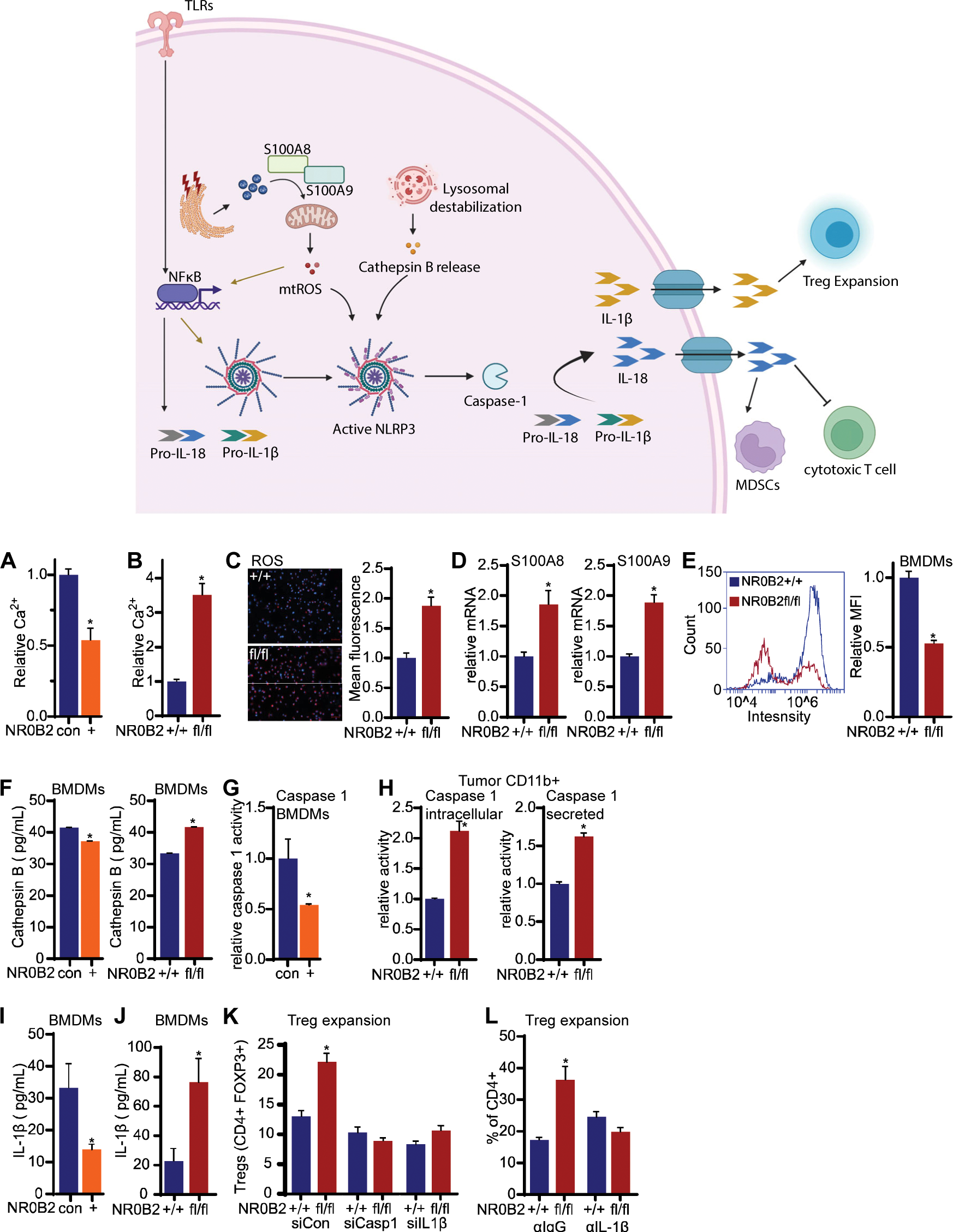
NR0B2 regulates many aspects of the inflammasome resulting in altered T_reg_ expansion. An overview cartoon of the relevant aspects of the inflammasome is depicted above the data. (**A**) Ca^2+^ concentration decreases in ionomycin-treated BMDMs overexpressing NR0B2 (N=3/group, t test), and (**B**) increased when NR0B2 is knocked out. (**C**) Reactive oxygen species (ROS) is increased in BMDMs where NR0B2 is knocked out. Representative images to the left of quantified data (N=3/group, t test). (**D**) S100A8 and S100A9 transcript levels are increased in CD11B+ cells isolated from tumors where NR0B2 was knocked out of the myeloid cell lineage (N=4/group, t test). (**E**) Lysosome stability is decreased in BMDMs lacking NR0B2. Representative flow cytometry histogram to the left of quantified mean fluorescence intensity (MFI, N=4/group). (**F**) Left panel: secreted cathepsin from control BMDMs or those overexpressing NR0B2 (+). Right panel: secreted cathepsin from control BMDMs (+/+) or those lacking NR0B2 (fl/fl). (N=4/group, t test) as indicated. (**G**) Intracellular caspase 1 activity in BMDMs transfected with control or NR0B2 expression plasmid (N=5/group, t test). (**H**) Intracellular and secreted caspase 1 activity in CD11B+ cells isolated from E0771 tumors. (**I&J**) IL-1β secreted protein from BMDMs (N=4/group, t test). **(K)** siRNA against caspase 1 or IL-1β attenuates induction of T_regs_ in co-culture with myeloid cells from NR0B1 ^fl/fl^;LysMcre^+^ mice (N=5/group) **(L)** Immune neutralization with αIL-1β attenuates induction of T_regs_ in co-culture with myeloid cells from NR0B1 ^fl/fl^;LysMcre^+^ mice.

When evaluating different myeloid populations, we observed that NR0B2 expression was downregulated in cells associated with an immune-suppressive phenotype such as M2 polarized macrophages^18^, and myeloid derived suppressor cells (**SFig 4C**). Furthermore, dendritic cells isolated from tumor bearing mice also had decreased NR0B2 (**SFig 4D**). Collectively, these observations suggest that NR0B2 within myeloid cells may exert anti-tumoral properties and is downregulated as part of the immune-suppressive program found in tumors. Importantly in this regard, we have previously shown that NR0B2 within macrophages was able to skew T cell expansion away from the highly immune-suppressive, regulatory T cell type (T_reg_)^18^. Tumoral infiltration of T_regs_ is a poor prognostic and implicated in resistance to most therapies^20–22^. We confirmed that genetic loss of NR0B2 in murine bone marrow derived macrophages (BMDMs, from NR0B2^fl/fl^;LysMcre^+^ mice) resulted in increased T_reg_ expansion **(Fig. 1H**). Furthermore, T cells expanded in co-culture with NR0B2^fl/fl^;LysMcre^+^ BMDMs inhibited the normal expansion of CD8+ T cells, demonstrating their suppressive nature (**Fig. 1I**).

To demonstrate that this axis is conserved in humans, we made use of cells from healthy volunteers and found that treatment of PBMC-derived-DCs with a small molecule agonist of NR0B2, DSHN, resulted in decreased T_reg_ expansion (**Fig. 1J**). To gain further insight into the relevance of these findings in human disease, we assessed the METABRIC dataset and found an inverse correlation between NR0B2 and FoxP3 expression, FoxP3 being a marker of T_regs_ (**Fig. 1K, SFig. 5**). We then obtained serial sections from 93 human breast tumors and stained for NR0B2, FoxP3 and CD8. Extensive optimization failed to find an antibody specific or suitable for protein staining of NR0B2, so we instead used an *in situ* hybridization approach for this target. Sections were counter-stained with cytokeratin allowing us to evaluate differences between tumor nests and stromal regions. CD8 cells were more abundant in the tumor stroma of HER2+ cases, but at similar concentrations within the tumor itself compared to other subtypes (**SFig. 6A**). FoxP3 cells were lower in ERα+/PR+ cases compared to HER2+ or TNBC cases, regardless of tumor or stroma (**SFig. 6B**). NR0B2 within the tumor or stroma was significantly more prevalent in ERα+/PR+ cases compared to HER2+ or TNBC cases (**SFig. 6C**). This is in congruence with the METABRIC data indicating that ERα^+^ tumors had elevated NR0B2 expression compared to ERα^-^ ones (**SFig. 1A**). In general, CD8 and FoxP3 expression were correlated within the tumoral and stromal areas (**SFig. 6D**).

**Figure 5:**
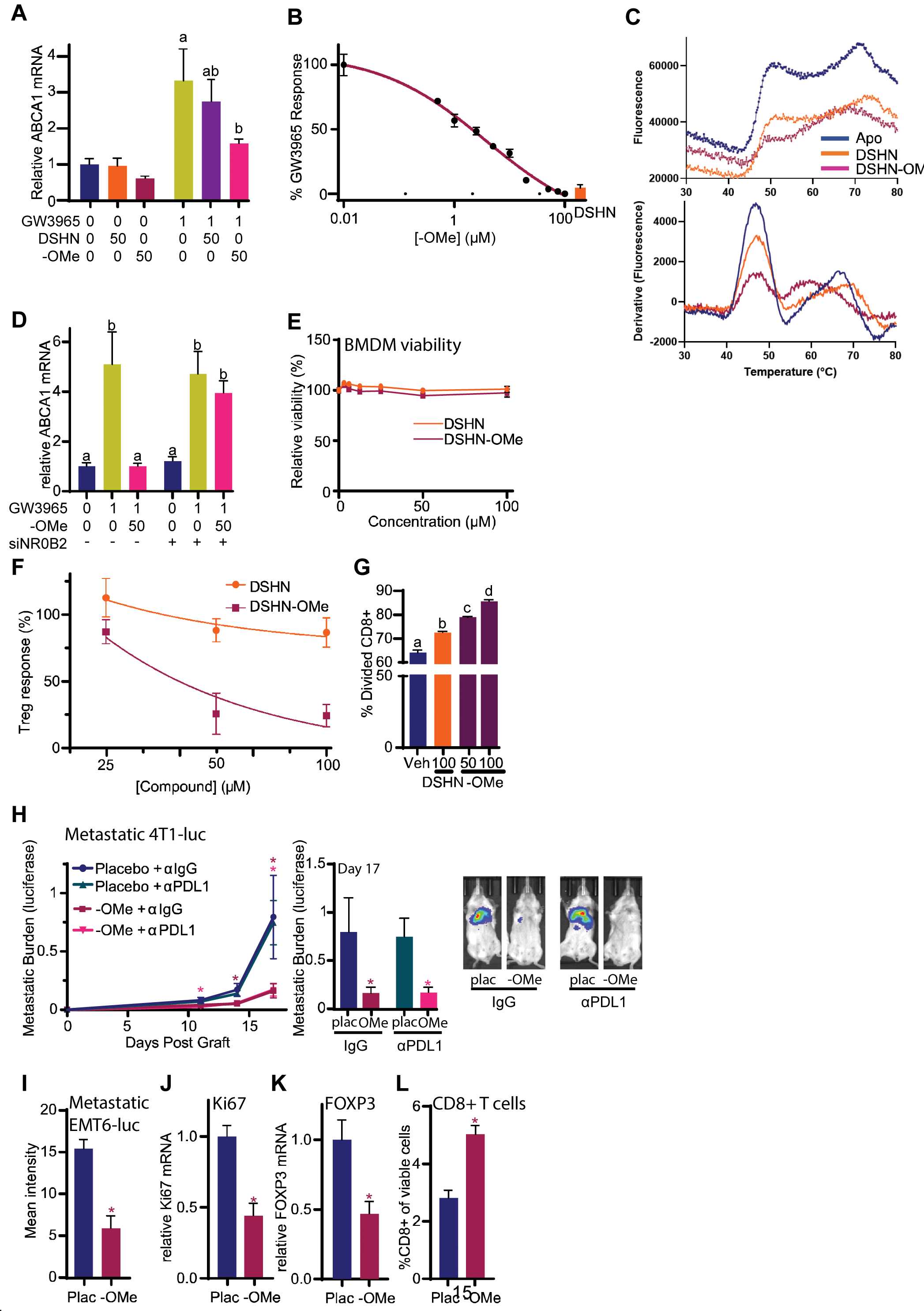
DSHN-OMe, an NR0B2 small molecule ligand with increased potency in terms of reducing T_reg_ expansion and strong efficacy against metastatic mammary tumors. **(A)** DSHN-OMe effectively reduces induction of LXR target gene ABCA1 by GW3965. Different letters denote statistical difference between GW3965 treated groups (N=3-4/group, 1-way ANOVA followed by Šidák test). **(B)** Dose response assay in BMDMs pre-treated with increasing doses of DSHN-OMe and then treated with 1µM GW3965, using ABCA1 mRNA as an endpoint. Data was normalized to 0.01µM DSHN-OMe (0%) and GW3965 alone (100%). DSHN (100µM) is indicated for comparison. Data was fit to a 4 parameter variable slope model, and IC_50_ estimated at 3.8µM. **(C)** Thermal shift assay using a recombinant protein presence of a fluorescent dye. Plot of raw data is above plot of first derivative. **(D)** DSHN-OMe is ineffective at regulating ABCA1 when NR0B2 is knocked down with siRNA in BMDMs. **(D)** DSHN-OMe has minimal effects on BMDM viability (N=4-6/group). **(F)** DSHN-OMe treated BMDMs reduce T_reg_ expansion in a dose related manner, with an increased biological maximum compared to DSHN (data normalized to mean vehicle, which was set at 100%. Non-linear three parameter regression shown, N=4/point). **(G)** T cells expanded in the presence of DSHN- or DSHN-OMe-treated BMDMs were less suppressive than those expanded in the presence of vehicle treated BMDMs, in terms of a subsequent co-culture of naïve T cells. The percentage of divided CD8+ T cells in the second round was quantified. (N=3-5/group, different letters indicating statistical significance, 1-way ANOVA followed by Tukey). **(H)** DSHN-OMe is an effective treatment against metastatic 4T1 mammary cancer growth. 4T1 cells were grafted intravenously, and lung metastatic lesions allowed to establish for 3d before treatment commenced. Metastatic burden through time is presented to the left of metastatic burden on the final day of imaging in the middle panel (N=12-17/group, mixed-effects model followed by Tukey’s multiple comparison test). Representative luciferase images in the right panel. **(I)** DSHN-OMe is an effective treatment against metastatic EMT6 mammary cancer growth. EMT6 cells were grafted intravenously, and lung metastatic lesions allowed to establish for 3d. DSHN-OMe was administered on days 4, 5, 8, 9 and 10 post-graft. 11d post-graft lungs were imaged *ex vivo* for luciferase (N=15/group). **(J)** EMT6 metastatic lungs were assessed for Ki67 mRNA and **(K)** FOXP3 mRNA. **(L)** EMT6 metastatic lungs were assessed for CD8+ cytotoxic T cells (CD8+). Asterisks indicate statistically significant differences.

In strong support of our preclinical findings, NR0B2 expression was inversely correlated with FoxP3 (**Fig. 1L. SFig. 6E**). When assessing all tumor types together, this inverse correlation was observed whether just the tumor nest or stromal sections were considered. However, when assessing specific tumor subtypes, only TNBC cases had the inverse correlation within the stromal compartment, while all three subtypes exhibited the inverse correlation within the tumor area itself (**Fig. 1L, SFig. 6E**).

### Myeloid cell loss of NR0B2 promotes tumor growth and metastasis in preclinical models

Given the strong correlational data observed in human patients (**Fig. 1, SFig. 5-6**), and our finding that NR0B2 in myeloid cells skews T cell expansion away from T_regs_, it was important to directly test whether NR0B2 altered tumor pathophysiology. Germline NR0B2^-/-^ mice were bred with MMTV-PyMT mice, that develop mammary tumors with progression reflective of human disease. Intriguingly, total tumor burden reached an endpoint size sooner in mice lacking NR0B2 compared to wildtype controls (**Fig. 2A**). Importantly, tumors from mice lacking NR0B2 had an increased T_reg_ and decreased T_eff_ infiltrate as determined by flow cytometry of digested tumors (**Fig. 2B**). Interestingly, the infiltrate of both DC- and M2-like macrophages was also increased in NR0B2^-/-^ tumors (**SFig. 7A**).

In order to evaluate the myeloid cell specific contributions of NR0B2, we generated NR0B2^fl/fl^;LysMcre^+^ mice ^18^. Syngeneic grafts with mammary tumor line E0771 grew faster in NR0B2^fl/fl^;LysMcre^+^ compared to control mice NR0B2^+/+^;LysMcre^+^ (**Fig. 2C**). Similar to the NR0B2^-/-^;PyMT^+^ tumors, E0771 tumors from myeloid cell specific knockout mice had increased T_regs_, which were also observed in lung metastatic lesions (**Fig. 2D, SFig. 7B**). They also had increased myeloid cell infiltrate in general, although among these cells there was no difference in MHCII positivity (a metric of the capacity to present antigen), but did have increased positivity for the immune checkpoint PD-L1 (**SFig. 7C**). This observation is likely significant given that high checkpoint levels are one mechanism for immune suppression^23,24^. The difference in tumor growth was consistent across two other syngeneic cancer models (melanoma and lung), suggesting that this axis is conserved between tumor microenvironments of different solid tumors (**Fig. 2E-F**, for the Lewis lung model a sub-optimal number of cells were grafted resulting in poor tumor growth in wildtype mice and allowing for increased time for an immune-mediated response).

### Tool compound, DSHN, an agonist of NR0B2 reduces breast tumor progression in preclinical models

T_regs_ have long been known to promote tumor progression and hinder the efficacy of various treatments, however, the main differentiating protein, FoxP3 has proven challenging to target therapeutically. Therefore, our finding that NR0B2 within myeloid cells results in skewed T_reg_ expansion presents an opportunity to alter T_reg_ abundance indirectly. Being a nuclear receptor, NR0B2 is highly amenable to small molecule modulation. One such compound, DSHN, has previously been described as an NR0B2 agonist^19^.

Initial characterization found that BMDMs treated with DSHN reduced subsequent T_reg_ expansion in a dose related manner (**Fig. 3A**). Since myeloid cells are thought to have been uniquely polarized within the tumor microenvironment, and in order to determine whether DSHN could overcome this, we isolated DCs (CD11B^+^;CD11C^+^) from MMTV-PyMT tumors. When these DCs were cultured in the presence of DSHN followed by LPS and IFNγ, they had decreased PD-L1 positivity, expectedly, the opposite to what was observed when NR0B2 was knocked out (**SFig 7D**). Importantly, they also had increased viability compared to vehicle treated DCs from the same tumor (**Fig. 3B** left panel). When co-cultured with CD4^+^ T cells, DSHN-treated DCs resulted in a decreased expansion of T_reg_ cells, indicating that this agonist may be useful in re-educating the myeloid tumor microenvironment (**Fig. 3B**, right panel). Cd4+ T cells expanded in co-culture with DSHN pre-treated BMDMs had decreased suppressive capacity in terms of secondary CD8+ expansion (**Fig. 3C**). Functionally, DSHN was able to reduce T_reg_ abundance in an *in vivo* model of immune tolerance to a chronic antigen, where T_regs_ are typically increased (**Fig. 3D**). Collectively, these results were consistent to those found in human cells as described earlier (**Fig. 1H**).

We then evaluated DSHN in a model where a low, suboptimal number of E0771-luc cells were grafted orthtopically (**Fig. 3E**). Since NR0B2 was also implicated in regulating myeloid cell expression of PD-L1 (**SFig. 7**), we investigated the effect of acute DSHN combined with anti-PD-L1 (αPDL1) treatment on subsequent outgrowth of E0771 tumors. Treatment was ceased after 18d and mice were monitored through time. Intriguingly, one DSHN-only treated mouse lived for more than 400 days post graft with no palpable tumor forming, while DSHN combined with αPDL1 significantly increased survival time compared to placebo treated mice (**Fig. 3E**). We next assessed the effects of DSHN combined with αPDL1 on growth of primary tumors (optimal number of cells used to establish graft). Treatment of animals bearing established E0771 or 4T1 tumors with DSHN combined with αPDL1 resulted in decreased growth (**Fig. 3F,G**). We then evaluated the ability of DSHN to impact established metastatic disease – a stage that is refractory to standard of care therapy and immune checkpoint blockade. As expected, αPDL1 did not significantly impact metastatic outgrowth in this model (**Fig. 3H**). However, DSHN alone slowed the growth of metastatic lesions. Furthermore, combining DSHN with αPDL1 reduced outgrowth even further than DSHN alone, indicating that simultaneously targeting the innate and adaptive arms of the immune system will be beneficial for the treatment of immune-suppressive tumors such as breast cancer. In summary, the tool compound DSHN provides proof of principle that the NR0B2-myeloid cell-T_reg_ axis can be targeted for the treatment of breast cancer.

### Inflammasome implicated in myeloid cell NR0B2 – T_reg_ axis

In order to gain insight into the potential mechanisms of action, we compared mRNA profiles from (1) loss of NR0B2 in PyMT tumors (from **Fig. 2A**) (2) *in vitro* BMDMs treated with the agonist DSHN, and (3) 4T1 metastatic lungs from mice treated with DSHN (**SFig. 8A-C**). Since NR0B2 is known to inhibit LXR, we also included groups of BMDMs treated with the synthetic LXR agonist GW3965, and both GW3965 and DSHN (**SFig. 8C**). A signature of upregulated genes in BMDMs treated with DSHN versus vehicle was found to be correlated with survival in breast cancer patients (**SFig. 8D**). In all three data sets, we observed that several genes associated with the inflammasome were altered (**SFig. 8E**). Given previous reports that the inflammasome can drive T_reg_ differentiation and expansion via myeloid cells^25^, we interrogated aspects of the inflammasome from upstream regulation to downstream execution (overview cartoon in top panel of **Fig. 4**). Intracellular Ca^2+^ was decreased when NR0B2 was overexpressed in naïve BMDMs or DCs treated with ionomycin or PMA, and *vice versa* when NR0B2 was knocked out (**Fig. 4A-B, SFig. 9A**). Reactive oxygen species (ROS) were increased in the absence of NR0B2 (**Fig. 4C**). S100A8 and S100A9 transcripts were altered by NR0B2 in CD11B+ myeloid cells isolated from tumors, or BMDMs co-cultured with E0771 cancer cells (**Fig. 4D, SFig. 9B-C**). Lysosome stability (ie: increased pH) was decreased when NR0B2 was knocked out in BMDMs or DCs (**Fig. 4E, SFig. 9D**). Cathepsin B mRNA was regulated when NR0B2 was manipulated in BMDMs or CD11B+ cells from tumors (**SFig. 9E**). Likewise, secreted cathepsin B protein was decreased when NR0B2 was overexpressed, or increased when NR0B2 was knocked out in BMDMs, CD11B+ cells isolated from tumors or DCs (**Fig. 4F, SFig. 9F**). When NR0B2 was overexpressed, the downstream executioner of the inflammasome, caspase 1, had decreased transcript (**SFig. 9G**) and activity (**Fig. 4G**), with the reverse being true in CD11B+ or CD11C+ cells from E0771 tumors grown in mice lacking myeloid NR0B2 (**Fig. 4H, SFig. 9H-I**). IL-1β mRNA was regulated in opposite ways when NR0B2 was knocked out or overexpressed (**SFig. 9J-M**), as was its secreted protein (**Fig. 4I-J, SFig. 9N-O**). These data suggest that NR0B2 is suppressing several different steps in the inflammasome cascade. Indeed, even when the inflammasome was activated with S100A8/9 and ionomycin, overexpression of NR0B2 blunted the induction of caspase 1, IL1β, IL18 and several downstream chemokines (**SFig. 9P**). IL-1β has previously been shown to promote FOXP3 expression and promote Treg differentiation^26^. Knockdown of IL-1β in BMDMs resulted in decreased T_reg_ expansion, confirming that decreased IL-1β is likely the downstream mechanism of NR0B2 (**SFig. 9Q**). Importantly, the increased T_reg_ expansion when cultured with BMDMs lacking NR0B2 was lost when BMDMs were treated with siRNA against caspase 1, siRNA against IL-1β (**Fig. 4K**), or an antibody to neutralize IL-1β (**Fig. 4L**), providing strong support for inflammasome pathway modulation being the primary mechanism by which NR0B2 influences T_reg_ expansion.

### Development of a more efficacious NR0B2 agonist

While DSHN was a useful proof-of-concept small molecule, macrophages required high doses and chronic exposure for robust influences on subsequent T_regs_ expansion. Furthermore, it proved to have poor aqueous solubility, potentially limiting its *in vivo* translation. Therefore, we synthesized several derivatives of DSHN and screened them using a reporter assay which makes use of the endogenous cholesterol homeostatic feedback loop, where FXR activation upregulates NR0B2, which then inhibits LXR and its normal induction of ABCA1 (**SFig. 10A,B**). We then followed this screen with several others with stringent cutoffs: effect on endogenous ABCA1 and IL1β expression, reduction of T_reg_ expansion in either macrophage or DC co-cultures, and low cellular toxicity (**Fig. 5A, SFig. 10C**). The methyl ester of DSHN (DSHN-OMe) emerged as the best performer across all of these endpoints (preparation and characterization in **Supplementary Materials**), with DSHN-OMe reducing LXR induction of ABCA1 in a dose-related manner (**Fig. 5B**; dose-related response of DSHN depicted in **SFig. 11A**). DSHN-OMeshowed a modest shift in NR0B2 thermal melt temperature. The T_M_ for apo/unliganded was 46.74 ± 0.22, DSHN-OMe at 46.92 ± 0.17 and DSHN at 46.34 ± 0.51°C (**Fig. 5C**). Both molecules similarly reduced the magnitude of the transition suggesting that they are likely binding to the protein but not significantly affecting thermal stability. The ability of DSHN-OMe to attenuate the induction of ABCA1 expression by the LXR agonist GW3965 was lost when NR0B2 was knocked down in BMDMs, indicating that DSHN-OMe was working through its intended target (**Fig. 5D**) further suggesting NR0B2 as the target of DSHN-OMe. Similar to DSHN, DSHN-OMe showed minimal cytotoxicity in BMDMs (**Fig. 5E**). However, compared to DSHN, DSHN-OMe treated macrophages or DCs elicited a reduction in T_reg_ expansion with a lower dose, and had a larger biological maximum (**Fig. 5F, SFig. 11B**). Pretreatment of BMDMs followed by washout prior to co-culture with T cells resulted in decreased T_reg_ expansion for DSHN-OMe, but only modest reductions for DSHN, indicating that (i) the effects of DSHN-OMe are mediated through BMDMs/DCs, and (ii) DSHN-OMe either had better cell penetration or more durable effects on BMDMs compared to DSHN (**SFig. 12A**). Importantly, T cells resulting from expansion in co-culture with BMDMs that were pretreated with either DSHN-OMe or DSHN were less suppressive when subsequently cultured with a second set of naïve T cells, and specifically assessing CD8+ expansion (**Fig. 5G**). Conversion of a carboxylic acid to an ester is often associated with better cellular uptake^27^. Indeed, DSHN had very poor uptake by RAW264.7 cells, while DSHN-OMe had considerable uptake at 4h. Conversion of DSHN-OMe to DSHN was minimal over the 24h time period of the experiment, suggesting that DSHN-OMe is likely the active modulator of NR0B2 (**Fig. S12B**).

Both DSHN and DSHN-OMe were able to blunt the expression of inflammasome associated genes in the presence of both a priming signal as well as an activation signal (LPS and nigericin respectively), but not with just a priming signal, indicating that the effects of NR0B2 are primarily with respect to activation of the inflammasome (**SFig. 13**).

Importantly, DSHN-OMe significantly reduced the outgrowth of established 4T1 metastatic lesions as a single agent (**Fig. 5H**). Addition of αPDL1 did not significantly influence the effects of DSHN-OMe in this assay, although the short time frame (due to humane endpoint of placebo groups) may not have captured the contribution of checkpoint inhibition. DSHN-OMe also significantly reduced metastatic burden in mice grafted with EMT6 cells, an alternative murine mammary cancer model (**Fig. 5I**). Ki67 mRNA was also decreased in EMT6 metastatic lungs of DSHN-OMe treated mice, indicating a decreased number of proliferating cancer cells (**Fig. 5J**). As expected, metastatic lungs from DSHN-OMe treated mice had decreased FOXP3 expression (**Fig. 5K**), and a corresponding increase in the relative abundance of CD8+ cytotoxic T cells (**Fig. 5L**).

## Discussion

NR0B2 is best known as a regulator of cholesterol and bile acid homeostasis, directly binding and inhibiting LXR and LRH-1 mediated induction of cholesterol catabolism and efflux. Emerging reports indicate roles outside this axis as well as extra-hepatic roles^28–31^. Various roles for NR0B2 in the immune system have been described^17,18^. Here, we demonstrate the importance of NR0B2 in breast cancer pathophysiology. By inhibiting the inflammasome within myeloid cells, NR0B2 results in reduced T cell expansion towards T_regs_, reduced immune suppression and ultimately, reduced tumor or metastatic outgrowth. Previous reports have indicated a role for NR0B2 in hepatic carcinomas, but have largely focused on cancer-cell intrinsic roles^17,32–34^. In the models of breast cancer evaluated here, NR0B2 had no significant effects on cellular proliferation or migration, indicating that these cell-intrinsic activities may be exclusive for cells of the hepatic origin. Importantly, NR0B2, as with all nuclear receptors, has a distinct ligand binding domain allowing for targeting with small molecules. Indeed, a previously reported agonist, DSHN was able to improve the efficacy of ICB, and the novel DSHN-OMe decreased metastatic outgrowth as a single agent in two models of TNBC. Several correlations using human breast cancer patient data support our preclinical observations: NR0B2 mRNA expression is associated with a good prognosis, NR0B2 expression is inversely correlated with the T_reg_ marker FOXP3 mRNA in the TCGA and FOXP3 protein in an independent cohort from France, and a signature of genes upregulated in BMDMs treated with DSHN. Furthermore, human PBMCs treated with DSHN also attenuated the expansion of T_regs_. Therefore, this target and axis demonstrate significant potential for translation.

There is significant effort to develop new ICB targets or improve the efficacy of current ICB. Cholesterol metabolism and homeostasis is emerging as a central axis regulating immune function. Cholesterol itself plays important roles in T cell biology^35^, in addition to LXR activation^36^. Cholesterol metabolites shift myeloid cells to being highly immune suppressive, in part through LXR activation^6,15,37^. However, LXR activity in myeloid immune cells appears complex, and these receptors are likely selectively modulated with different ligands exerting varying effects^16^; this complexity providing pharmacological opportunity. Downstream cholesterol homeostasis and bile acid signaling have also been implicated, with microbial transformed bile acids shown to alter T_reg_ expansion in a T cell-intrinsic manner^38,39^. There is some evidence that these effects are mediated through the bile acid receptors Farnesoid X Receptor (FXR) or TGR5^38,40^, but this still requires further investigation. NR0B2 is a direct target gene of FXR, leaving the possibility that NR0B2 may also have T cell-intrinsic effects, in addition to the myeloid cell effects described here. Regardless, the effects would be expected to be beneficial with respect to cancer therapy.

Previous work has described NR0B2 in regulating various aspects of immune function. Mice with a whole-body knockout of NR0B2 were protected from septic shock induced by LPS^41^. Several mechanisms were proposed for the suppressive effects of NR0B2, including direct interactions with NF-κB and TRAF6, which in the case of TRAF6 led to a reduction in its ubiquitination. We have also shown that NR0B2 and NF-κB appear to colocalize, although direct binding or subsequent consequences of this interaction are less clear^18^. The observations that NR0B2 is protective against septic shock are in contrast to what we would expect if T_regs_ were increased in the absence of NR0B2. However, the authors also note that upon LPS stimuli, macrophages upregulate NR0B2 via a Ca^2+^-dependent activation of AMPK, which would fit a model whereby NR0B2 is upregulated to reduce T_reg_ expansion and drive an immune-response. Interestingly, NR0B2 was previously found to directly interact with NLRP3 to attenuate the NLRP3 inflammasome^42^. While this may be the case, our data indicates that to some degree NR0B2 regulates every aspect of the inflammasome investigated: Ca^2^ flux, ROS generation, S100A8 and S100A9 mRNA regulation, lysosome stability, cathepsin B (mRNA and protein), caspase 1 activity and IL-1β (mRNA and protein). This subtle regulation across the board indicates the critical role for NR0B2 in shaping the myeloid immune response. Importantly, these observations were made in both BMDMs and DCs, suggesting that the same target (NR0B2) will have utility across all antigen presenting cells. Ultimately, the reduced inflammasome activity resulted in decreased expansion of T_regs_, as knockdown of caspase 1 or IL-1β, or immune-neutralization of IL-1β were able to inhibit the effects of loss of NR0B2. There is indication that NR0B2 also regulates TGF-β and IL-2, two other cytokines involved in T_reg_ differentiation/expansion^18^. Whether these are downstream effects of an attenuated inflammasome remains to be determined. Collectively, NR0B2 appears to subtly re-educate myeloid cells across several aspects of the inflammasome and cellular function. These actions subsequently decrease T_reg_ expansion, shifting the tumor microenvironment towards being less immune-suppressive.

Being a nuclear receptor, NR0B2 has a well-defined ligand binding domain, through which small molecules can bind and regulate its activity. Here, we show that a first generation NR0B2 agonist, DSHN, regulates T_reg_ expansion as expected after myeloid cell treatment, and when combined with ICB reduces the tumoral outgrowth of syngeneic mammary tumors. Using the 4T1 model of mammary metastasis, DSHN had significant effects as a single agent, perhaps reflecting the variable immune-suppressiveness in different microenvironmental niches. *In vitro*, DSHN required high doses and continuous exposure in order to alter T_reg_ expansion, and lacked the solubility for effective translation as a therapeutic. Therefore, we developed and screened for improved derivatives, identifying DSHN-OMe as having increased efficacy in terms of T_reg_ expansion. DSHN-OMe also had superior qualities in terms of low cellular toxicity. In two models of murine mammary cancer metastasis, DSHN-OMe had robust effects as a single agent.

To identify DSHN-OMe, we made use of the endogenous cholesterol homeostasis feedback loop, whereby NR0B2 inhibits LXR. The bile acid receptor, FXR, serves to up-regulate NR0B2 when activated. Thus, in addition to targeting NR0B2 directly, it may be possible to utilize the FXR. In this regard, the semi-synthetic bile acid, obeticholic acid is FDA approved for the treatment of primary biliary cholangitis, and may represent a rapidly translatable avenue for this axis. Obeticholic acid is associated with several side effects including pruritus, resulting in low patient compliance. This provides rationale for the continued development of downstream targets such as NR0B2. However, it may also be possible to use obeticholic acid acutely to re-educate the tumor microenvironment making it more susceptible to ICB. Collectively, our data provide proof of concept that mediators of downstream cholesterol homoeostasis can be leveraged for the treatment of solid tumors.

## Methods

DSHN was synthesized by Sai Life (Hyderabad, India). GW3965, PMA, Ionomycin, Obeticholic acid and GW4064 were obtained from Cayman (Ann Arbor, MI). DSHN-OMe was synthesized in house. Anti-CD4, anti-Foxp3, anti-CD25, anti-CD8, anti-CD3, and anti-CD45 were from BD Biosciences. Carboxy fluoroscein succinimidyl ester (CFSE) and Cell Trace Violet (CTV) were from BioLegend (San Diego, CA) and Invitrogen (Thermo Fisher Scientific, USA). LIVE/DEAD fixable viability kits were from Invitrogen. Foxp3/transcription factor staining buffer set (00-5523-00) was purchased from eBioscience. IL-1β (432604) and Cathepsin B (EA100429) enzyme-linked immunosorbent assay (ELISA) kits were from BioLegend (San Diego, CA) and Origene (Rockville, MD) respectively. Fetal bovine serum (FBS) was from Cytiva HyClone. Non-essential amino acids, sodium pyruvate, penicillin/streptomycin, DPBS without Ca^2+^ and Mg^2+^, cell stripper, trypsin and RPM1-1640 were purchased from Corning. GlutaMAX, ACK lysis buffer, β-mercaptoethanol and Type II Collagenase were from Gibco (Thermo Fisher Scientific, USA).

HepG2 cells were a gift from Sayeepriyadarshini Anakk (University of Illinois at Urbana-Champaign). EMT6-luc cells were a gift from Hasan Korkaya (Augusta University). E0771 and 4T1-luc cells were a gift from Mark Dewhirst (Duke University). RAW 264.7, Lewis lung carcinoma and B16 cells were purchased from American Type Culture Collection. Cell lines were not cultured longer than 2 months after thawing or after passage 20. Cell lines were routinely tested for mycoplasma.

### Survival and Correlational Analysis of human tumors

Survival analysis in **Fig. 1** was performed the Kaplan-Meier Plotter webtool (https://kmplot.com/analysis/)^43^ and cBioPortal (https://www.cbioportal.org/)^44–46^. The Kaplan-Meier Plotter webtool uses aggregated data from GEO, EGA, and TCGA. Differentially upregulated genes between vehicle and DSHN treated BMDMs from RNA-seq analysis (fold change threshold of 2 fold, and FDR<0.01), were used to create a non-weighted signature of 22 genes. This signature was then used to probe the METABRIC dataset as obtained through cBioPortal. Tumors were then parsed into upper and lower quartiles and used to examine survival data using Kaplan-Meier analysis, similar to^47^.

### Animal tissue and in vivo studies

All protocols involving animals were approved by the Institutional Animal Care and Use Committee (IACUC) at the University of Illinois Urbana-Champaign. Female wild-type C57BL/6 and BALB/C mice were purchased from Charles River Laboratory. Mice were 8–12 weeks old at the start of the experiment. Founder OT-II mice and MMTV-PyMT were purchased from Jackson Laboratory and bred in-house. Founder NR0B2^fl/fl^ mice were a kind gift from John Auwerx and Kristina Shoonjans (Ecole Polytechnique de Lausanne). The mice were matched for age in each experiment.

### Preparation of Bone Marrow-Derived Macrophages (BMDMs) and Dendritic Cells

Bone marrow cells were collected from mouse tibia and femur. Cells were passed through 70-micron cell strainer and subsequently cultured 10mL complete RPMI media supplemented with 20 ng/mL of recombinant murine M-CSF (576406; BioLegend, 315-02 PeproTech). On day 3, additional 5 ml of the complete media was added. On day 7, media was replaced. BMDMs were harvested on day 10 using cell stripper solution for further experiments. Dendritic cells were isolated from the spleens of wildtype, using mouse CD11c UltraPure MicroBeads (130-125-835; Miltenyi Biotec) according to the manufacturer’s instructions) and cultured in RPMI medium supplemented with 10% heat-inactivated charcoal-stripped FBS, 50 µM β-mercaptoethanol, 1% sodium pyruvate, 1% nonessential amino acids, % penicillin/ streptomycin, and 1% Glutamax.

### Cell Isolation

Fresh tumors, and lungs were collected separately from mice, and digested in DMEM/F12 supplemented with 2mg/ml type II collagenase and 1% penicillin/streptomycin for 45 mins at 37°C while shaking. Subsequently, cells were passed through a 70-µm filter into single-cell suspension, washed with FACS buffer, incubated with ACK lysis buffer for 1 min, and washed with FACS buffer before antibody staining for flow cytometry. Single cell suspensions of spleens were obtained by mechanical dissociation through a 70-µm filter in isolation buffer (DPBS supplemented with 0.5% BSA and 2mM EDTA). Subsequently, cells were washed with isolation buffer, incubated with ACK lysis buffer, and washed with isolation buffer before antibody staining for flow cytometry or immune cell isolation. Cd11b+ (130-126-725) and Ly6G+ (130-120-337) were isolated from single cell suspensions using UltraPure MicroBeads from Miltenyi Biotec according to the manufacturer’s instructions.

### In vitro T_reg_ Expansion Assays

Assays were performed similar to previously described^15,18^. T cells were cultured in RPMI medium supplemented with 10% heat-inactivated charcoal-stripped FBS, 50 µM β-mercaptoethanol, 1% sodium pyruvate, 1% nonessential amino acids, 1% penicillin/ streptomycin, and 1% Glutamax. Naïve CD4+ T cells were isolated from the spleens of wildtype or OT-II mice, using Naive CD4+ T Cell Isolation Kit (130-104-453; Miltenyi Biotec) according to the manufacturer’s instructions. T cells were labeled with the vital dye 2.5 μM CFSE or 5 μM Cell Trace Violet according to the manufacturer’s instructions. For wildtype T_reg_ expansion in presence of antigen presenting cells (APCs), naïve CD4+ T cells were co-cultured with APCs at indicated ratio in the presence of 0.5 μg/mL (anti-CD3 BioLegend), 0.5ng/mL TGFβ (BioLegend) and 0.5ng/mL IL-2 (BioLegend) for 72 hr at 37 °C. For OT-II Treg expansion, OTII naïve CD4+ T cells from OT-II mice were activated by antigen presentation by culturing with BMDMs primed with 10 μg/mL OVA_323–339_ (Bachem) and 0.5 μg/mL LPS (Sigma) or dendritic cells primed with 10 μg/mL OVA_323–339_. To deplete IL1β (clone B122; BE0246; BioXCell) and IL18 (clone YIGIF74-1G7; BE0237; BioXCell), isotype control or respective neutralizing antibodies were added at 5 μg/mL to T cell cocultures. The proliferation of T cells was assessed by flow cytometry 72h after co-culture at indicated ratio.

### In vitro T Cell Suppression Assays

BMDMs were cocultured with naïve CD4+ T cells and differentiated into T_regs_ in the presence 0.5 μg/mL (anti-CD3 BioLegend), 1ng/mL TGFβ (BioLegend) and 1ng/mL IL-2 (BioLegend) for 72 hr. Next, expanded CD4+ T cells were cocultured at different ratios with CFSE/CTV labeled CD8+ T cells and stimulated with Mouse T-Activator CD3/CD28 Dynabeads at 2 cells per bead ratio and 0.5ng/mL IL-2 (BioLegend). Cell culture media consisted with RPMI 1640, supplemented with 10% heat-inactivated charcoal-stripped FBS, 50 µM β-mercaptoethanol, 1% sodium pyruvate, 1% nonessential amino acids, 1% penicillin/ streptomycin, and 1% Glutamax. CD8+ T cells were isolated from wildtype mice spleens using CD8a+ T Cell Isolation Kit (130-104-075) from Miltenyi Biotec according to the manufacturer’s instructions. Flow cytometric analysis of suppressive cultures was preformed after 72 h. Beads were removed using DynaMag magnet (Invitrogen) prior to flow cytometry staining and analysis.

### Flow Cytometry Staining and Analysis

For surface staining, cells were stained with LIVE/DEAD fixable viability dye diluted in PBS (1:1000) for 30 minutes at 4°C in the dark, washed and subsequently stained with fluorochrome-conjugated antibodies for cell surface antigens in FACS buffer (DPBS supplemented with 2% FBS and 1% penicillin/streptomycin) at 1:100 for 30 minutes at 4°C in the dark. Next, cells were fixed with 4% Formalin and incubated at 4°C in the dark until analysis. For intracellular staining, cells were stained with intracellular cytokines according to the manufacturer’s protocol using the ebioscience Foxp3/transcription factor staining buffer set. In brief, after surface staining, live cells were fixed and permeabilized in 1× fix/perm buffer for 30 minutes at room temperature or overnight at 4°C in the dark. Fluorochrome antibodies against intracellular antigens were diluted in 1× permeabilization buffer at 1:50 and stained for 30 minutes at room temperature. Samples were washed serially with 1× permeabilization buffer and FACS buffer and resuspended in FACS buffer for analysis. Cytometry data were acquired on either BD LSRII, BD Fortessa, BD Symphony A1, BD Accuri C6 and Thermo Attune NxT.

### Gene Silencing and Overexpression

For gene knockdown small interfering RNA was delivered using HiPerFect transfection reagent (Qiagen). BMDMs were transfected with either negative control or Caspase-1/IL-1β-complementary siRNA (SMARTpool, ON-TARGETplus siRNA, Dharmacon) at 50nM. NR0B2 siRNA was from Sigma. Expression of the target genes after the knockdown was assessed using qPCR. For NR0B2 overexpression, NR0B2 overexpressing plasmid was delivered using Effectine transfection reagent (Qiagen). All transfections were conducted in accordance with the manufacturer’s protocol. Media was changed 12 h post-transfection and cells were harvested for subsequent experiments 48 h post transfection.

### Screening to identify NR0B2 agonists

HepG2 cells were co-transfected with ABCA1 renilla reporter plasmid and TK-control firefly luciferase vector at 20:1 using Lipofectamine 3000 reagent according to manufacturer’s instructions for 24hrs. Afterwards, 7000 cells were seeded in to 384 well f-bottom, white plates (Greiner Bio-One). Test compounds were added to the plates in a final working volume of 30 µL (for a final concentration of 50 µM) in presence of DMSO or 1 µM GW3965 and incubated for 20-24hrs at 37°C. Luciferase activity was measured using the Luc-Pair Duo-Luciferase HT Assay kit (GeneCopoeia, Rockville, MD, USA) by adding luciferase substrates sequentially following manufacturer instructions. Luminescence was measured with 2 s integration times in a microplate reader (BioTek Cytation 5), 15 min after adding each substrate. The addition of reagents to the microplates was timed at each step to match the reading time delays and reading sequence of the microplate reader.

### Resazurin cell viability assay

20,000 cells were seeded into 96 well f-bottom, black plates (Corning Costar) and cell viability was measured using 100µL resazurin per well at 0.3 mg/mL (Acros Organics). The fluorescence signal was quantified by plate reader (560 nm excitation/590 nm emission) at 2-3 h.

### Inflammasome activation, RNA extraction, quantitative real PCR

When comparing inflammasome responses, BMDMs were treated with 500ng/mL LPS for 4 h 37 °C in complete RPMI 1640. After 4hr priming, 15 µM Nigericin was added for 45 mins at 37 °C. Next, media was removed, washed with PBS and inflammasome activation was assessed using qPCR or inflammasome caspase-1 activity was measured using Caspase-Glo® 1 inflammasome assay kit (G9951; Promega) according to manufacturer’s instructions.

### Measurement of Intracellular Ca2^+^ Signaling

Intracellular Ca2^+^ was measured using a Fluo-4 direct calcium assay kit (F10471; Invitrogen) according to manufacturer’s instructions. In brief, BMDMs, Cd11c^+^, or Cd11b^+^ cells were cultured in 96 well or 384 well black flat bottom plates (Corning Costar). The cells were incubated in 1x Fluo-4 Direct calcium reagent with probenecid for 30 mins at 37°C and 5% CO_2_ and 30mins at room temperature. Then cells were stimulated with ionomycin or PMA at indicated concentrations, immediately measured the fluorescence signal for excitation at 494 nm and emission at 516 nm using BioTek Cytation 5 plate reader.

### Measurement of Intracellular Cellular Oxidative Stress

Intracellular cellular oxidative stress was measured using CellROX™ Deep Red Reagent (C10422; Invitrogen) according to manufacturer’s instructions.

### Compound Uptake Assay

Cellular uptake of DSHN and DSHN-OMe was determined in RAW 264.7 cells cultured in 60×15mm tissue culture dishes. Cells were treated with vehicle (0.1% DMSO), 50 μM DSHN or 50 μM DSHN-OMe for 4 h, 8 h, or 24 h. After incubation, cells were harvested, washed with PBS and cell pellets were incubated at −80°C freezer. Next, cell pellets were resuspended in 200 μL 70:30 MeOH:H_2_0 and sonicated to lyse cells. Debris from lysed cell suspension was removed by centrifugation. Resulting supernatant was analyzed by LC-MS/MS.

### Protein expression, purification, and thermal shift assay

A gene containing a hexa-His-TEV fusion with full length NR0B2 in pET21(a)+ was transformed in *E.coli* BL21(DE3). A single colony was used to inoculate a 100 mL starter culture of LB broth containing 100 µg/mL ampicillin, which was allowed to grow overnight with shaking at 37°C. This starter culture was used to inoculate flasks containing autoinducing media, which were grown at 37°C with shaking until they reached an OD^600^ of 0.8. Subsequently, the temperature was reduced to 16°C and cells were allowed to grow for another 24 hours. After harvesting by centrifugation, cells were resuspended at 15 mL/g cell paste in a buffer comprised of 25 mM HEPES pH 8.0, 250 mM NaCl, 20 mM imidazole pH 8.0, 5% glycerol, 0.5 mM TCEP, and Roche EDTA-free protease inhibitor cocktail. Cells were lysed by sonication then centrifuged at 18,000 *xg* for 30 minutes to remove insoluble material. The lysate was loaded onto a gravity flow Ni-NTA column, washed with 5 column volumes (CVs) of resuspension buffer, and protein was eluted using the resuspension buffer supplemented with 500 mM imidazole. Collected protein was dialyzed overnight in the resuspension buffer without imidazole then concentrated and purified on a Superdex 200 HiLoad 200 16/600 size exclusion column. A peak corresponding to ∼29 kDa was collected and molecular weight was verified using SDS-PAGE. For thermal shift assays, vehicle (DMSO), 1 mM DSHN-OME, or 1 mM DSHN was incubated with 6 µM purified NR0B2 overnight at 4°C. We found that this concentration was the minimal required to give a reliable melt curve. The next morning the mixtures were centrifuged at 20,000 *xg* for 30 minutes to remove any insoluble protein or ligand. The supernatant was mixed with SYPRO orange (Thermo) then placed in 96-well qPCR plates in triplicate. Melt curves were obtained with a Applied Biosystems qPCR machine between 25 and 95°C at a gradient of 0.15°C/second. Melt curve data were fit using Thermo Protein Thermal Shift Software. These experiments were performed three independent times with three technical replicates each.

### Human Tumor Specimens

Deidentified serial breast cancer sections and corresponding subtype information were obtained from archival collections at the Département de Biologie et de Pathologie des Tumeurs, Centre Georges-François Leclerc, Dijon, France. They were stained and analyzed by the Research Histology and Tissue Imaging Core, University of Illinois at Chicago, Illinois, USA. Three consecutive sections from each sample were stained with dual immunostain for FoxP3 and panCK, RNAscope assay with NR0B2 probe, and dual immunostain for CD8 and panCK. Staining for all targets was performed on BOND RX autostainer (Leica Biosystems, Deer Park, IL) using preset protocols. For dual immunostaining, the first and third slides were stained with either FoxP3 (1:100, #12653, Cell Signaling Technology, Danvers, MA) or CD8 (1:100, #ACI3160, Biocare Medical, Pacheco, CA) antibodies using BOND Polymer Refine Detection kit (#DS9800), followed by staining with panCK antibody (1:4000, #M351501, Agilent, Santa Clara, CA) using BOND Polymer Red Detection Kit (#DS9390). The middle slide was stained with RNAscope 2.5 LS Probe - Hs-NR0B2 assay (#877508, Advanced Cell Diagnostics, Newark, CA) using RNAscope 2.5 Leica assay-RED reagents for hybridization and detection. The standard ACD Red Rev BOND Rx protocol was modified to include BOND Epitope Retrieval Solution 2 (Leica Biosystems) pretreatment for 25min at 95°C and to increase Amp5 step time to 30 minutes. Positive (Hs-PPIB, #313908) and negative (DapB, #3120) probe assays were included with each batch of samples. The stained slides were scanned at 40X magnification on PhenoIMager HT (Akoya Biosciences, Malborough, MA). All three slides were registered using HALO software (Indicalabs, Albuquergue, NM). HALO algorithms were used to segment tumor from stroma and count positive T cells and NR0B2 transcripts, at single-cell resolution. We computed Spearman correlation coefficients to evaluate associations between CD8 and FoxP3 density and NR0B2 expression in the tumor and stromal compartments. These correlations were also computed within strata defined by ER/PR, HER2 and triple negative status. CD8, FoxP3 and NROB2 expression was compared among the three subtypes using pairwise two-sided multiple comparison-adjusted analysis. All P values were two-sided with a 0.05 threshold for statistical significance.

## Acknowledgments

We would like to thank the patients whose tumors populated data in the TCGA, METABRIC and Human Protein Atlas initiatives, and the patients whose archival tumor tissues were used for NR0B2 staining. We would also like to thank our breast cancer advocate team: Sarah Adams, Renaé Strawbridge, Jamie Holloway, Lea Ann Carson, Susan Stewart and Catherine Applegate. We are grateful for comments and input from Sayeepriyadarshini Anakk (University of Illinois Urbana Champaign), advice on high throughput screening from Chen Zang (University of Illinois Urbana Champaign), and LC-MS/MS expertise from Lucas Li (Duke University) Founder NR0B2^fl/fl^ mice were a kind gift from John Auwerx and Kristina Shoonjans (Ecole Polytechnique de Lausanne). The Tumor Engineering and Phenotyping Core at the Cancer Center at Illinois assisted with Nanostring analysis.

## Notes

### Competing Interest Statement

ERN, PJN, SA, RF, HEVG and SHS have filed a provisional patent describing DSHN-OME and its use.

