## Supplementary Material for "Re-education of myeloid immune cells to reduce regulatory T cell expansion and impede breast cancer progression"

**Supplementary Materials Included:**

Page 2: Extended Methods and Results regarding ligand synthesis

Page 20: Figs. S1 to S13

### Extended Methods and Results

#### *Preparation and Characterization of DSHN derivatives*

*(corresponding derivative numbers are outlined in SFig. 10C).*

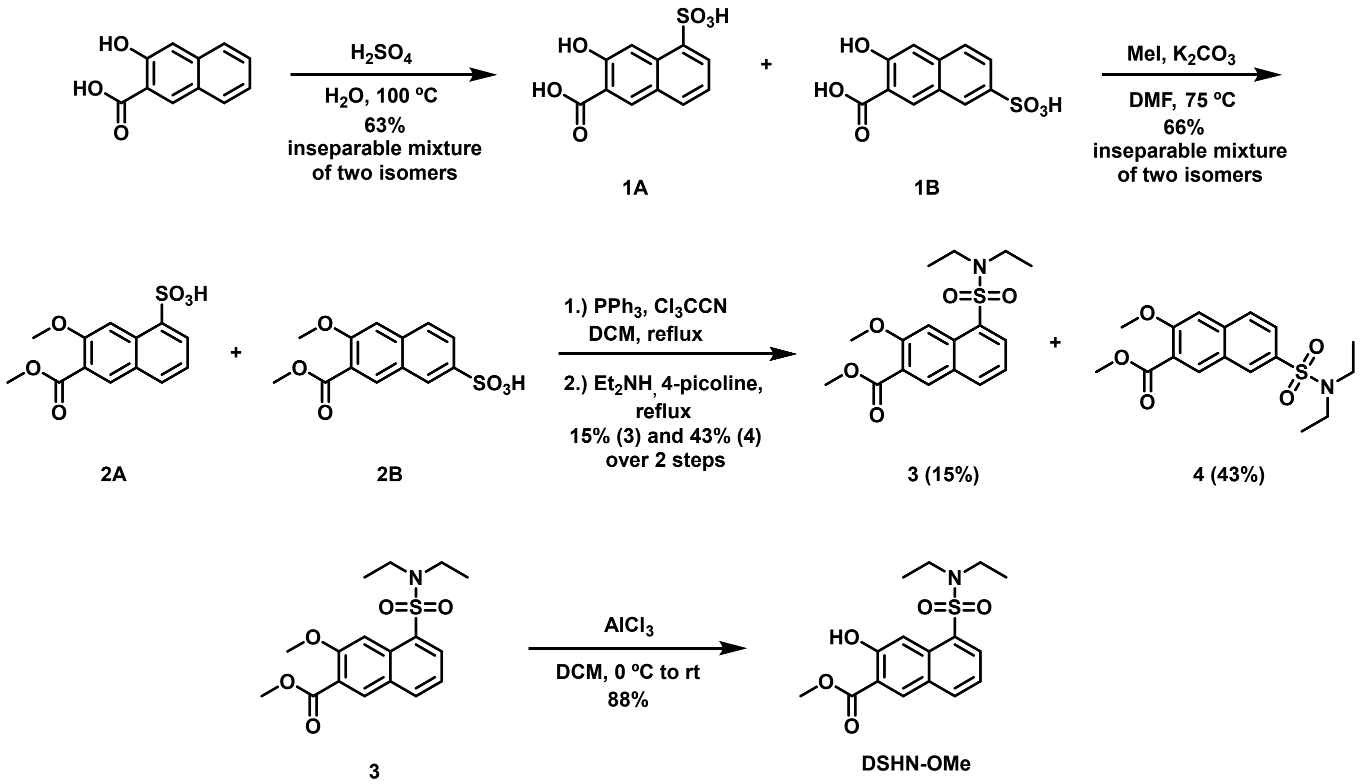


**Scheme 1. Synthesis of DSHN-OMe.**

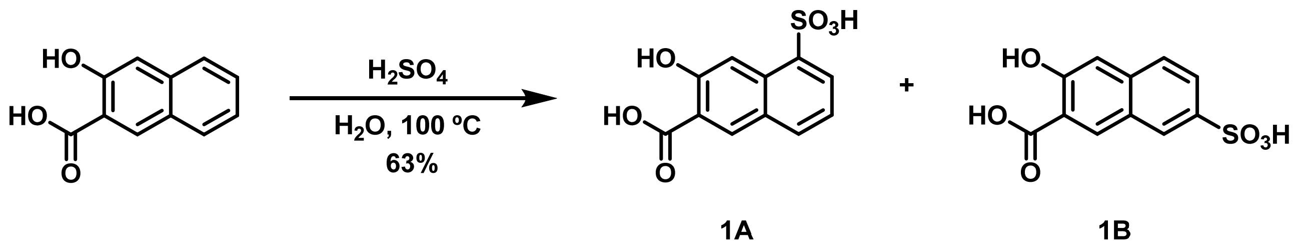


**Scheme 2. Synthesis of the mixture of 1A and 1B.**

**3-hydroxy-5-sulfo-2-naphthoic acid (1A) and 3-hydroxy-7-sulfo-2-naphthoic acid (1B)**

**Procedure:** To a 50 mL round bottom flask was added concentrated sulfuric acid (15 mL, 280 mmol, 5.3 eq.) and water (10 mL). The solution was heated to 100 °C and 3-hydroxy-2-naphthoic acid (10 g, 52.88 mmol, 1.0 eq.) was added. The reaction was stirred at 100 °C for 3 hours. The reaction mixture was then removed from heat and concentrated *in vacuo* to remove water. The crude product was purified via reverse phase flash column chromatography ($H_2O$ to 1:1 $H_2O$/Acetonitrile) yielding a 1:3 mixture of isomers **1A** and **1B**, respectively, as a yellow solid (9 g, 63%).

**$^1$H NMR** (500 MHz, Methanol-d4) δ **Major (1B)**: 8.62 (s, 3H), 8.32 (s, 3H), 7.89 (d, *J* = 8.7 Hz, 3H), 7.77 (d, *J* = 8.7 Hz, 3H), 7.30 (s, 3H). **Minor (1A)**: 8.58 (s, 1H), 8.24 (s, 1H), 8.18 (d, *J* = 7.2 Hz, 1H), 7.96 (d, *J* = 8.2 Hz, 1H), 7.34 (t, *J* = 7.7 Hz, 1H).

**$^{13}$C NMR** (125 MHz, Methanol-d4) δ **Major (1B)**: 172.91, 159.39, 139.72, 134.80, 127.90, 127.66, 127.17, 127.08, 123.24, 116.81, 112.26. **Minor (1A)**: 172.84, 158.51, 134.75, 134.50, 134.12, 133.79, 129.78, 129.23, 116.10, 112.51, 112.02.

**HRMS** (ESI): calc. for $C_{11}H_7O_6S$ $[M-H]^-$: 266.9964, found: 266.9959.

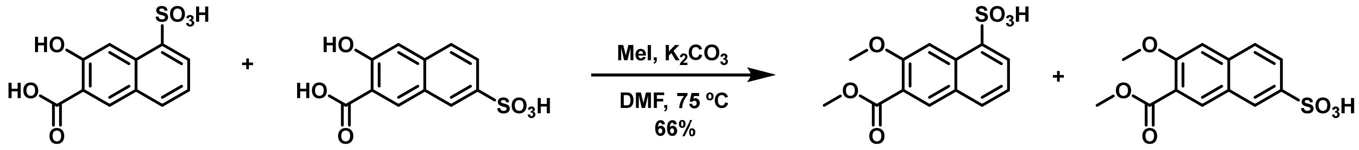


1A 1B 2A 2B

**Scheme 3. Synthesis of the mixture of 2A and 2B.**

**7-methoxy-6-(methoxycarbonyl)naphthalene-1-sulfonic acid (2A) and 6-methoxy-7-(methoxycarbonyl)naphthalene-2-sulfonic acid (2B)**

**Procedure:** To an oven dried 200 mL round bottom flask was added a mixture of isomers **1A** and **1B** (1.1 g. 4.1 mmol, 1.0 eq.) and $K_2CO_3$ (4g, 28.7 mmol, 7.0 eq.). The flask was purged with nitrogen and DMF (18 mL, 0.25 M) was added. Methyl iodide (2.3 mL, 36.9 mmol, 9.0 eq.) was added dropwise and the reaction was subsequently heated to 75 °C and stirred for 2 hours. Upon completion, the reaction was cooled to room temperature and quenched with 50 mL of methanol. The crude mixture was azeotroped via rotary evaporation with toluene (3 x 50 mL) to remove DMF. The crude product was purified via reverse phase flash column chromatography ($H_2O$ to acetonitrile) to yield a 1:3.5 mixture of isomers **2A** and **2B**, respectively, as a yellow solid (2.9 g, 66%).

**$^1$H NMR** (500 MHz, Methanol-d4) δ **Major (2B)**: 8.29 (s, 3.5H), 8.27 (s, 3.5H), 7.89 (dd, *J* = 8.7, 1.8 Hz, 3.5H), 7.84 (d, *J* = 8.6 Hz, 3.5H), 7.38 (s, 3.5H), 3.93 (s, 10.5H), 3.89 (s, 10.5H). **Minor (2A)**: 8.31 (s, 1H), 8.27 (s, 1H), 8.18 (dd, *J* = 7.3, 1.3 Hz, 1H), 7.93 (d, *J* = 8.2 Hz, 1H), 7.37 – 7.34 (m, 1H), 4.00 (s, 3H), 3.93 (s, 3H).

**$^{13}$C NMR** (125 MHz, Methanol-d4) δ **Major (2B)**: 166.79, 156.64, 136.86, 132.55, 126.80, 126.25, 125.85, 125.24, 122.79, 122.65, 106.59, 55.08, 51.45. **Minor (2A)**: 166.82, 155.82, 140.64, 139.18, 132.18, 131.80, 131.72, 128.29, 127.76, 121.88, 106.06, 55.01, 51.41.

**HRMS** (ESI): calc. for $C_{13}H_{13}O_6S$ $[M+H]^+$: 297.0432, found: 297.0421.

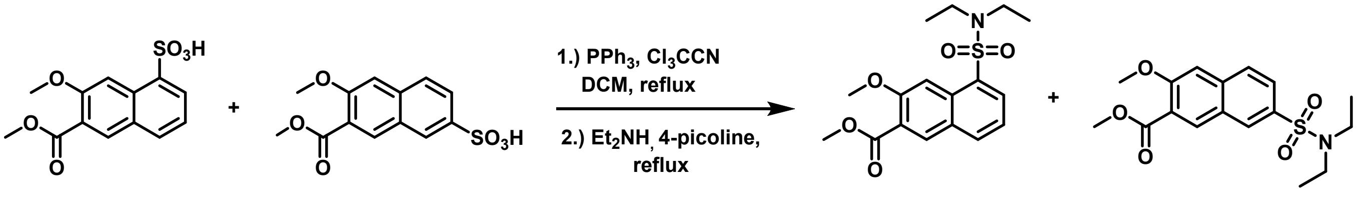


2A 2B 3 (15%) 4 (43%)

**Scheme 4. Synthesis of compounds 3 and 4.**

**Procedure:** In an oven dried, three neck, 25 mL round bottom flask, a mixture of **2A** and **2B** (250 mg, 0.75 mmol, 1.0 eq.) was dissolved in DCM (2.5 mL). A reflux condenser was attached, and the entire apparatus was purged with nitrogen. $Cl_3CCN$ (250 µL, 2.53 mmol, 3.0 eq.) was added dropwise at room temperature. The reaction was then heated to 45 °C and stirred for 10 minutes before addition of $PPh_3$ (660 mg, 2.53 mmol, 3.0 eq.) in 2.5 mL of DCM. The reaction was then stirred at reflux for an additional 5 hours until consumption of starting material was observed by TLC. $Et_2NH$ (262 µL, 2.53 mmol, 3.0 eq.) and freshly distilled 4-picoline (760 µL, 7.6 mmol, 9.0 eq.) were added dropwise as a mixture to the reaction at reflux. After 2 hours, the reaction was diluted with DCM (100 mL) and washed with 1M HCl (50 mL, 5x). The organic layer was washed with saturated sodium bicarbonate and brine, then dried over sodium sulfate, and concentrated *in vacuo*. The crude product was purified by column chromatography (hexanes to 3:1 hexanes/ethyl acetate) to yield the isolated products **3** (48 mg, 15%) as a light brown solid and **4** (137 mg, 43%) as a light brown solid.

**methyl 5-(*N*,*N*-diethylsulfamoyl)-3-methoxy-2-naphthoate (3)**

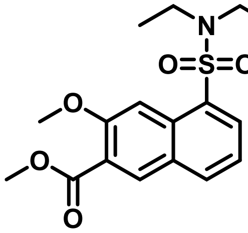


**$^1$H NMR** (500 MHz, $CDCl_3$) δ 8.30 (s, 1H), 8.19 (dd, $J$ = 7.4, 1.3 Hz, 1H), 8.05 (s, 1H), 7.97 (d, $J$ = 8.1 Hz, 1H), 7.38 (t, $J$ = 7.8 Hz, 1H), 4.01 (s, 3H), 3.93 (s, 3H), 3.33 (q, $J$ = 7.2 Hz, 4H), 1.04 (t, $J$ = 7.1 Hz, 6H).

**$^{13}$C NMR** (125 MHz, $CDCl_3$) δ 166.09, 156.86, 134.49, 133.79, 133.23, 131.77, 131.64, 128.55, 122.71, 122.61, 104.85, 56.08, 52.47, 40.98, 13.78.

**HRMS** (ESI): calc. for $C_{17}H_{22}NO_5S$ $[M+H]^+$: 352.1218, found: 352.1227.

**methyl 7-(*N*,*N*-diethylsulfamoyl)-3-methoxy-2-naphthoate (4)**

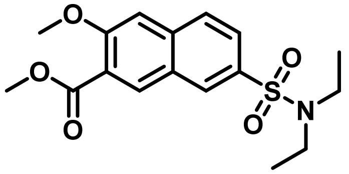

**$^1$H NMR** (500 MHz, Methanol-d$_4$) δ 8.42 (s, 1H), 8.41 (d, *J* = 1.8 Hz, 1H), 8.00 (d, *J* = 8.7 Hz, 1H), 7.85 (dd, *J* = 8.7, 1.9 Hz, 1H), 7.51 (s, 1H), 4.02 (s, 3H), 3.95 (s, 3H), 3.32 (m, 4H), 1.14 (t, *J* = 7.1 Hz, 6H).

**$^{13}$C NMR** (125 MHz, Methanol-d$_4$) δ 168.00, 158.90, 139.13, 137.24, 134.20, 129.87, 129.15, 127.63, 126.05, 125.13, 108.00, 56.59, 52.94, 43.42, 14.69.

**HRMS** (ESI): calc. for $C_{17}H_{22}NO_5S$ $[M+H]^+$: 352.1218, found: 352.1216.

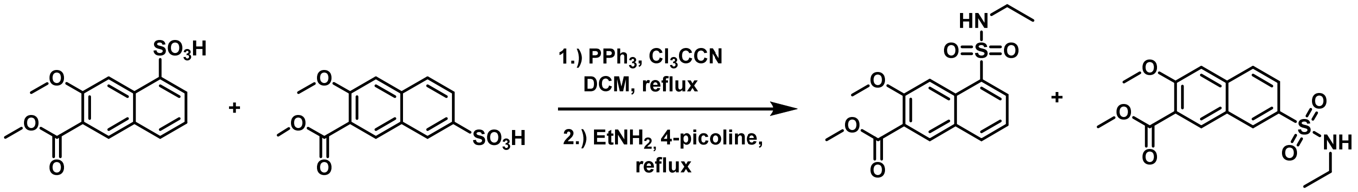


**2A** **2B** **5 (9%)** **6 (54%)**

**Scheme 5. Synthesis of compounds 5 and 6.**

**Procedure:** In an oven dried, three neck, 25 mL round bottom flask, a mixture of **2A** and **2B** (330 mg, 0.99 mmol, 1.0 eq.) was dissolved in DCM (3 mL). A reflux condenser was attached, and the entire apparatus was purged with nitrogen. $Cl_3CCN$ (330 µL, 2.97 mmol, 3.0 eq.) was added dropwise at room temperature. The reaction was then heated to 45 °C and stirred for 10 minutes before addition of $PPh_3$ (870 mg, 3.34 mmol, 3.0 eq.) in 3 mL of DCM. The reaction was then stirred at reflux for an additional 5 hours until consumption of starting material observed by TLC. $EtNH_2$ (346 µL, 2.97 mmol, 3.0 eq.) and freshly distilled 4-picoline (1 mL, 10.0 mmol, 9.0 eq.) were added dropwise as a mixture to the reaction at reflux. After 2 hours, the reaction was diluted with DCM (100 mL) and washed with 1M HCl (50 mL, 5x). The organic layer was washed with saturated sodium bicarbonate and brine, then dried over sodium sulfate, and concentrated *in vacuo*. The crude product was purified by column chromatography (hexanes to 3:1 hexanes/ethyl acetate) to yield the isolated products **5** (29 mg, 9%) as a white solid and **6** (173 mg, 54%) as a white solid.

**methyl 5-(*N*-ethylsulfamoyl)-3-methoxy-2-naphthoate (5)**

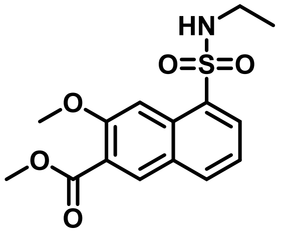

**$^1$H NMR** (500 MHz, DMSO-$d_6$) δ 8.42 (s, 1H), 8.26 (d, *J* = 8.1 Hz, 1H), 8.18 (dd, *J* = 7.3, 1.3 Hz, 1H), 8.06 (s, 1H), 7.96 (t, *J* = 5.7 Hz, 1H), 7.55 (t, *J* = 7.8 Hz, 1H), 3.99 (s, 3H), 3.87 (s, 3H), 2.80 (qd, *J* = 7.2, 5.5 Hz, 2H), 0.90 (t, *J* = 7.2 Hz, 3H).

**$^{13}$C NMR** (125 MHz, DMSO-$d_6$) δ 165.70, 155.77, 134.27, 134.01, 132.20, 130.84, 130.31, 128.08, 123.06, 122.93, 104.38, 55.99, 52.36, 37.33, 15.00.

**HRMS** (ESI) calc. for $C_{15}H_{18}NO_5S$ $[M+H]^+$: 324.0905, found: 324.0908.

**methyl 7-(*N*-ethylsulfamoyl)-3-methoxy-2-naphthoate (6)**

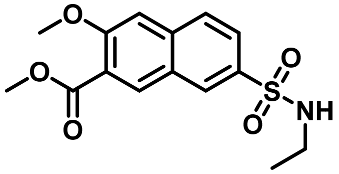

**$^1$H NMR** (500 MHz, $CDCl_3$) δ 8.37 (d, *J* = 1.8 Hz, 1H), 8.33 (s, 1H), 7.87 (dd, *J* = 8.7, 1.8 Hz, 1H), 7.81 (d, *J* = 8.7 Hz, 1H), 7.23 (s, 1H), 4.01 (s, 3H), 3.95 (s, 3H), 3.02 (m, *J* = 6.9 Hz, 2H), 1.09 (t, *J* = 7.2 Hz, 3H). (N-H proton exchanges and is therefore not visible on $^1$H spectrum).

**$^{13}$C NMR** (125MHz, $CDCl_3$) δ 166.12, 157.86, 137.75, 135.52, 133.68, 129.15, 127.98, 126.16, 124.99, 123.65, 106.84, 56.26, 52.62, 38.39, 15.17.

**HRMS** (ESI): calc. for $C_{15}H_{18}NO_5S$ $[M+H]^+$: 324.0905, found: 324.0908.

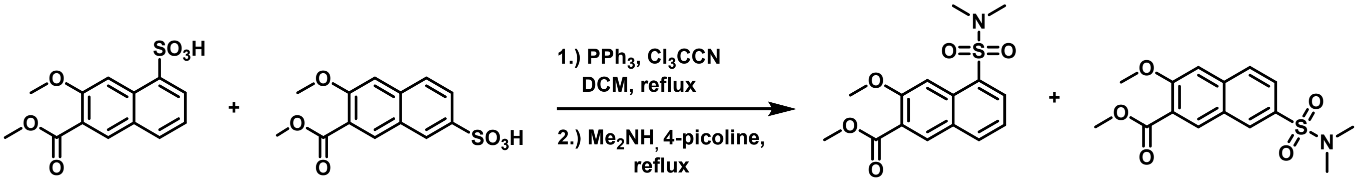


2A 2B 7 (9%) 8 (51%)

**Scheme 6. Synthesis of compounds 7 and 8.**

**Procedure:** In an oven dried, three neck, 25 mL round bottom flask, a mixture of **2A** and **2B** (330 mg, 0.99 mmol, 1.0 eq.) was dissolved in DCM (3 mL). A reflux condenser was attached, and the entire apparatus was purged with nitrogen. $Cl_3CCN$ (330 μL, 2.97 mmol, 3.0 eq.) was added dropwise at room temperature. The reaction was then heated to 45 °C and stirred for 10 minutes before addition of $PPh_3$ (870 mg, 3.34 mmol, 3.0 eq.) in 3 mL of DCM. The reaction was then stirred at reflux for an additional 5 hours until consumption of starting material was observed by TLC. $Me_2NH$ (356 μL, 2.97 mmol, 3.0 eq.) and freshly distilled 4-picoline (1 mL, 10.0 mmol, 9.0 eq.) were added dropwise as a mixture to the reaction at reflux. After 2 hours, the reaction was diluted with DCM (100 mL) and washed with 1M HCl (50 mL, 5x). The organic layer was washed with saturated sodium bicarbonate and brine, then dried over sodium sulfate, and concentrated *in vacuo*. The crude product was purified by column chromatography (hexanes to 3:1 hexanes/ethyl acetate) to yield the isolated products **7** (28 mg, 9%) as a white solid and **8** (166 mg, 51%) as a white solid.

**methyl 5-(*N*,*N*-dimethylsulfamoyl)-3-methoxy-2-naphthoate (7)**

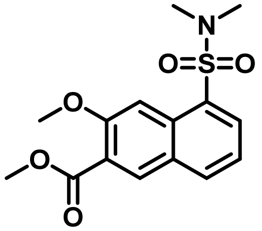

**$^1$H NMR** (500 MHz, $CDCl_3$) δ 8.34 (s, 1H), 8.25 (s, 1H), 8.22 (d, *J* = 7.4 Hz, 1H), 8.04 (d, *J* = 8.1 Hz, 1H), 7.46 (t, *J* = 7.7 Hz, 1H), 4.04 (s, 3H), 3.97 (s, 3H), 2.82 (s, 6H).

**$^{13}$C NMR** (125 MHz, $CDCl_3$) δ 166.23, 157.10, 134.95, 133.29, 132.71, 132.28, 131.18, 128.71, 122.87, 122.81, 105.22, 56.22, 52.65, 37.64.

**HRMS** (ESI): calc. for $C_{15}H_{18}NO_5S$ $[M+H]^+$: 324.0905, found: 324.0907.

**methyl 7-(*N*,*N*-dimethylsulfamoyl)-3-methoxy-2-naphthoate (8)**

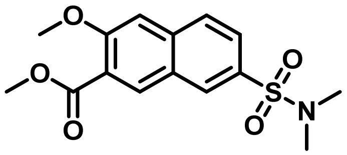

**$^1$H NMR** (500 MHz, $CDCl_3$) δ 8.36 (s, 1H), 8.27 (s, 1H), 7.84 (d, *J* = 8.7 Hz, 1H), 7.77 (dd, *J* = 8.6, 1.8 Hz, 1H), 7.25 (s, 1H), 4.01 (s, 3H), 3.94 (s, 3H), 2.72 (s, 6H).

**$^{13}$C NMR** (125 MHz, $CDCl_3$) δ 166.02, 157.83, 137.76, 133.66, 131.30, 129.71, 127.66, 126.22, 125.57, 123.73, 106.77, 56.25, 52.57, 38.03.

**HRMS** (ESI): calc. for $C_{15}H_{18}NO_5S$ $[M+H]^+$: 324.0905, found: 324.0898.

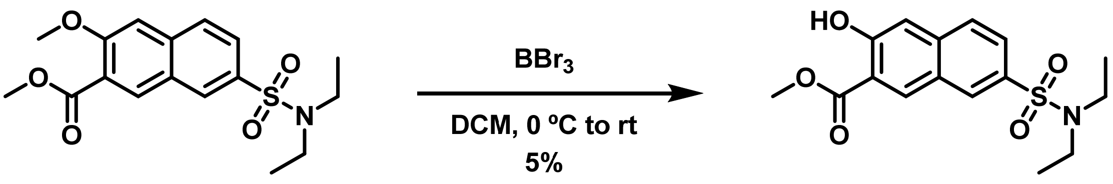


**4** **9**

Scheme 7. Synthesis of 9.

**methyl 7-(*N*,*N*-diethylsulfamoyl)-3-hydroxy-2-naphthoate (9)**

**Procedure**: To an oven dried 4 mL scintillation vial was added **4** (11.5 mg, 0.03 mmol, 1.0 eq.). The flask was purged with nitrogen, DCM (0.2 mL, 0.15 M) was added, and the reaction was cooled to 0 °C. $BBr_3$ (0.14 mL, 0.18 mmol, 3.3 eq.) was added dropwise as a 1M solution in DCM. The ice bath was removed, and the reaction was stirred at room temperature for 16 hours. The reaction mixture was then poured into ice water (15 mL) and extracted with ethyl acetate (15 mL, 3x). The combined ethyl acetate layers were then washed with brine, dried over sodium sulfate, and concentrated *in vacuo*. The crude product was purified with reverse phase flash column chromatography ($H_2O$ to acetonitrile) to yield **9** as a light brown solid (0.53 mg, 5%).

**$^1$H NMR** (500 MHz, $CDCl_3$) δ 10.68 (s, 1H), 8.61 (s, 1H), 8.35 (d, *J* = 1.2 Hz, 1H), 7.77 (s, 1H), 7.77 (s, 1H), 7.36 (s, 1H), 4.06 (s, 3H), 3.29 (q, *J* = 7.2 Hz, 4H), 1.14 (t, *J* = 7.1 Hz, 6H).

**$^{13}$C NMR** (125 MHz, $CDCl_3$) δ 169.97, 158.70, 139.18, 135.74, 133.96, 129.86, 127.81, 125.71, 125.66, 115.76, 112.25, 53.09, 42.13, 14.34.

**HRMS** (ESI): calc. for $C_{16}H_{20}NO_5S$ $[M+H]^+$: 338.1062, found: 338.1061.

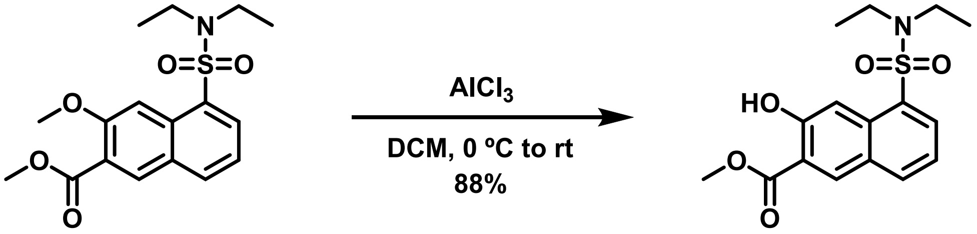


**3** **DSHN-OMe**

Scheme 8. Synthesis of DSHN-OMe.

**methyl 5-(*N*,*N*-diethylsuıramoyl)-3-hydroxy-2-naphthoate (DSHN-OMe)**

**Procedure**: To an oven dried 4 mL scintillation vial was added **3** (50 mg, 0.142 mmol, 1.0 eq.). The vial was purged with nitrogen and DCM (1 mL, 0.2 M) was added. The reaction was cooled to 0 °C and $AlCl_3$ (47 mg, 0.355mmol, 2.5 eq) was added in one portion. The ice bath was removed, and the reaction was stirred at room temperature for 2.5 hours until full consumption of starting material was observed by TLC. The reaction mixture was diluted with $H_2O$ and DCM (20 mL each). The organic layer was washed with saturated sodium bicarbonate and brine, then dried over sodium sulfate and concentrated *in vacuo*. The crude material was then recrystallized from minimal DCM and hexanes to afford **DSHN-OMe** as a light brown solid (43 mg, 88%).

**$^1$H NMR** (500 MHz, $CDCl_3$) δ 10.55 (s, 1H), 8.54 (s, 1H), 8.28 (dd, *J* = 7.3, 1.3 Hz, 1H), 8.06 (s, 1H), 7.99 (d, *J* = 8.2 Hz, 1H), 7.37 (dd, *J* = 8.2, 7.3 Hz, 1H), 4.05 (s, 3H), 3.38 (q, *J* = 7.1 Hz, 4H), 1.08 (t, *J* = 7.1 Hz, 6H).

**$^{13}$C NMR** (125 MHz, $CDCl_3$) δ 169.84, 157.98, 135.20, 133.99, 133.30, 133.28, 133.26, 128.11, 122.25, 115.01, 110.49, 53.00, 40.79, 13.72.

**HRMS** (ESI): calc. for $C_{16}H_{20}NO_5S$ $[M+H]^+$: 338.1062, found: 338.1059.

### $^{1}H$ and $^{13}C$ NMR Spectra

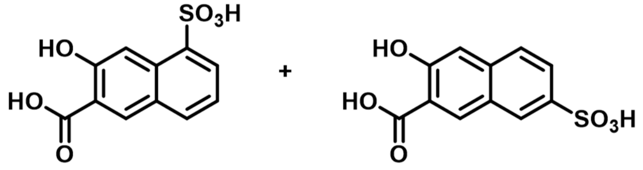


1A 1B

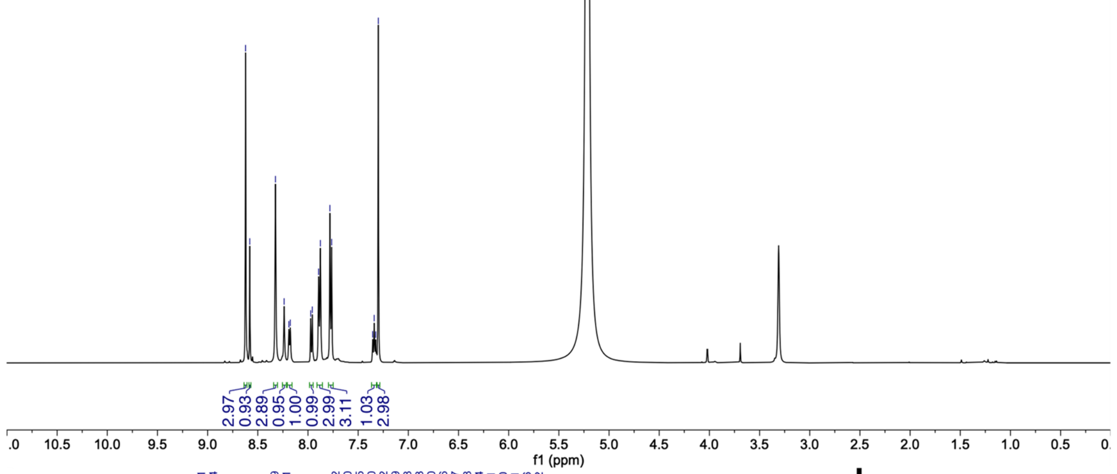


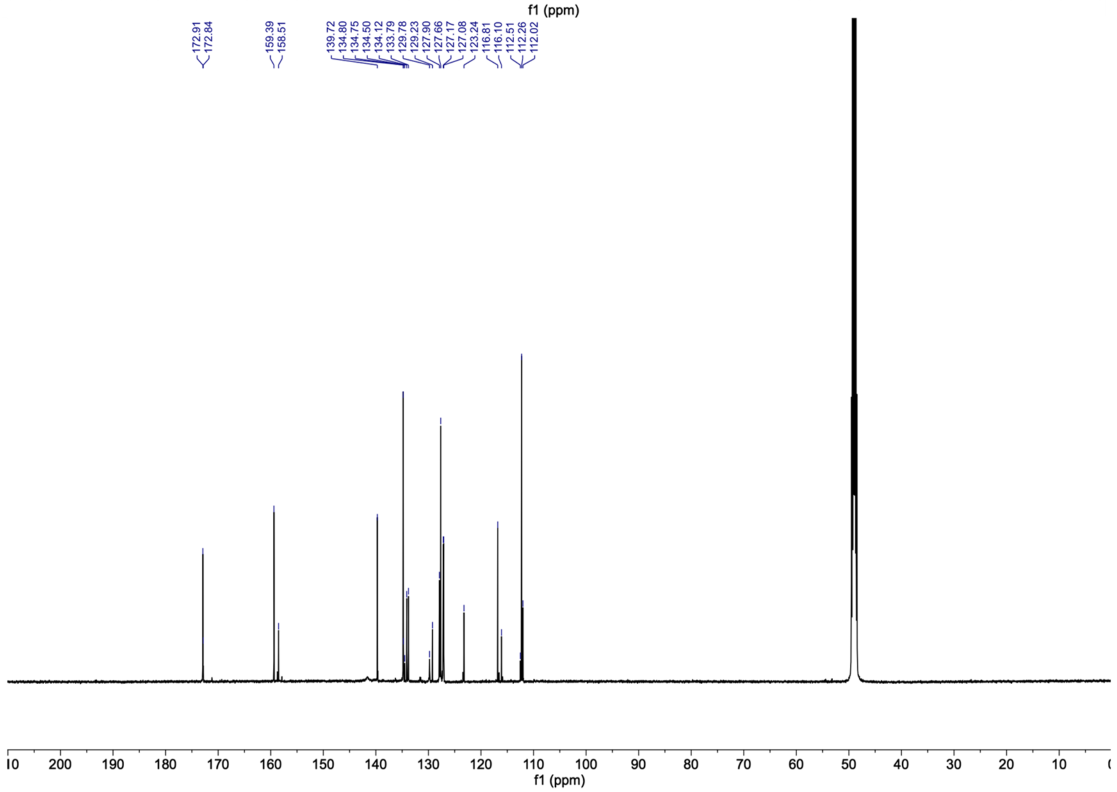

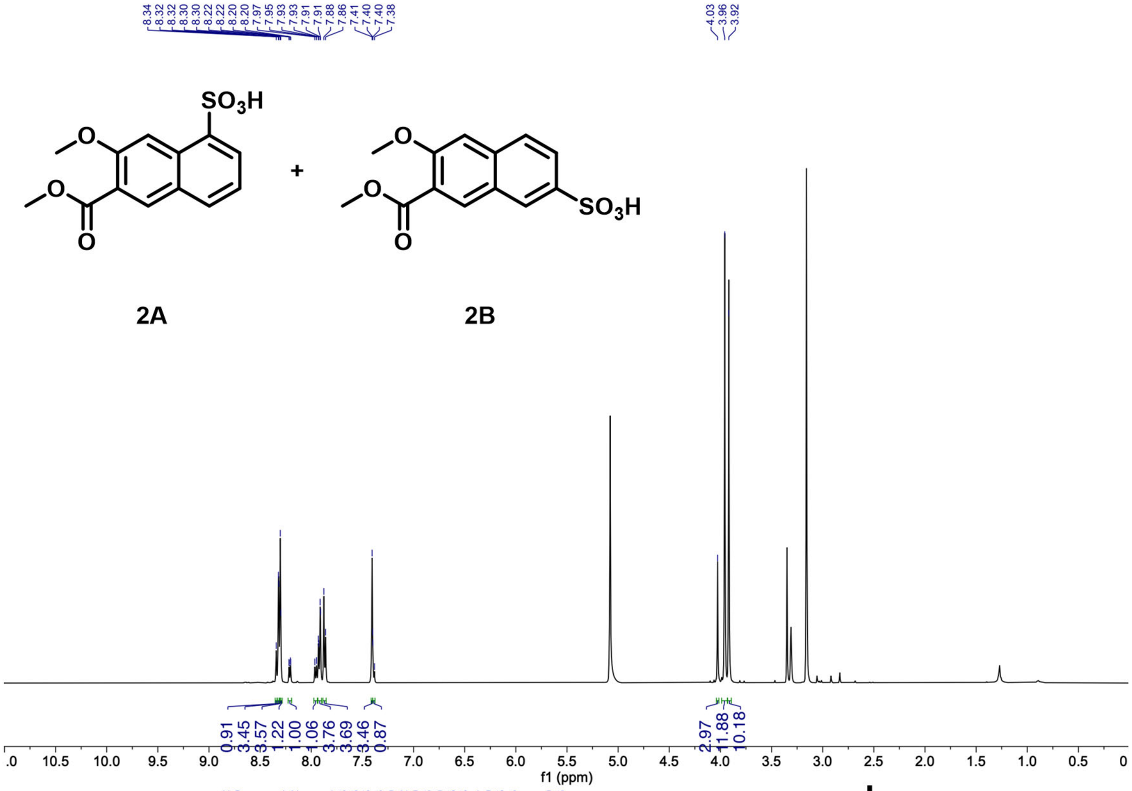
SO3H
O
O
O
+
SO3H
2A
2B
f1 (ppm)

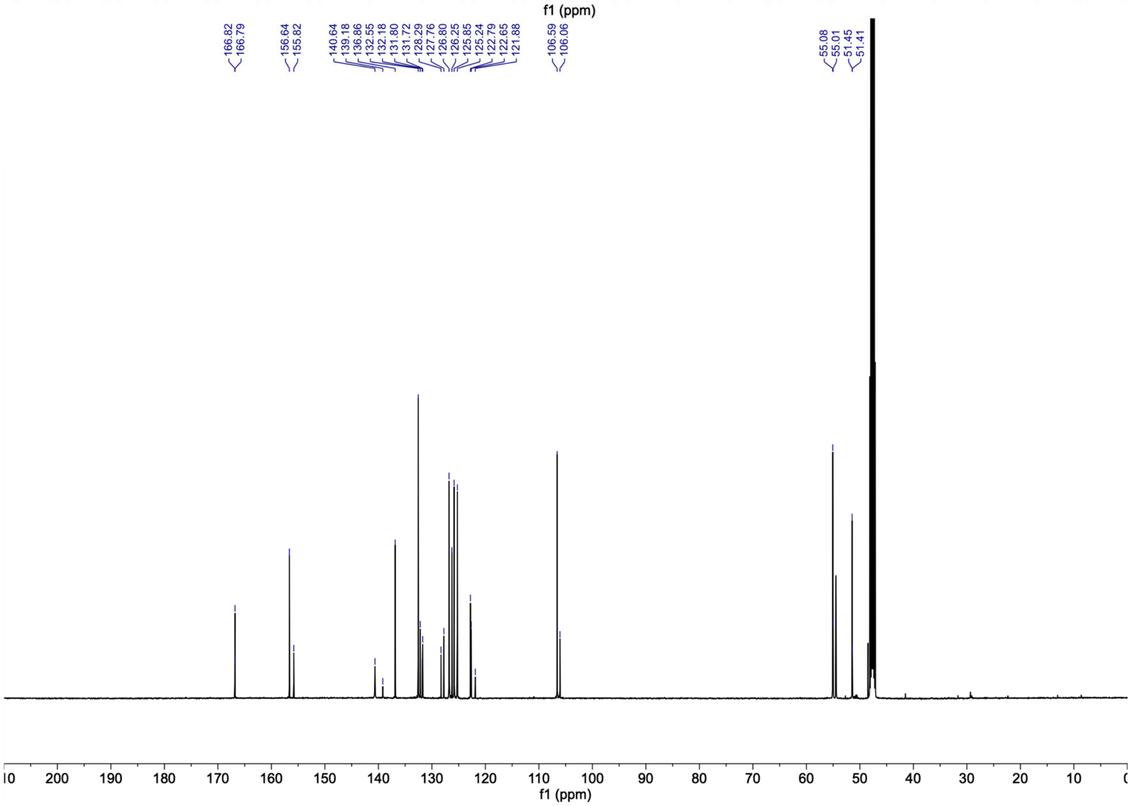
f1 (ppm)

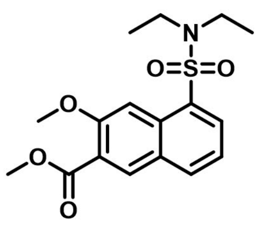
O
O
S
O
N
O
O

**3**

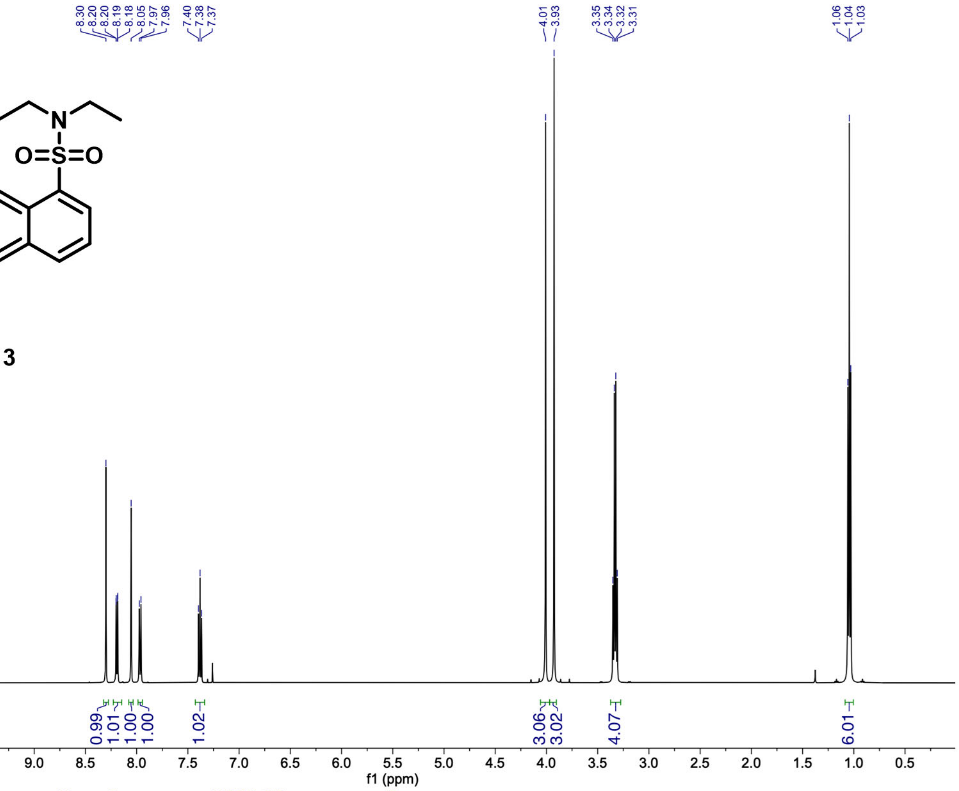
8.30
8.20
8.20
8.19
8.18
8.05
7.97
7.96
7.40
7.38
7.37
4.01
3.93
3.35
3.34
3.32
3.31
1.06
1.04
1.03
0.99
1.01
1.00
1.00
1.02
3.06
3.02
4.07
6.01
f1 (ppm)

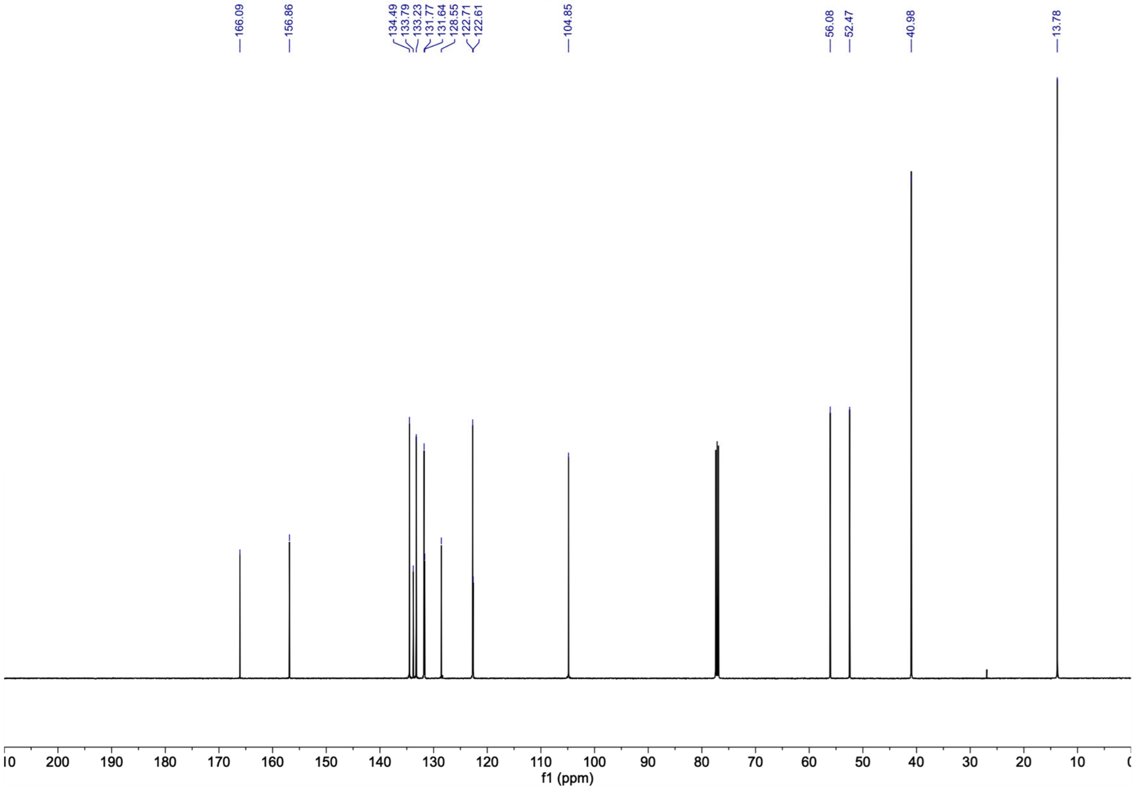
166.09
156.86
134.49
133.79
133.23
131.77
131.64
128.55
122.71
122.61
104.85
56.08
52.47
40.98
13.78
f1 (ppm)

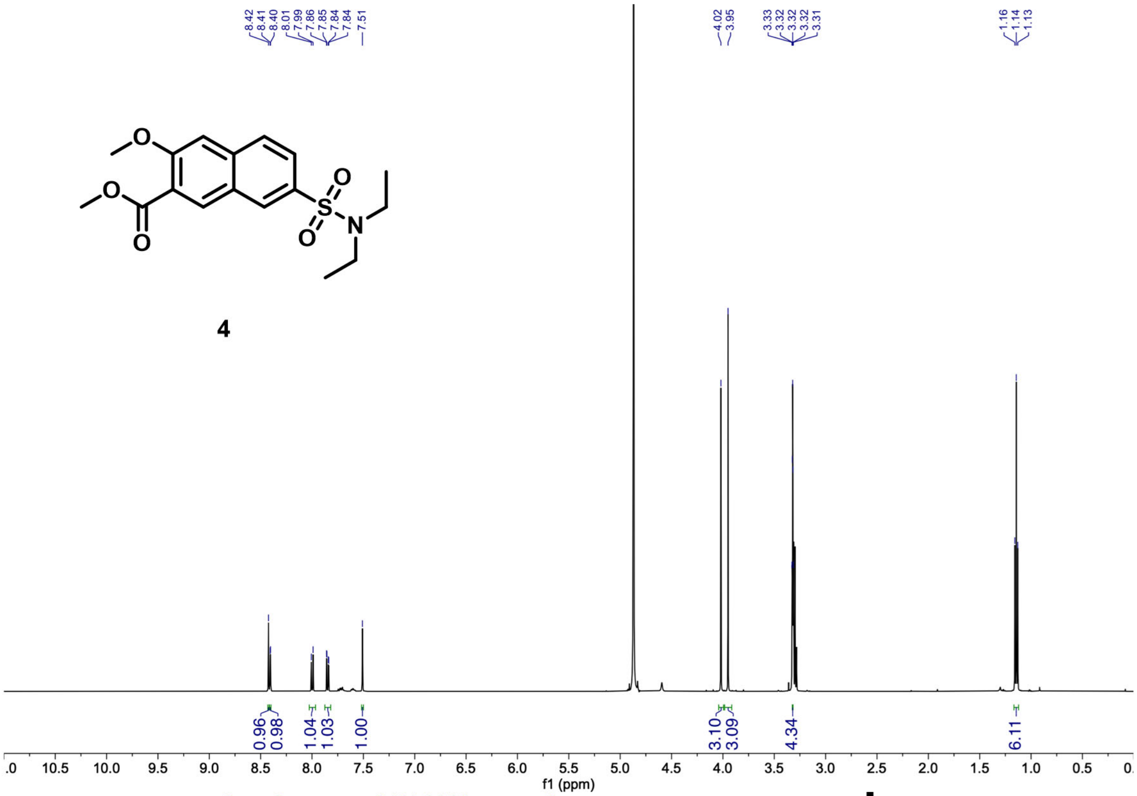

**4**

8.42
8.41
8.40
8.01
7.99
7.86
7.85
7.84
7.84
7.51
4.02
3.95
3.33
3.32
3.32
3.32
3.31
1.16
1.14
1.13
0.96
0.98
1.04
1.03
1.00
3.10
3.09
4.34
6.11
f1 (ppm)

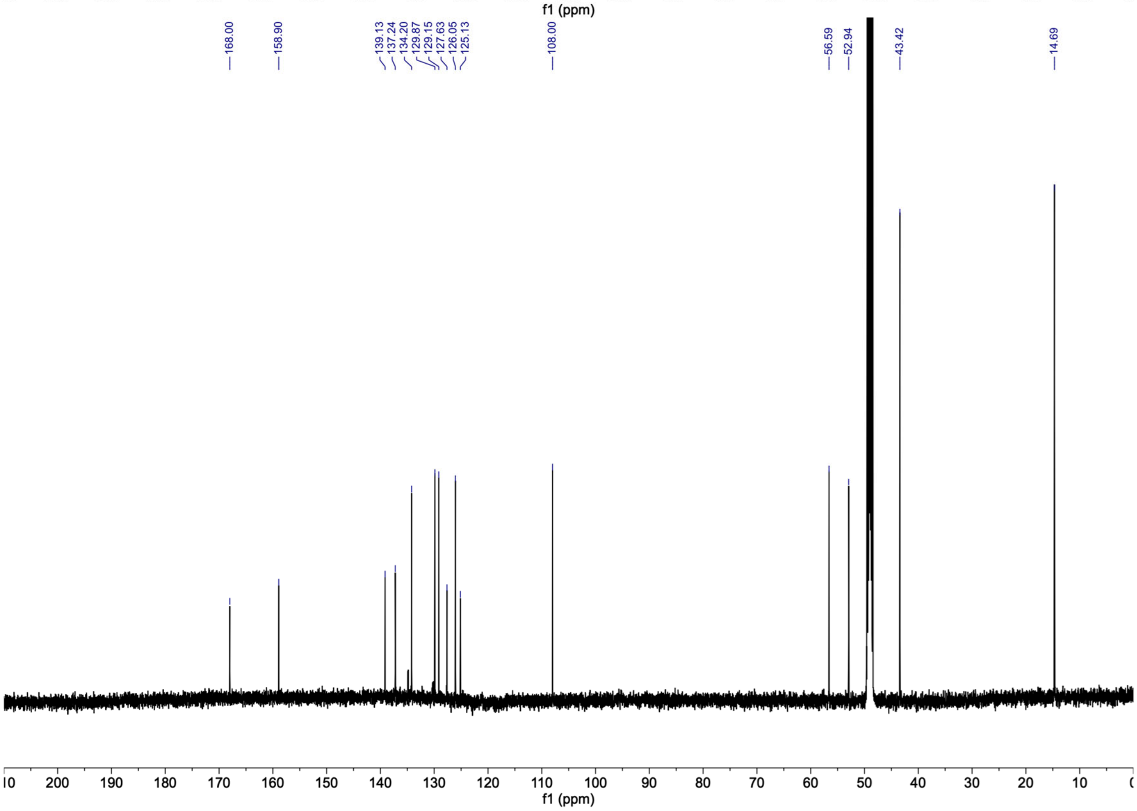
168.00
158.90
139.13
137.24
134.20
129.87
129.15
127.63
126.05
125.13
108.00
56.59
52.94
43.42
14.69
f1 (ppm)

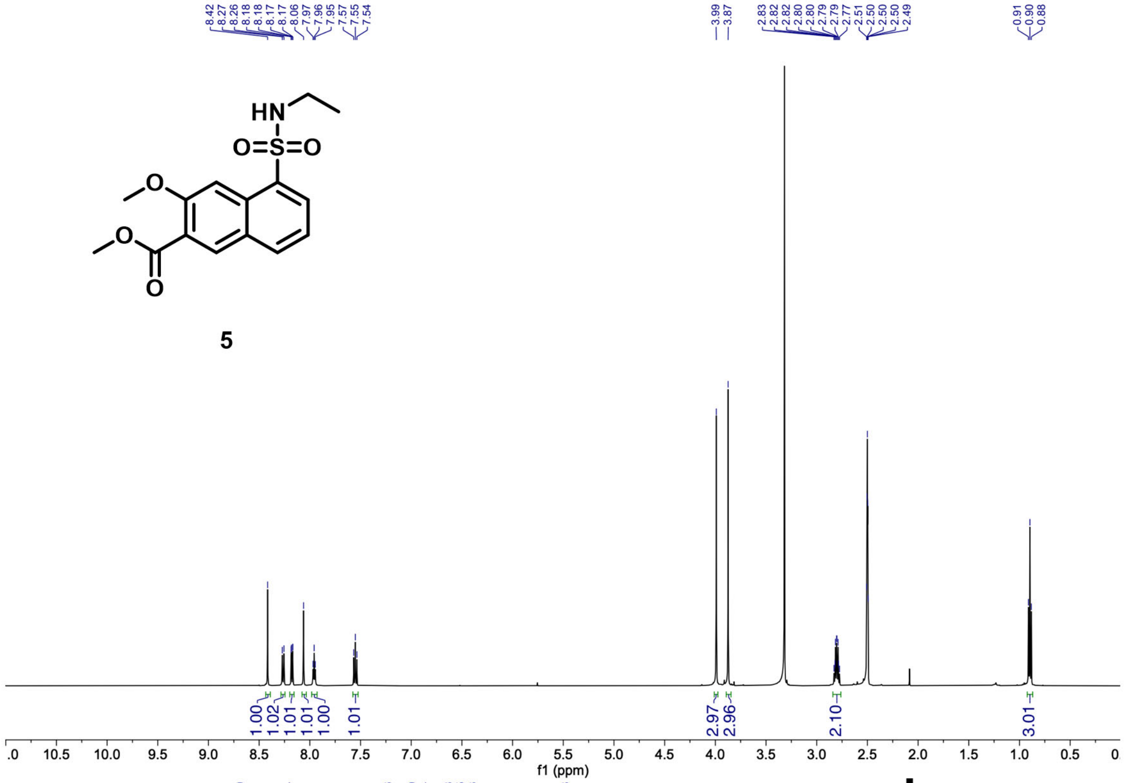
HN
O=S=O
O
O
O

**5**

8.42 8.27 8.26 8.18 8.18 8.17 8.17 8.06 7.97 7.96 7.95 7.57 7.55 7.54
3.99 3.87
2.83 2.82 2.82 2.80 2.80 2.79 2.77 2.51 2.50 2.50 2.50 2.49
0.91 0.90 0.88
1.00 1.02 1.01 1.01 1.00 1.01 2.97 2.96 2.10 3.01
f1 (ppm)

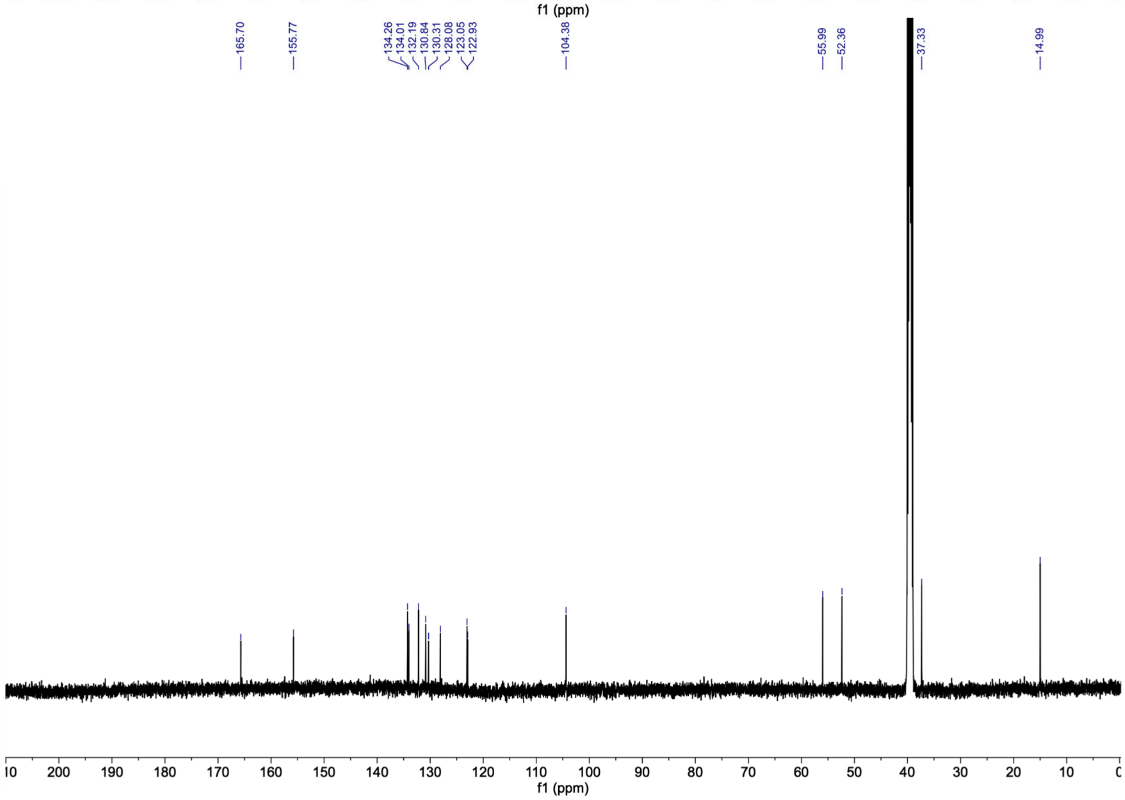
165.70
155.77
134.26 134.01 132.19 130.84 130.31 128.08 123.05 122.93
104.38
55.99 52.36
37.33
14.99
f1 (ppm)

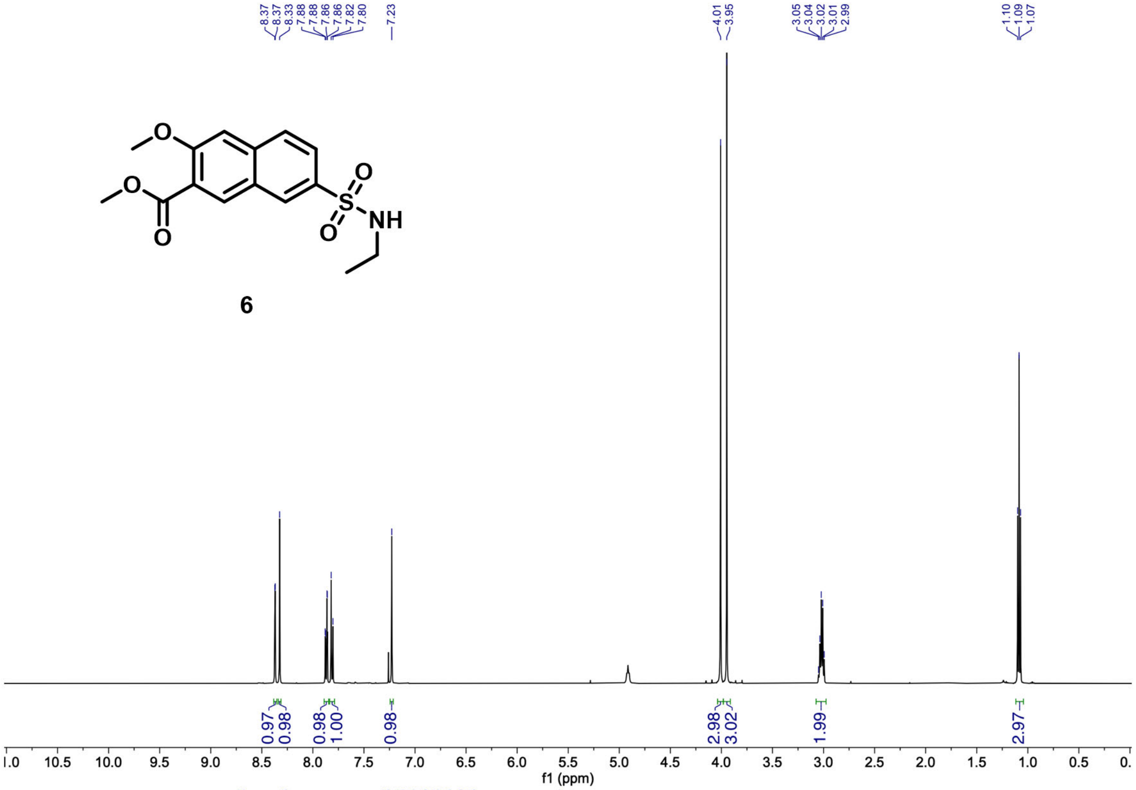

O
O
O
S
O
O
NH
6
8.37
8.37
8.33
7.88
7.88
7.86
7.86
7.82
7.80
7.23
4.01
3.95
3.05
3.04
3.02
3.01
2.99
1.10
1.09
1.07
0.97
0.98
0.98
1.00
0.98
2.98
3.02
1.99
2.97
f1 (ppm)


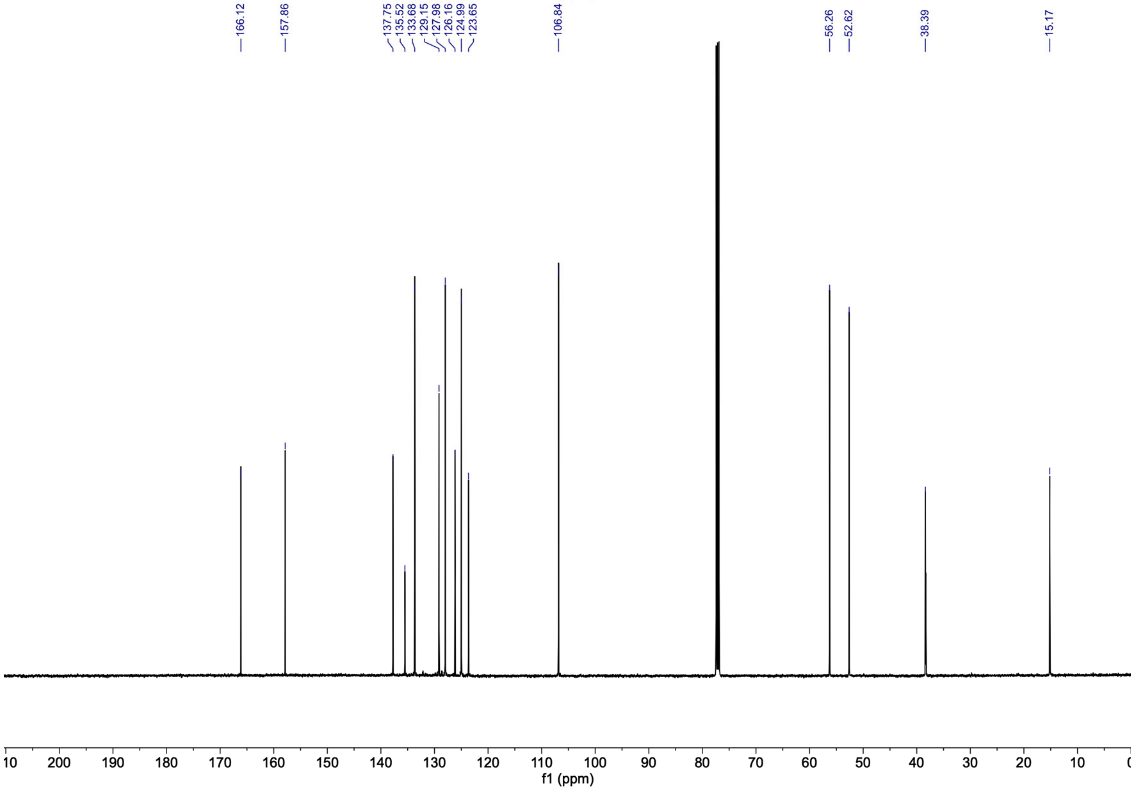

166.12
157.86
137.75
135.52
133.68
129.15
127.98
126.16
124.99
123.65
106.84
56.26
52.62
38.39
15.17
f1 (ppm)

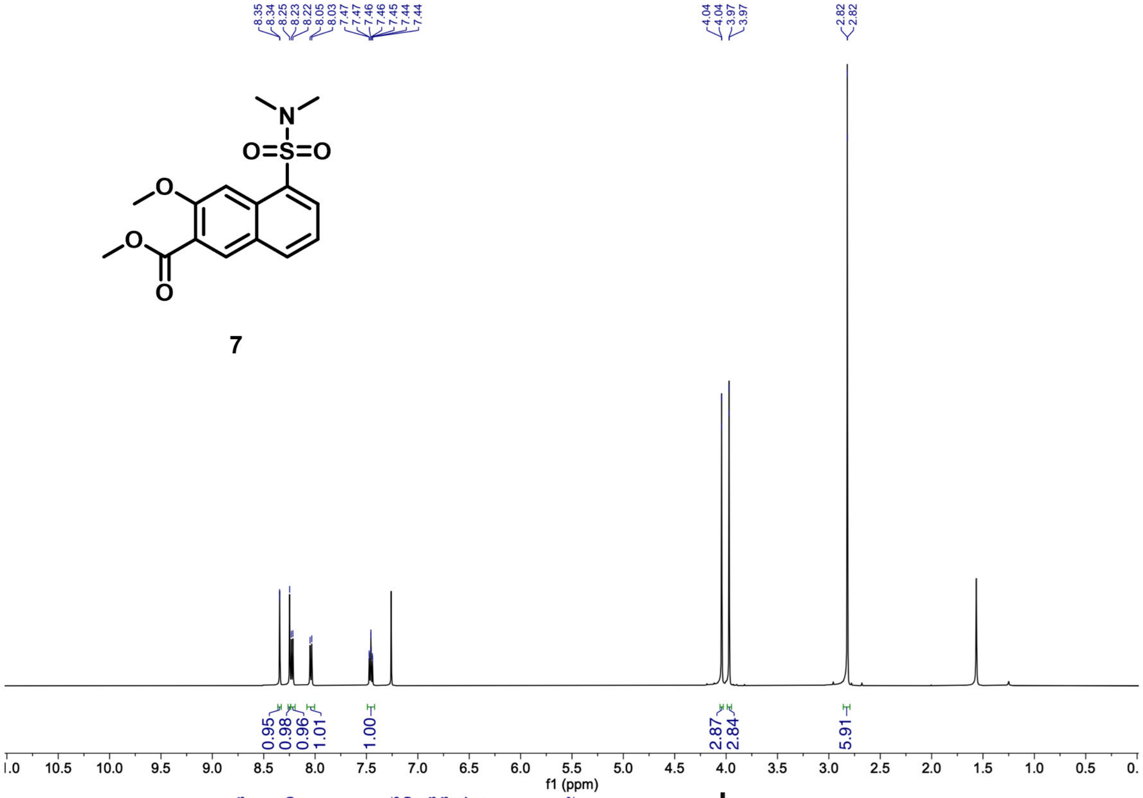
N
O=S=O
O
O
O
7
8.35 8.34 8.25 8.23 8.22 8.05 8.03 7.47 7.46 7.46 7.45 7.44 7.44
4.04 4.04 3.97 3.97
2.82 2.82
0.95 0.98 0.96 1.01 1.00
2.87 2.84
5.91
11.0 10.5 10.0 9.5 9.0 8.5 8.0 7.5 7.0 6.5 6.0 5.5 5.0 4.5 4.0 3.5 3.0 2.5 2.0 1.5 1.0 0.5 0.0
f1 (ppm)

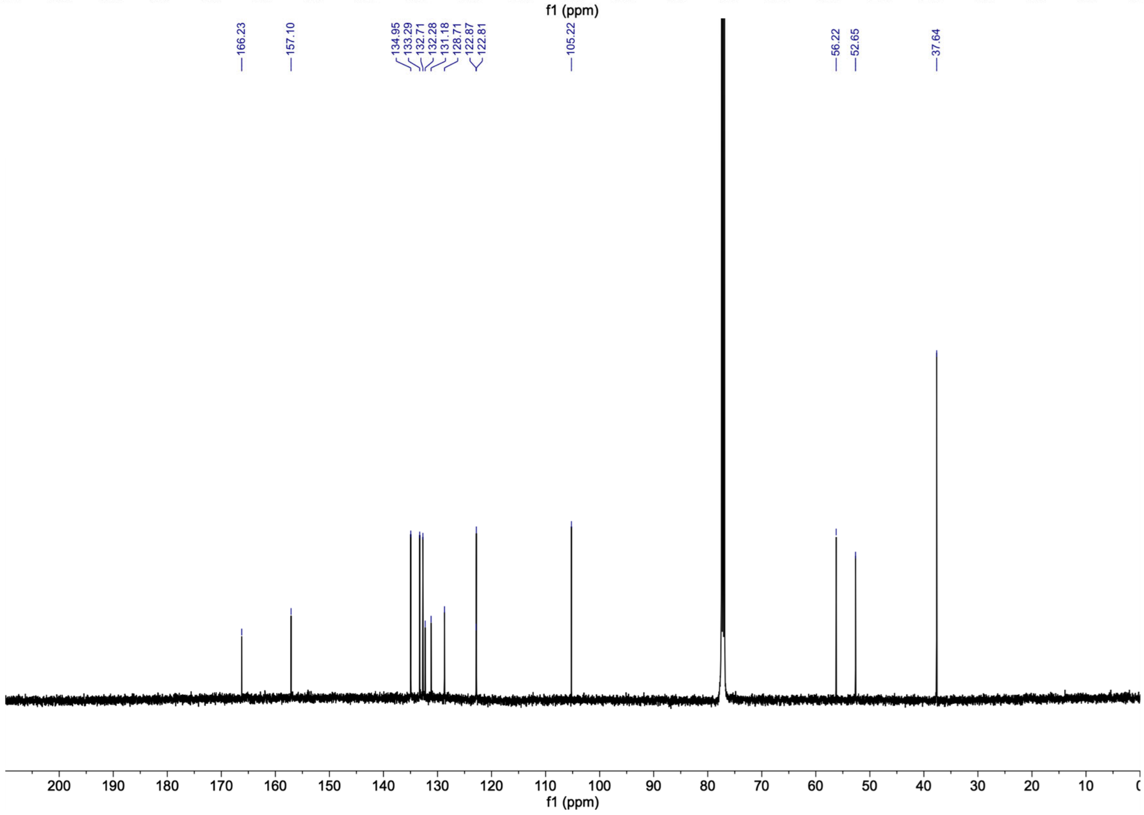
166.23
157.10
134.95 133.29 132.71 132.28 131.18 128.71 122.87 122.81
105.22
56.22 52.65
37.64
200 190 180 170 160 150 140 130 120 110 100 90 80 70 60 50 40 30 20 10 0
f1 (ppm)

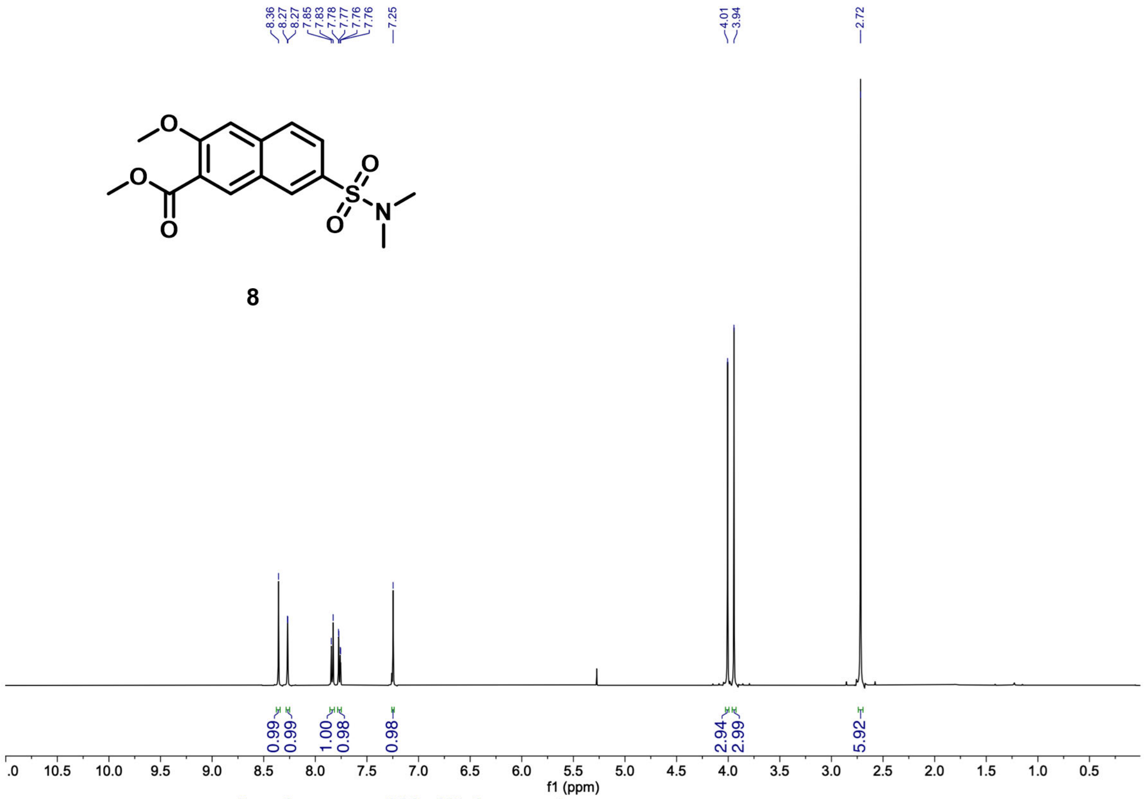

8
8.36
8.27
8.27
7.85
7.83
7.78
7.77
7.76
7.76
7.25
4.01
3.94
2.72
0.99
0.99
1.00
0.98
0.98
2.94
2.99
5.92
f1 (ppm)


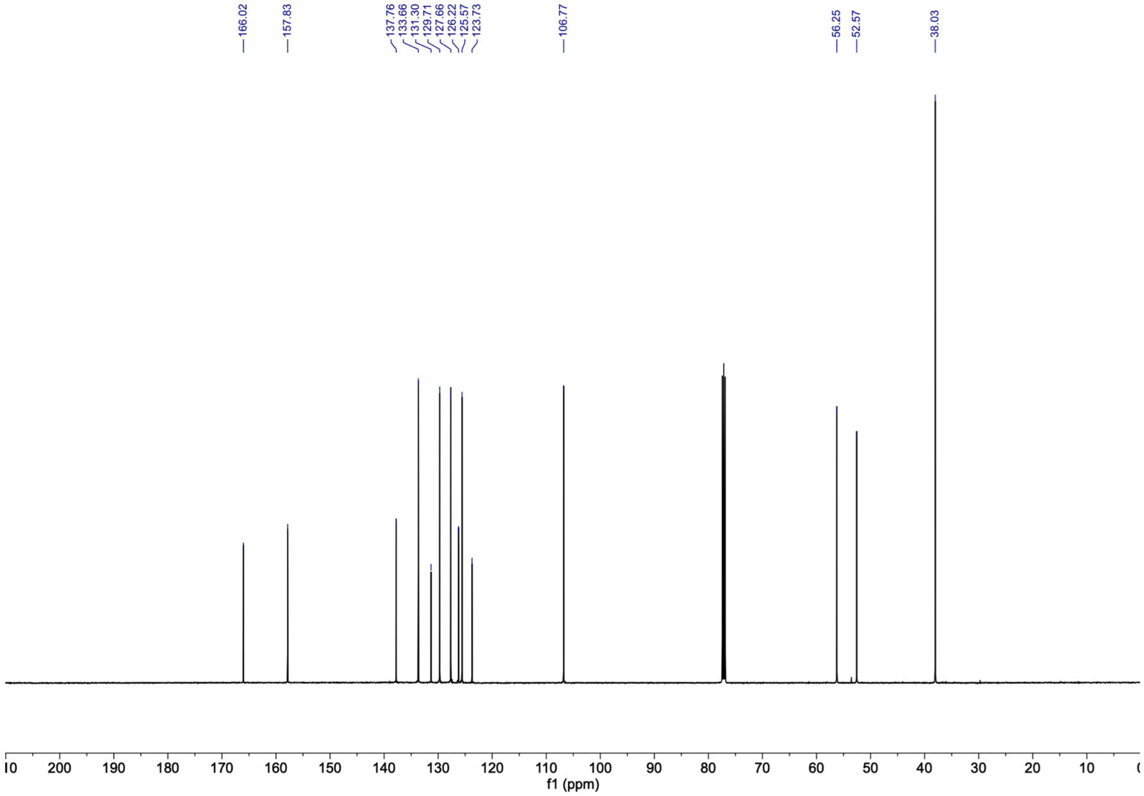

166.02
157.83
137.76
133.66
131.30
129.71
127.66
126.22
125.57
123.73
106.77
56.25
52.57
38.03
f1 (ppm)

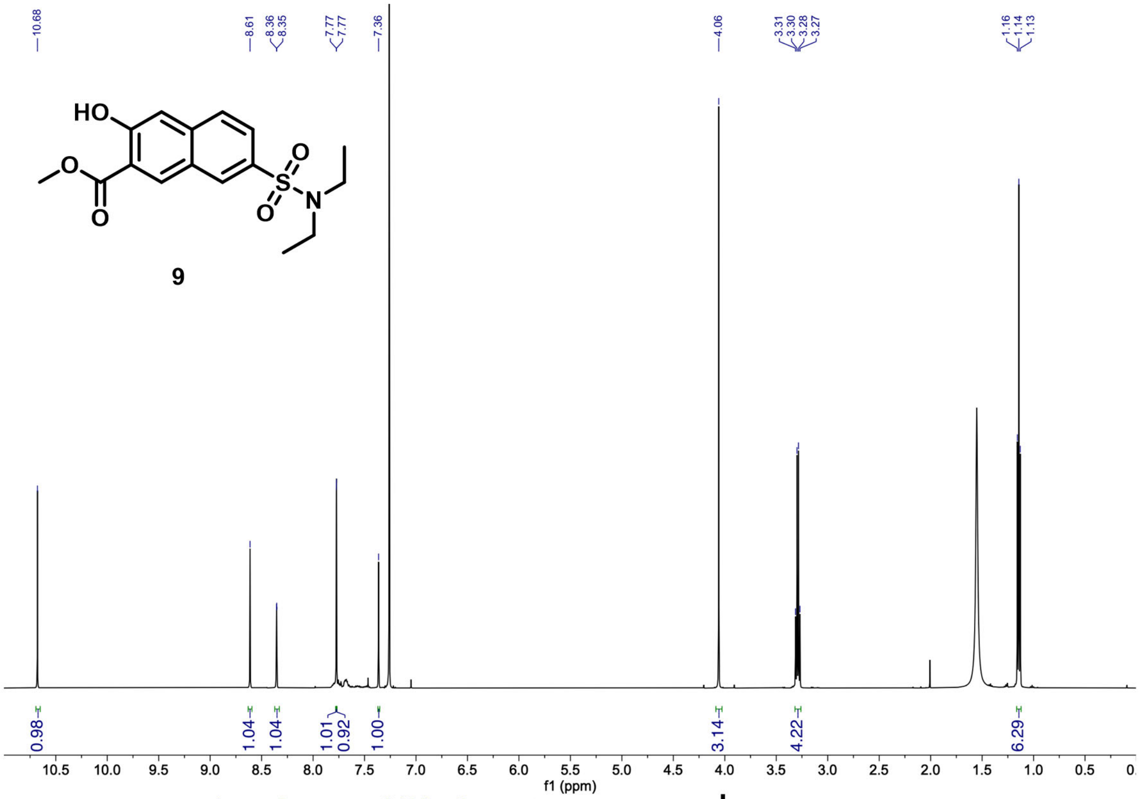

10.68
8.61
8.36
8.35
7.77
7.77
7.36
4.06
3.31
3.30
3.28
3.27
1.16
1.14
1.13
HO
O
O
S
O
O
N
9
0.98
1.04
1.04
1.01
0.92
1.00
3.14
4.22
6.29
f1 (ppm)


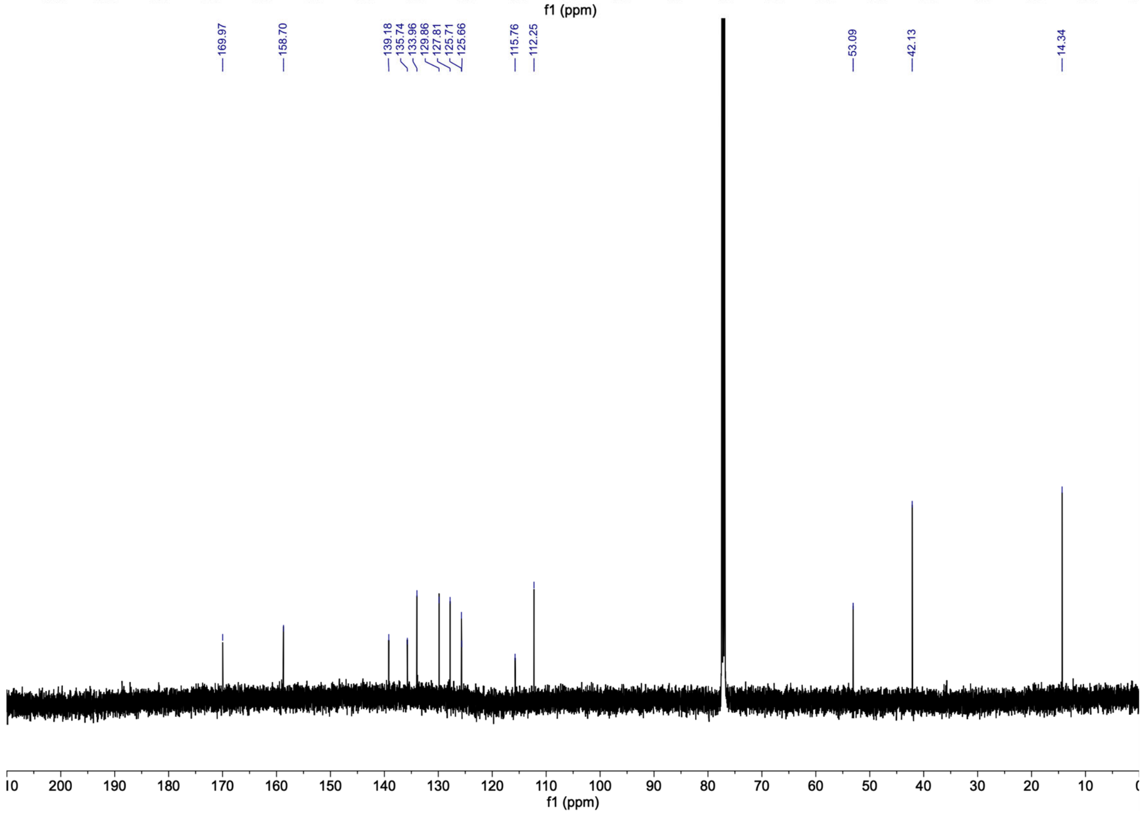

169.97
158.70
139.18
135.74
133.96
129.86
127.81
125.71
125.66
115.76
112.25
53.09
42.13
14.34
f1 (ppm)

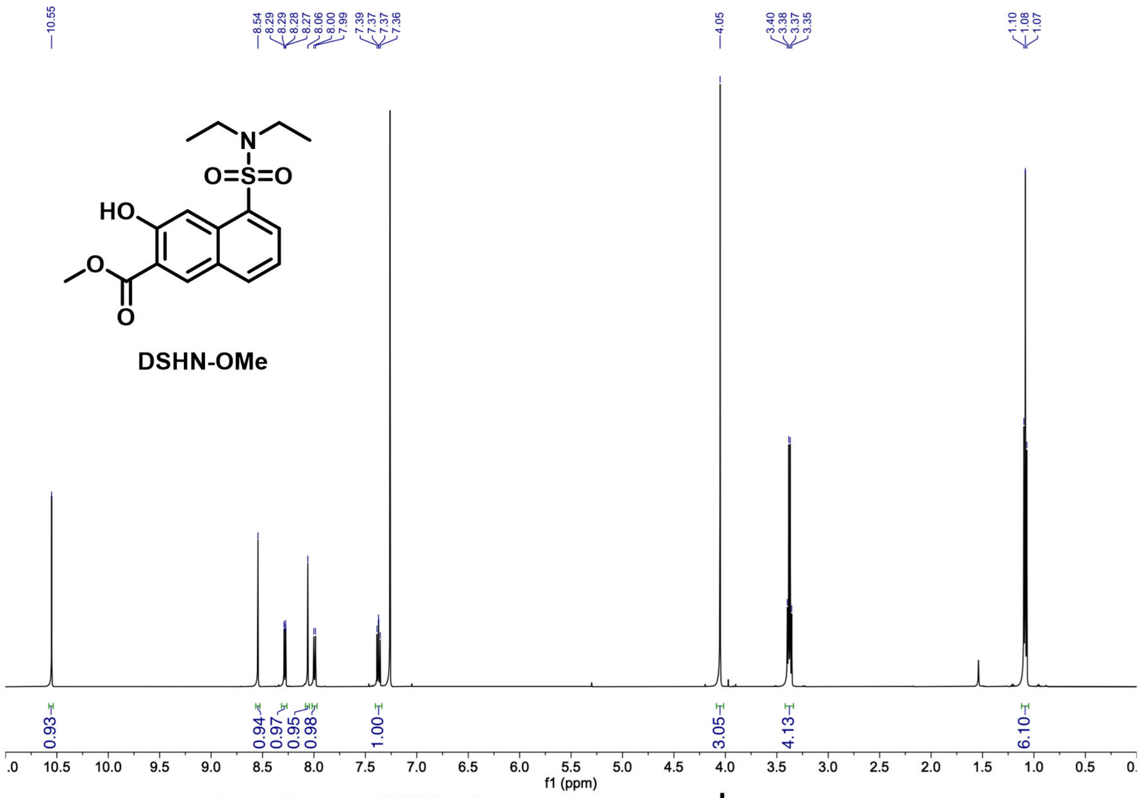
10.55
8.54
8.29
8.29
8.28
8.27
8.06
8.00
7.99
7.39
7.37
7.37
7.36
4.05
3.40
3.38
3.37
3.35
1.10
1.08
1.07
HO
O
N
S
O
O
O
DSHN-OMe
0.93
0.94
0.97
0.95
0.98
1.00
3.05
4.13
6.10
f1 (ppm)

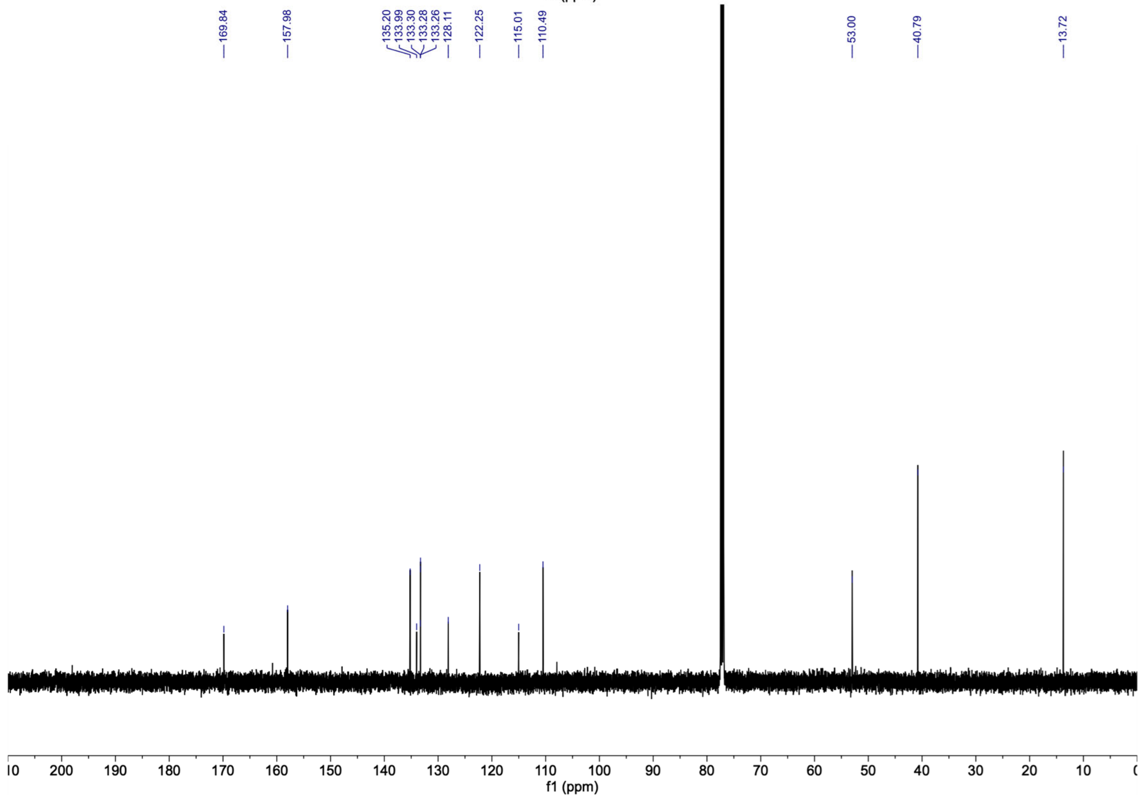
169.84
157.98
135.20
133.99
133.30
133.28
133.26
128.11
122.25
115.01
110.49
53.00
40.79
13.72
f1 (ppm)

### Supplemental Figures

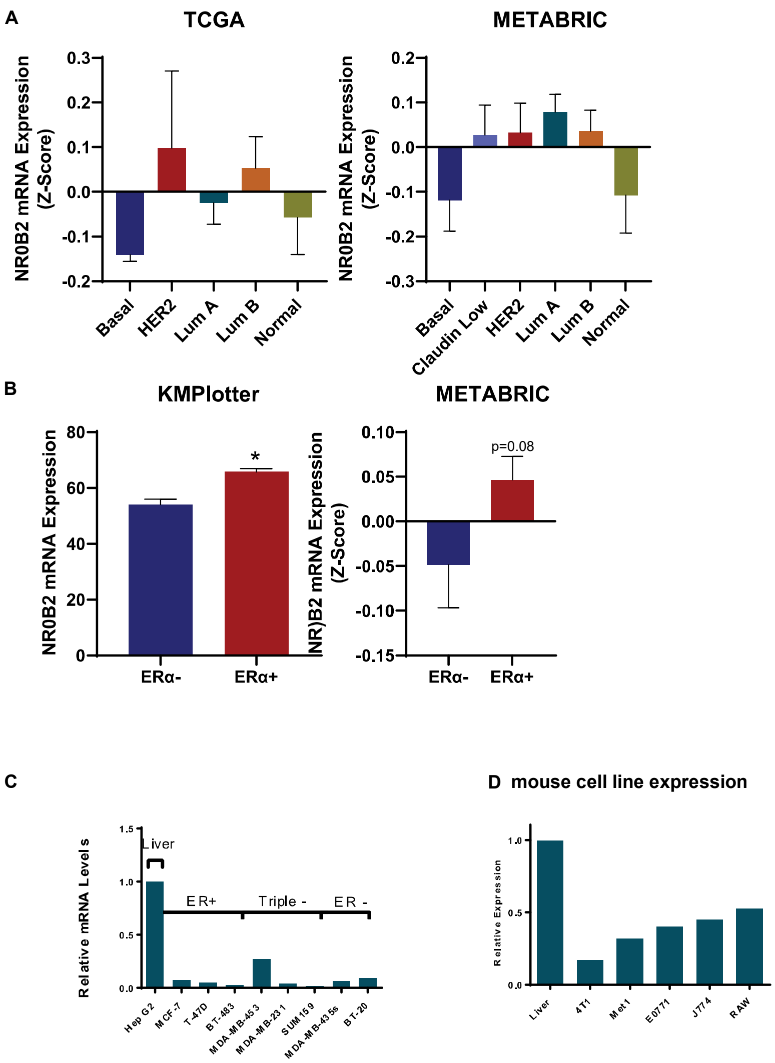


**Supplemental Figure 1 (SFig. 1): NR0B2 mRNA expression levels vary with breast tumor subtype, but are relatively low in breast cancer cell lines. (A)** Analysis of the TCGA (N=981) and METABRIC (N=1898) databases indicates that NR0B2 is expressed at different levels depending on the subtype. **(B)** Analysis of the compiled KMPlotter (N=4929) or METABRIC (N=1904) datasets indicates that NR0B2 has elevated expression in ERα+ disease compared to ERα- (t test) **(C)** NR0B2 is expressed at low levels in various human breast cancer cell lines compared to HEPG2, a liver cancer cell line. **(D)** NR0B2 is expressed at higher concentrations in models of myeloid cells compared to murine mammary cancer lines, but less than liver tissue.

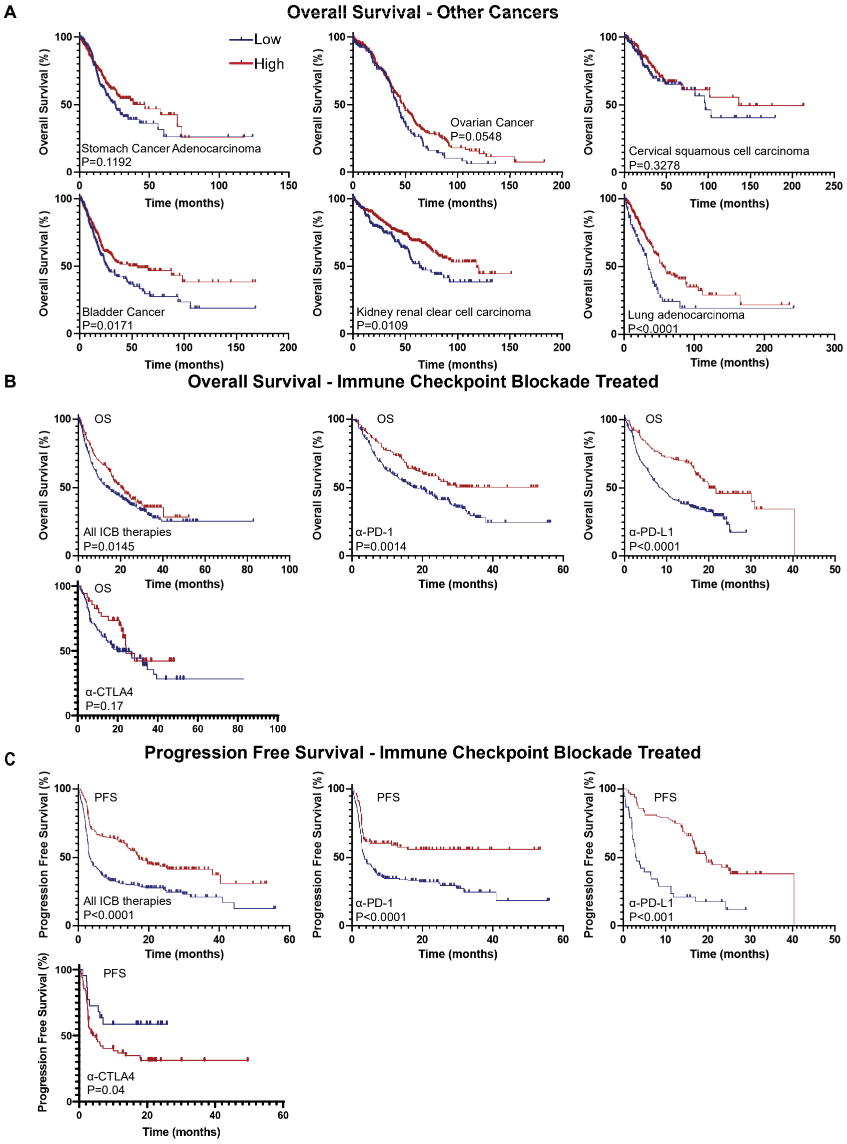


**Supplemental Figure 2 (SFig. 2): (A) NR0B2 mRNA expression within tumors is associated with an increased overall survival time in various cancers**. The Kaplan-Meier plotter was used to probe associations between NR0B2 expression and indicated cancers. The auto-cutoff method was used to parse patients between low and high expression. [stomach N=742, ovarian N=746, cervical N=608, bladder N=808, kidney N=1060, lung N=1008]. **(B)** NR0B2 mRNA expression within tumors from patients treated with immune checkpoint blockers (ICB) is associated with an increased overall survival time (OS). The Kaplan-Meier plotter was used to probe associations between NR0B2 expression in both genders and “all cancers” within the database [bladder N=90, esophageal adenocarcinoma N=103, glioblastoma N=28, hepatocellular carcinoma N=22, HNSCC N=110, melanoma N=570, NSCLC N=21, urothelial N=348]. The auto-cutoff method was used to parse patients between low and high expression. All ICB therapies were considered together or parsed into α-PD-1, α-PD-L1 or α-CTLA4 as indicated. **(C)** NR0B2 mRNA expression within tumors from patients treated with immune checkpoint blockers (ICB) is associated with an increased progression free survival time (PFS). The Kaplan-Meier plotter was used to probe associations between NR0B2 expression in both genders and all cancers within the database. P values were generated using the Log-rank (Mantel-Cox) method.

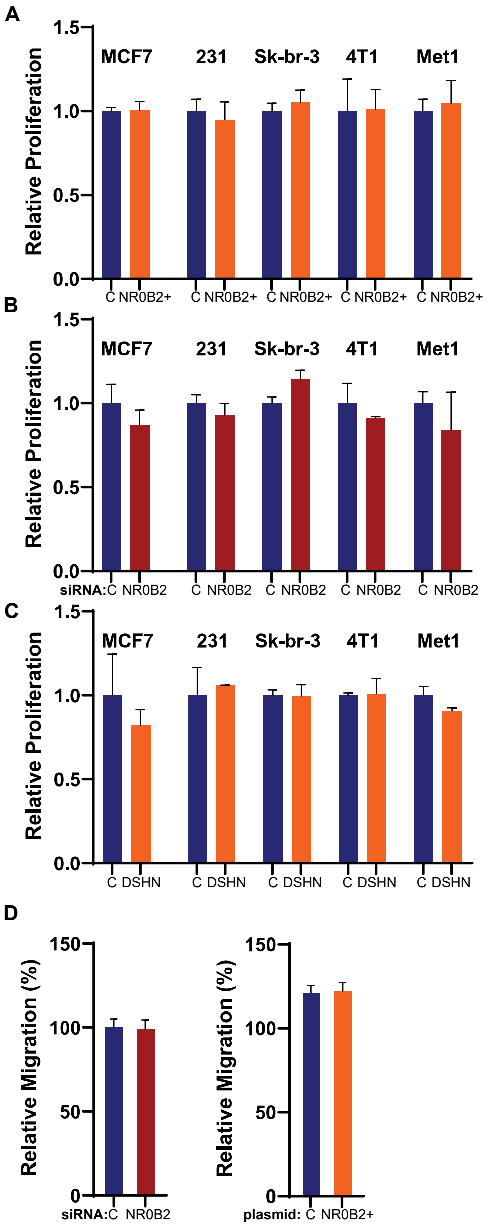


**Supplemental Figure 3 (SFig. 3): Modulation of NR0B2 within breast cancer cell lines does not impact proliferation. (A)** A control (C) or expression vector for NR0B2 was introduced to MCF7, MDA-MB-231 (231), Sk-br-3, 4T1 or Met1 mammary cancer cells and proliferation assessed (N=3/group). **(B)** Control (C) or siRNA against NR0B2 was introduced to indicated cell lines and proliferation assessed (N=3/group). **(C)** Indicated cell lines were treated with vehicle (V) or the NR0B2 agonist DSHN, and proliferation assessed (N=3/group). **(D)** MDA-MB-231 migration assays when NR0B2 is knocked down by siRNA or overexpressed. Wound healing assays measuring gap closure (N=5/group). Data is presented as mean ± SEM.

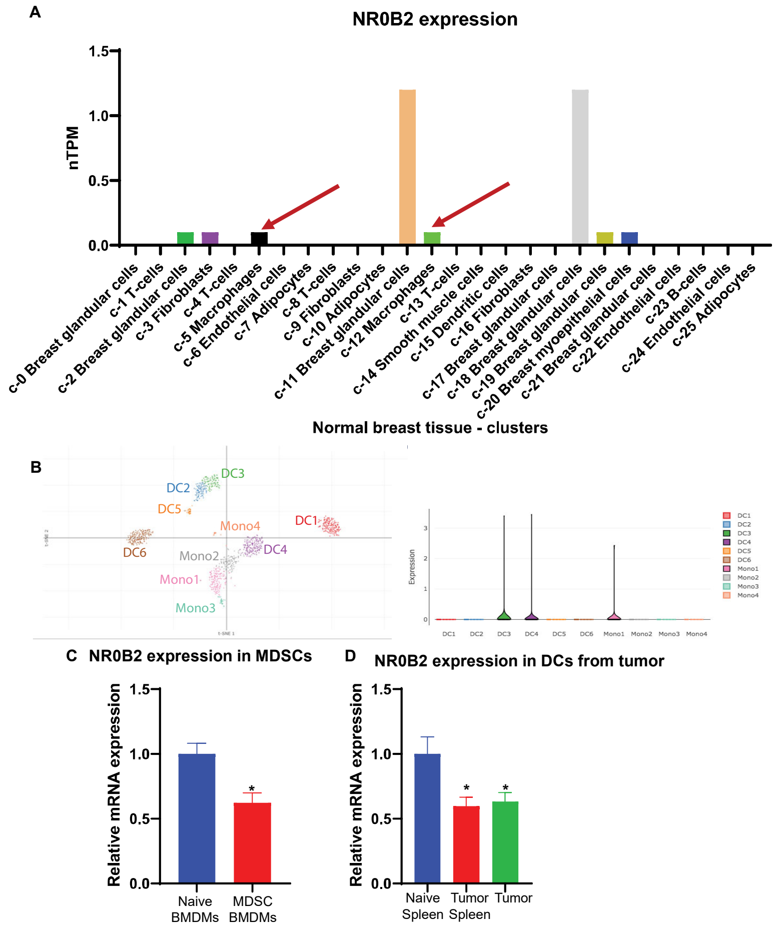


**Supplemental Figure 4 (SFig. 4): NR0B2 is expressed in different human myeloid populations. (A)** Within healthy breast tissue, NR0B2 is more highly expressed in breast glandular cells and macrophages. scRNA-seq data obtained from The Human Protein Atlas. **(B)** NR0B2 expression in different PBMC populations. scRNA-seq data obtained from the BROAD Institute Atlas of human blood dendritic cells and monocytes. **(C)** NR0B2 expression in naïve BMDMs and those differentiated into myeloid derived suppressor cells (MDSCs) (N=3/group, t test). **(D)** NR0B2 expression in DCs isolated from spleens of naïve mice, spleens in mice bearing 4T1 tumors or 4T1 tumors themselves (N=3/group, Šidák test).

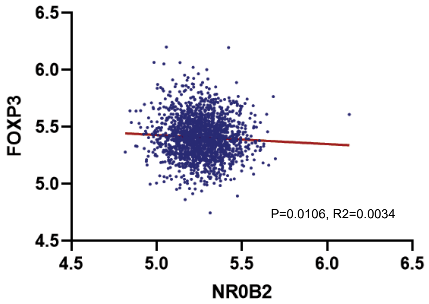


**Supplemental Figure 5 (SFig. 5): NR0B2 mRNA expression is inversely correlated with FoxP3 expression within human breast tumors.** Linear regression of expression data obtained from METABRIC. Slope of the line is significantly different than 0 (N=1904, P=0.0106).

### A - CD8 staining

Tumor: Distribution of Wilcoxon Score　Stroma: Distribution of Wilcoxon Score

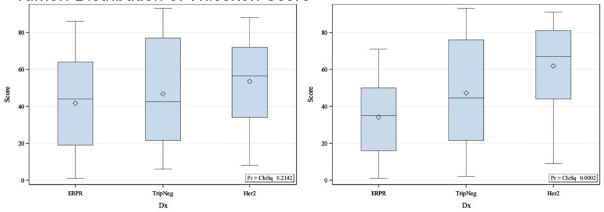


| Pairwise Two-Sided Multiple Comparison Analysis | | | |
|---|---|---|---|
| Dwass, Steel, Critchlow-Fligner Method | | | |
| Variable: TumorPrctCD8Pos | | | |
| Dx | Wilcoxon Z | DSCF Value | Pr > DSCF |
| ERPR vs. TripNeg | -0.6363 | 0.8998 | 0.8001 |
| ERPR vs. Her2 | -1.8423 | 2.6054 | 0.1559 |
| TripNeg vs. Her2 | -0.8403 | 1.1883 | 0.6779 |

| Pairwise Two-Sided Multiple Comparison Analysis | | | |
|---|---|---|---|
| Dwass, Steel, Critchlow-Fligner Method | | | |
| Variable: StromaPrctCD8Pos | | | |
| Dx | Wilcoxon Z | DSCF Value | Pr > DSCF |
| ERPR vs. TripNeg | -1.6599 | 2.3474 | 0.2208 |
| ERPR vs. Her2 | -4.3556 | 6.1598 | <.0001 |
| TripNeg vs. Her2 | -1.7739 | 2.5087 | 0.1784 |

### B - FoxP3 staining

Tumor: Distribution of Wilcoxon Score　Stroma: Distribution of Wilcoxon Score

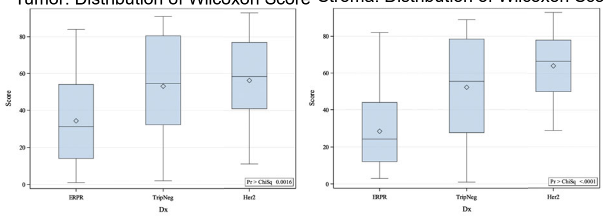


| Pairwise Two-Sided Multiple Comparison Analysis | | | |
|---|---|---|---|
| Dwass, Steel, Critchlow-Fligner Method | | | |
| Variable: TumorPrctFoxP3Pos | | | |
| Dx | Wilcoxon Z | DSCF Value | Pr > DSCF |
| ERPR vs. TripNeg | -2.6281 | 3.7167 | 0.0233 |
| ERPR vs. Her2 | -3.4082 | 4.8199 | 0.0019 |
| TripNeg vs. Her2 | -0.3268 | 0.4621 | 0.9428 |

| Pairwise Two-Sided Multiple Comparison Analysis | | | |
|---|---|---|---|
| Dwass, Steel, Critchlow-Fligner Method | | | |
| Variable: StromaPrctFoxP3Pos | | | |
| Dx | Wilcoxon Z | DSCF Value | Pr > DSCF |
| ERPR vs. TripNeg | -3.3059 | 4.6752 | 0.0027 |
| ERPR vs. Her2 | -5.4873 | 7.7602 | <.0001 |
| TripNeg vs. Her2 | -1.4471 | 2.0466 | 0.3168 |

### C - NR0B2 staining

Tumor: Distribution of Wilcoxon Score　Stroma: Distribution of Wilcoxon Score

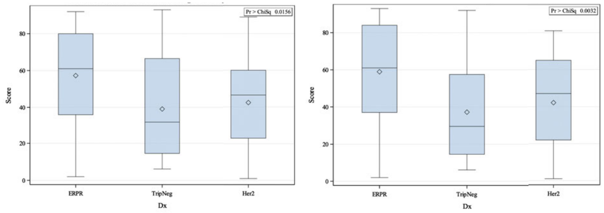


| Pairwise Two-Sided Multiple Comparison Analysis | | | |
|---|---|---|---|
| Dwass, Steel, Critchlow-Fligner Method | | | |
| Variable: AvgProbe1CopiesPerCell | | | |
| Dx | Wilcoxon Z | DSCF Value | Pr > DSCF |
| ERPR vs. TripNeg | 2.5728 | 3.6384 | 0.0273 |
| ERPR vs. Her2 | 2.2765 | 3.2195 | 0.0591 |
| TripNeg vs. Her2 | -0.5446 | 0.7702 | 0.8492 |

| Pairwise Two-Sided Multiple Comparison Analysis | | | |
|---|---|---|---|
| Dwass, Steel, Critchlow-Fligner Method | | | |
| Variable: AvgProbe1AreaPerCellum2 | | | |
| Dx | Wilcoxon Z | DSCF Value | Pr > DSCF |
| ERPR vs. TripNeg | 3.1261 | 4.4209 | 0.0050 |
| ERPR vs. Her2 | 2.5265 | 3.5731 | 0.0309 |
| TripNeg vs. Her2 | -0.7625 | 1.0783 | 0.7261 |

### D - Tumor vs. Stroma Correlations

Spearman correlation and P value for CD8 and FoxP3 cells in tumor and stroma: ER/PR (n=35)

| | Tumor % CD8+ | Stroma % CD8+ | Tumor % FoxP3+ | Stroma % FoxP3+ |
|---|---|---|---|---|
| Tumor % CD8+ | 1.00 | | | |
| Stroma % CD8+ | 0.538 (0.001) | 1.00 | | |
| Tumor % FoxP3+ | 0.146 (0.401) | 0.125 (0.476) | 1.00 | |
| Stroma % FoxP3+ | -0.192 (0.270) | 0.263 (0.127) | 0.560 (<0.001) | 1.00 |

Spearman correlation and P value for CD8 and FoxP3 cells in tumor and stroma: HER2 (n=30)

| | Tumor % CD8+ | Stroma % CD8+ | Tumor % FoxP3+ | Stroma % FoxP3+ |
|---|---|---|---|---|
| Tumor % CD8+ | 1.00 | | | |
| Stroma % CD8+ | 0.721 (<0.0001) | 1.00 | | |
| Tumor % FoxP3+ | 0.220 (0.242) | 0.177 (0.349) | 1.00 | |
| Stroma % FoxP3+ | 0.198 (0.294) | 0.263 (0.161) | 0.609 (<0.001) | 1.00 |

Spearman correlation and P value for CD8 vs. FoxP3 cells in tumor and stroma: TNBC (n=28)

| | Tumor % CD8+ | Stroma % CD8+ | Tumor % FoxP3+ | Stroma % FoxP3+ |
|---|---|---|---|---|
| Tumor % CD8+ | 1.00 | | | |
| Stroma % CD8+ | 0.885 (<0.0001) | 1.00 | | |
| Tumor % FoxP3+ | 0.705 (<0.001) | 0.668 (0.0001) | 1.00 | |
| Stroma % FoxP3+ | 0.612 (<0.001) | 0.662 (0.001) | 0.702 (<0.0001) | 1.00 |

### E - NR0B2 - FoxP3 Correlations

ALL CASES: Spearman correlation and P value of NROB2 probe metrics vs. CD8 and FoxP3 cells

| | Mean probes per tumor cell | Mean probe area per tumor cell area | Mean probes per tumor area | Percent tumor cells with >0 probes |
|---|---|---|---|---|
| Tumor % CD8+ | -0.001 (0.939) | -0.033 (0.751) | 0.039 (0.708) | -0.005 (0.965) |
| Stroma % CD8+ | -0.115 (0.272) | -0.131 (0.211) | -0.083 (0.427) | -0.114 (0.276) |
| Tumor % FoxP3+ | -0.425 (<0.0001) | -0.427 (<0.0001) | -0.371 (<0.001) | -0.405 (<0.0001) |
| Stroma % FoxP3+ | -0.353 (<0.001) | -0.365 <(0.001) | -0.344 (<0.001) | -0.333 (0.001) |

ER/PR: Spearman correlation and P value of NROB2 probe metrics vs. CD8 and FoxP3 cells

| | Mean probes per tumor cell | Mean probe area per tumor cell area | Mean probes per tumor area | Percent tumor cells with >0 probes |
|---|---|---|---|---|
| Tumor % CD8+ | -0.043 (0.81) | -0.029 (0.87) | -0.016 (0.93) | -0.049 (0.78) |
| Stroma % CD8+ | -0.020 (0.91) | 0.030 (0.86) | 0.010 (0.96) | -0.041 (0.81) |
| Tumor % FoxP3+ | -0.287 (0.09) | -0.263 (0.13) | -0.283 (0.10) | -0.258 (0.13) |
| Stroma % FoxP3+ | -0.019 (0.91) | -0.003 (0.98) | -0.006 (0.97) | -0.007 (0.97) |

HER2: Spearman correlation and P value of NROB2 probe metrics vs. CD8 and FoxP3 cells

| | Mean probes per tumor cell | Mean probe area per tumor cell area | Mean probes per tumor area | Percent tumor cells with >0 probes |
|---|---|---|---|---|
| Tumor % CD8+ | 0.252 (0.18) | 0.236 (0.21) | 0.420 (0.02) | 0.318 (0.09) |
| Stroma % CD8+ | 0.230 (0.22) | 0.225 (0.22) | 0.291 (0.12) | 0.256 (0.17) |
| Tumor % FoxP3+ | -0.441 (0.01) | -0.453 (0.01) | -0.306 (0.10) | -0.371 (0.04) |
| Stroma % FoxP3+ | -0.184 (0.33) | -0.208 (0.27) | -0.186 (0.32) | -0.079 (0.68) |

TNBC: Spearman correlation and P value of NROB2 probe metrics vs. CD8 and FoxP3 cells

| | Mean probes per tumor cell | Mean probe area per tumor cell area | Mean probes per tumor area | Percent tumor cells with >0 probes |
|---|---|---|---|---|
| Tumor % CD8+ | -0.081 (0.68) | -0.159 (0.42) | -0.070 (0.73) | -0.117 (0.55) |
| Stroma % CD8+ | -0.269 (0.17) | -0.309 (0.11) | -0.191 (0.33) | -0.280 (0.15) |
| Tumor % FoxP3+ | -0.334 (0.08) | -0.323 (0.09) | -0.258 (0.19) | -0.349 (0.07) |
| Stroma % FoxP3+ | -0.428 (0.02) | -0.477 (0.01) | -0.348 (0.07) | -0.442 (0.02) |

**Supplemental Figure 6 (SFig. 6):** Analysis and correlations between CD8, FOXP3 and NR0B2 staining in human breast tumor samples. Breast tumors were serially sectioned and stained with NR0B2 (*in situ* hybridization), FoxP3 (IHC) or CD8 (IHC). Sections were counterstained with cytokeratin to differentiate between tumoral and stromal regions. Analysis was performed collectively and for each subtype (ER/PR+, HER2+ and TNBC).

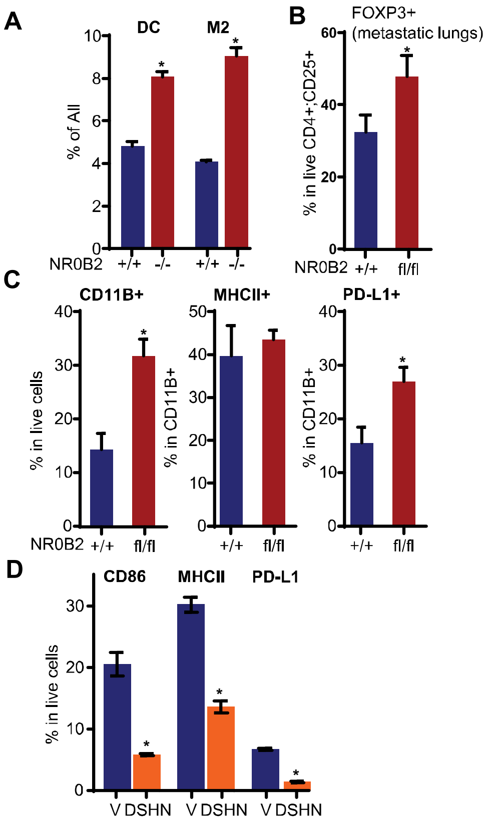


**Supplemental Figure 7 (SFig. 7): Modulation of NR0B2 results in altered immune function.** **(A)** Loss of NR0B2 in MMTV-PyMT tumors results in increased frequency of pro-inflammatory DC ($CD11b^+$;$CD11c^+$) and anti-inflammatory M2 ($CD11b^+$;$CD206^+$) -like macrophages, as determined by flow cytometry (this corresponds to **Fig. 2A-B** (N=3-6/group, t test). **(B)** $T_{regs}$ were increased in the E0771 metastatic lungs of $NR0B2^{fl/fl}$;$LysMcre^+$ mice compared to their wildtype controls. Mice were grafted i.v. with E0771 cells (N=15/group, t test). **(C)** Changes in myeloid cell populations observed in E0771 tumors grown in $NR0B2^{fl/fl}$;$LysMCre^+$ mice compared to control ($NR0B2^{fl/fl}$;$LysMCre^-$) mice (N=10-11/group, t test). **(D)** DCs ($CD11B^+$;$CD11C^+$) were isolated from MMTV-PyMT tumors and cultured in the presence of vehicle or DSHN followed by LPS and IFNγ, and subsequent populations assessed by flow cytometry (N=4/group, t test).

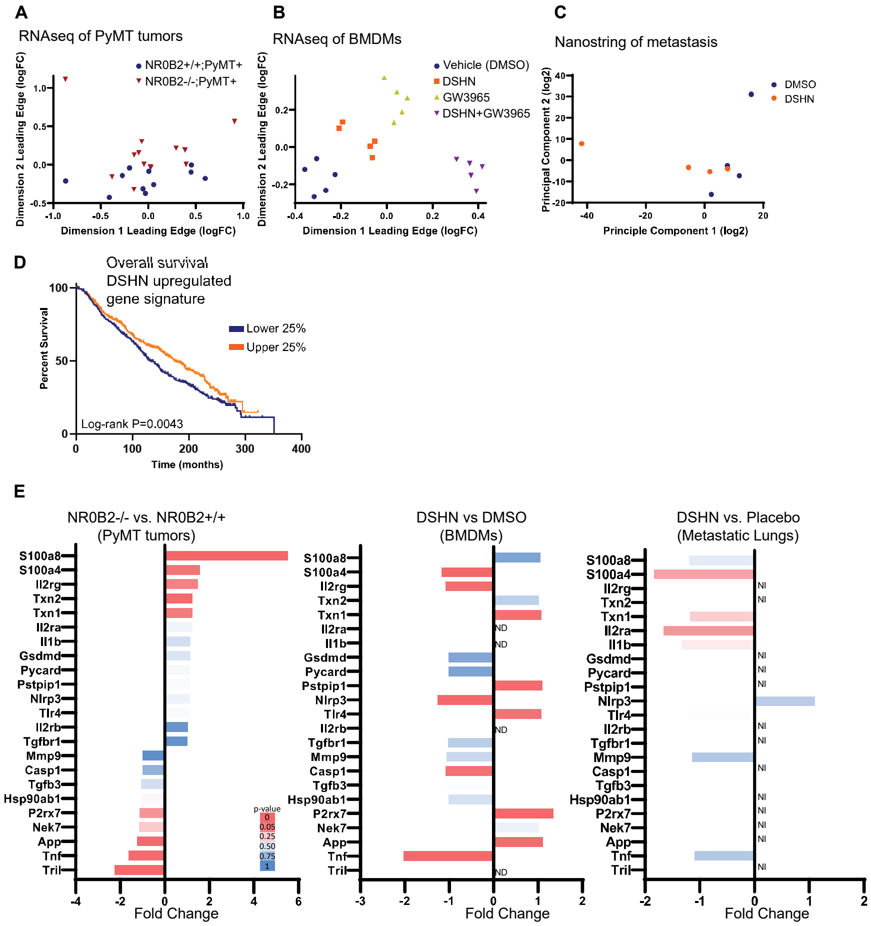


**Supplementary Figure 8 (SFig. 8): mRNA analysis indicates that inflammasome may be mediating downstream effects of myeloid cell NR0B2 on $T_{reg}$ expansion. (A)** RNAseq of MMTV-PyMT tumors from NR0B2-/- versus wildtype control mice. MDS plot shown here. **(B)** RNAseq of murine BMDMs treated with placebo (DMSO), the NR0B2 agonist DSHN, the LXR agonist GW3965 of a combination of GW3965 and DSHN. MDS plot shown here. **(C)** Nanostring analysis of isolated CD11B+;CD11C+ cells from 4T1 metastatic lungs in mice treated with either placebo or DSHN. Principal component analysis shown here, when all assessed gene expressions were input. **(D)** Signature of genes differentially upregulated genes by DSHN in BMDMs (from SFig. 8B) is associated with improved overall survival in breast cancer patients. **(E)** Fold change of select genes within the inflammasome pathway are highlighted from the three datasets. NI indicates not included for that Nanostring panel.

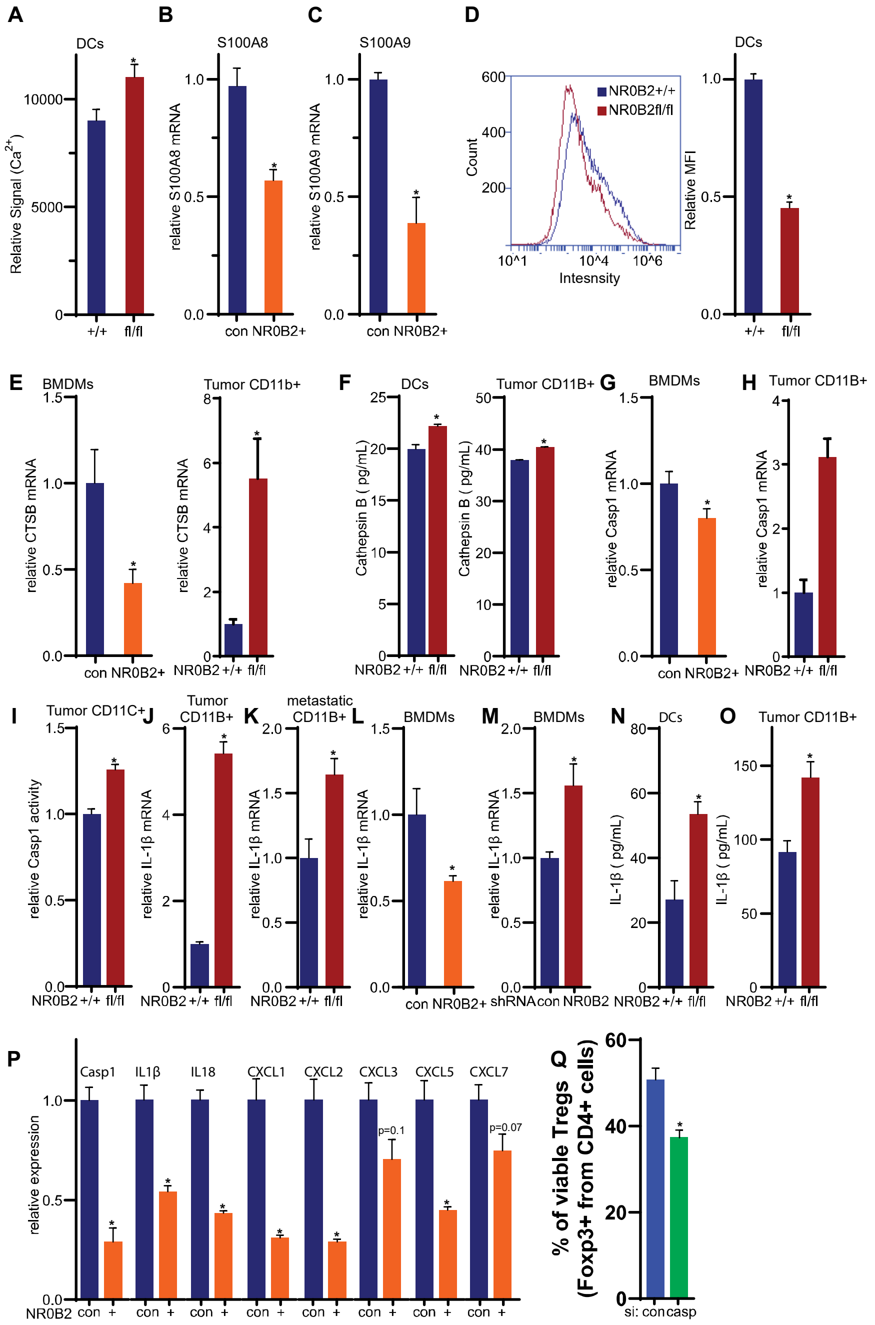


**Supplementary Figure 9 (SFig. 9): NR0B2 regulates various aspects of the inflammasome. (A)** $Ca^{2+}$ concentration in DCs treated with PMA. **(B)** S100A8 and **(C)** S100A9 mRNA in BMDMs transfected with control or NR0B2 expression plasmids, and co-cultured with E0771 cells. (**D**) Lysosome stability is decreased in DCs lacking NR0B2. Representative flow cytometry histogram to the left of quantified mean fluorescence intensity (MFI, N=4/group, t test). **(E)** Cathepsin B mRNA is downregulated when NR0B2 is overexpressed in naïve BMDMs, increased in tumor CD11B+ cells lacking NR0B2 (N=4/group, t test). **(F)** Secreted cathepsin B from DCs, or CD11b+ cells isolated from E0771 tumors, as measured by ELISA**. (G)** Caspase 1 mRNA is decreased in BMDMs overexpressing NR0B2. **(H**) Caspase 1 mRNA is increased in CD11B+ cells isolated from E0771 tumors. **(I)** Caspase 1 activity in CD11C+ cells isolated from E0771 tumors (N=4/group). **(J)** IL-1β mRNA in CD11B+ cells from primary E0771 tumors (N=4/group), or **(K)**

E0771 metastatic lungs. **(L)** IL-1β mRNA in primary BMDMs transfected with control or NR0B2 expression plasmids. **(M)** IL-1β mRNA in primary BMDMs transfected with control or shRNA plasmid against NR0B2. **(N)** IL-1β protein in media secreted from DCs. (**O**) IL-1β protein in media secreted CD11B+ cells isolated from E0771 tumors. For O and P an ELISA was performed and values below the detection limits were assigned a value of 0. **(P)** Relative expression of indicated genes associated with the inflammasome pathway. BMDMs were transfected with control pcDNA or an NR0B2 overexpression plasmid, followed by treatment with inflammasome activators (S100A8/9 + ionomycin). **(Q)** siRNA against caspase 1 in BMDMs results in decreased $T_{reg}$ expansion.

**A**

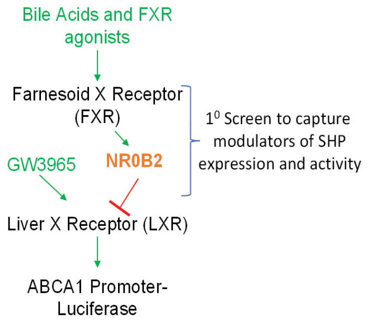


**B**

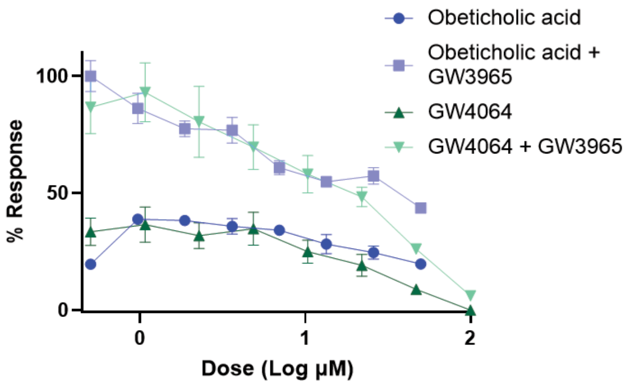


**C**

| Compound | ABCA1 reporter screen | Treg expansion macrophages | Treg expansion Dendritic cells | ABCA1 mRNA reduction (Raw Cells) | IL1b mRNA reduction (Raw Cells) | Low Toxicity |
|---|---|---|---|---|---|---|
| DSHN-OMe 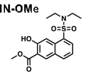 | ✓ | ✓ | ✓ | ✓ | ✓ | ✓ |
| 5 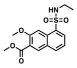 | X | ✓ | ✓ | X | ✓ | ✓ |
| 6 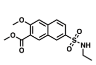 | ✓ | ✓ | ✓ | X | X | X |
| 7 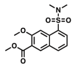 | ✓ | ✓ | ✓ | X | ✓ | ✓ |
| 8 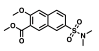 | ✓ | ✓ | ✓ | X | X | ✓ |
| 9 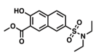 | X | ✓ | ✓ | X | ✓ | ✓ |
| 10 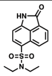 | ✓ | ✓ | ✓ | ✓ | ✓ | ✓ |

**Supplementary Figure 10 (SFig. 10): DSHN-OMe identified as NR0B2 agonist. (A)** Overview of screen used. **(B)** Screen effectively identifies that FXR ligands obetecholic acid and GW4064 reduce ABCA1-luciferase induction by LXR agonist GW3965, in a dose related manner. Data was normalized to 100 µM GW4064 (0%) and GW3965 alone (100%). **(C)** Overview of screening DSHN derivatives, using strict cutoffs with respect to DSHN for passing each metric.

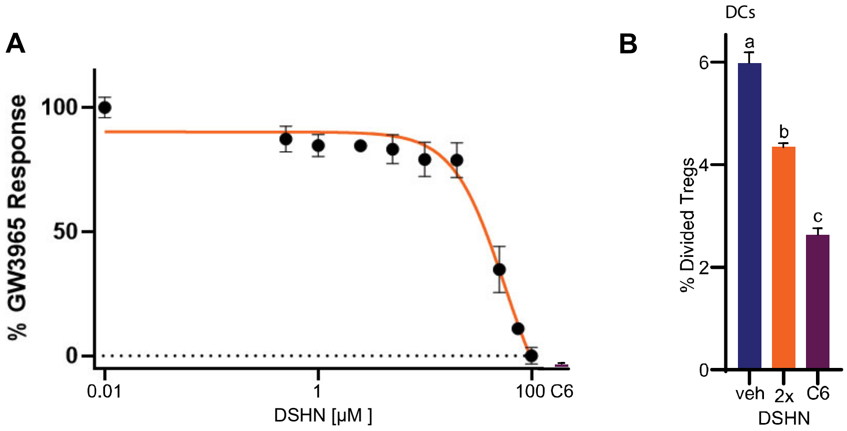


**Supplementary Figure 11 (SFig. 11): (A)** Dose response assay in BMDMs pre-treated with increasing doses of DSHN and then treated with 1μM GW3965, using ABCA1 mRNA as an endpoint. Data was normalized to 0.01μM DSHN (0%) and GW3965 alone (100%). DSHN-OMe (20μM) is indicated for comparison. Data was fit to a 4 parameter variable slope model, and $IC_{50}$ estimated at 56.2μM. **(B)** DSHN-OMe treated DCs result in decreased $T_{reg}$ expansion compared to DSHN. 50μM DSHN-OMe treated DCs primed with OVA results in decreased Treg expansion of OTI T cells compared to vehicle (DMSO) or 100μM DSHN. Different letters denote $P<0.05$.

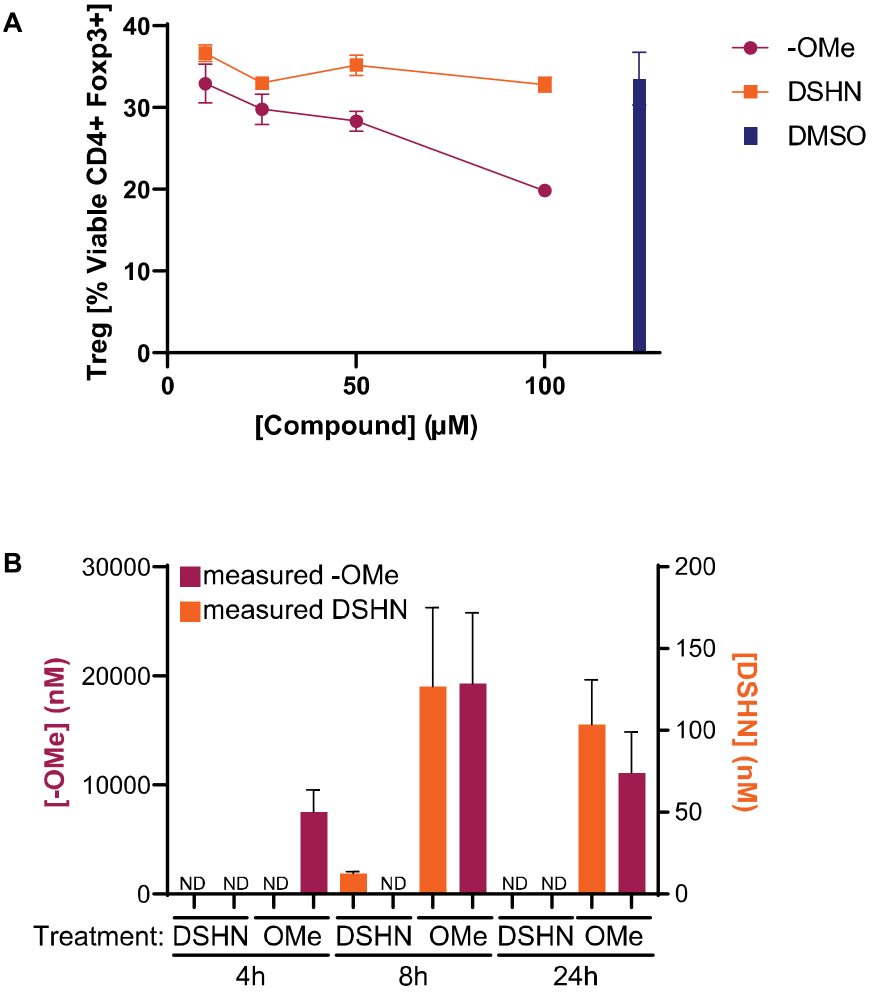


**Supplementary Figure 12: (A)** BMDMs pretreated with DSHN-OMe for 24h prior to washout and subsequent co-culture results in decreased $T_{reg}$ expansion, while DSHN does not. Vehicle (DMSO) treated BMDMs are indicated on the right for comparison. **(B)** Cellular uptake of DSHN-OMe greatly exceeds that of DSHN though time. RAW264.7 cells were treated with 50μM of vehicle (DMSO) or indicated compound for 4h, 8h or 24h. Compound was washed off, cells were lysed and resulting DSHN and DSHN-OMe contents were assessed by LC-MS/MS. Note two Y-axes. ND signifies when compounds were not detected. (N=3/group. 4h timepoint was run in an experiment independent of the 8 and 24hr timepoints).

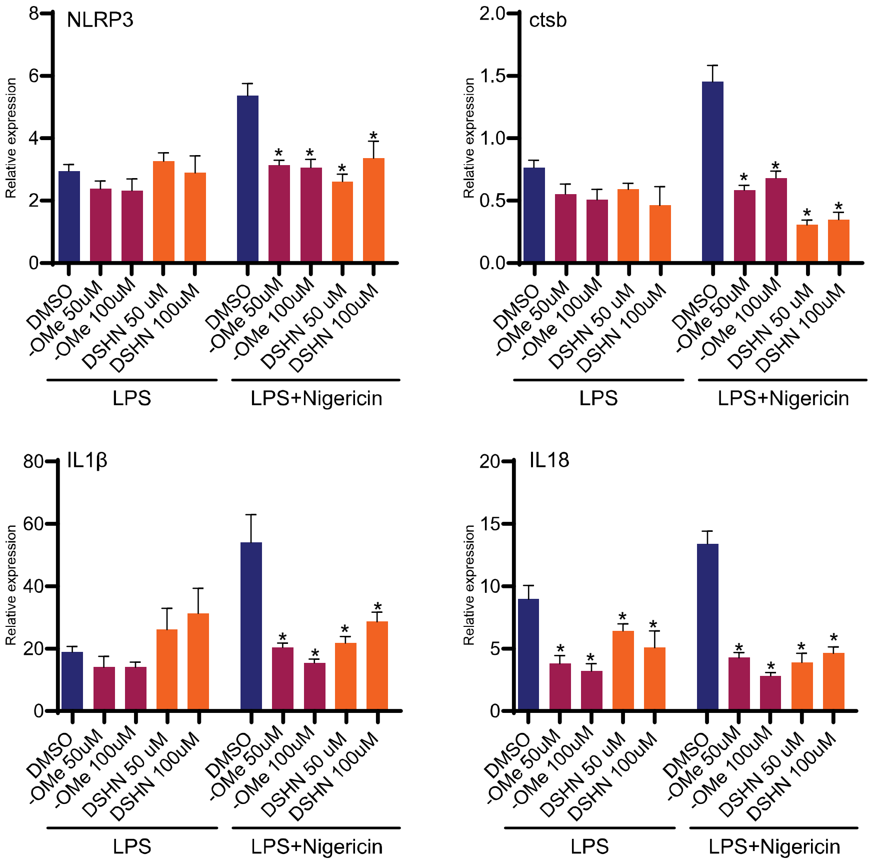


**Supplementary Figure 13**: DSHN-OMe attenuates induction of inflammasome associated genes when BMDMs are treated with both LPS and nigericin, not LPS alone. * indicates P<0.05 when comparing to DMSO.
